# Pathway redistribution across cellular states reveals a shared signaling backbone and context-dependent regulatory modules in RNA-binding protein networks

**DOI:** 10.1101/2025.03.03.641203

**Authors:** Naoki Osato, Kengo Sato

## Abstract

Understanding how regulatory architectures are reorganized during biological state transitions remains a central challenge in functional genomics. Here, we integrate co-expression–derived regulatory interactions with interpretable deep learning to compute gene-level contribution scores and introduce ΔNES (normalized enrichment score difference) to quantify pathway redistribution across biological states. Applying this framework to neural progenitor and leukemia-associated cellular states, we identify systematic redistribution of functional modules across RNA-binding proteins, including PKM, HNRNPK, and NELFE. Neural System– and Immune System–associated modules are differentially positioned along contribution-ranked regulatory landscapes, while Signal Transduction consistently forms a conserved signaling backbone. These findings suggest that pathway redistribution reflects regulatory reorganization across biological states and provides a framework for interpreting large-scale regulatory coordination beyond expression-centric analyses.

## Introduction

Understanding how regulatory architectures are reorganized across biological states remains a central challenge in molecular biology ^1–10^. Nucleic acid-binding proteins (NABPs), including DNA-binding proteins (DBPs) and RNA-binding proteins (RBPs), play key roles in gene regulation; however, their functional characterization across diverse cell types remains incomplete. While experimental approaches such as ChIP-seq and eCLIP can identify binding sites, these methods are not easily scalable and often fail to capture indirect or context-dependent regulatory interactions^11^, particularly for RBPs lacking well-defined binding motifs^12–14^.

Computational approaches, including deep learning models, have improved the prediction of gene expression by integrating regulatory features derived from binding data. However, these methods typically rely on experimentally defined binding sites or sequence features and do not directly reveal how individual NABPs contribute to gene expression programs at the systems level. In particular, it remains unclear how regulatory influence is organized across pathways and how it is reconfigured between cellular contexts.

Gene co-expression analysis provides a complementary perspective by capturing coordinated expression patterns across diverse biological conditions, reflecting both direct and indirect regulatory relationships^15–23^. Integrating such information into interpretable deep learning frameworks offers the potential to quantify regulatory influence at the gene level without requiring prior knowledge of binding motifs or direct interaction data.

Here, we integrate co-expression–derived regulatory interactions with interpretable deep learning to compute gene-level contribution scores and introduce ΔNES (normalized enrichment score difference) to quantify pathway redistribution between cellular contexts. Rather than focusing on individual regulatory interactions, this framework enables the analysis of regulatory architecture at the pathway level.

Applying this approach across distinct biological states represented by neural progenitor and leukemia-associated systems, we find that regulatory influence is not organized as distinct pathway sets, but instead appears to be structured around a shared signaling backbone, while functional modules are redistributed in a context-dependent manner. These findings suggest a framework for interpreting regulatory architecture, offering a systems-level perspective on gene regulation across cellular states.

## Results

### Gene co-expression features enhance expression prediction accuracy

Replacing low-contribution DBP inputs with gene co-expression data led to a substantial improvement in model performance (**Fig. 1**). In human foreskin fibroblasts (HFFs), substituting 302 out of 1,310 DBPs with co-expression profiles increased the correlation between predicted and observed gene expression from 0.70 to 0.80 (**Fig. 2a, b**). These co-expression inputs included both NABP-encoding and non-NABP mRNAs, with gene symbols for DNA- and RNA-binding proteins obtained from the ENPD (Eukaryotic Nucleic acid Binding Protein) database^24^ and co-expression profiles derived from COXPRESdb^15^. A subsequent experiment replaced 325 low-contribution DBPs with co-expression interactions involving RBPs further improved prediction accuracy (*r* = 0.81; **Fig. 2c**). However, excessive replacement of input features with gene co-expression data reduced the biological interpretability of model predictions, suggesting that overreliance on co-expression signals may compromise the model’s capacity to learn gene expression patterns from primary regulatory features. Collectively, these findings demonstrated that the selective integration of co-expression interactions can enhance predictive performance while preserving biologically meaningful regulatory inference beyond direct binding site information.

**Figure 1.**
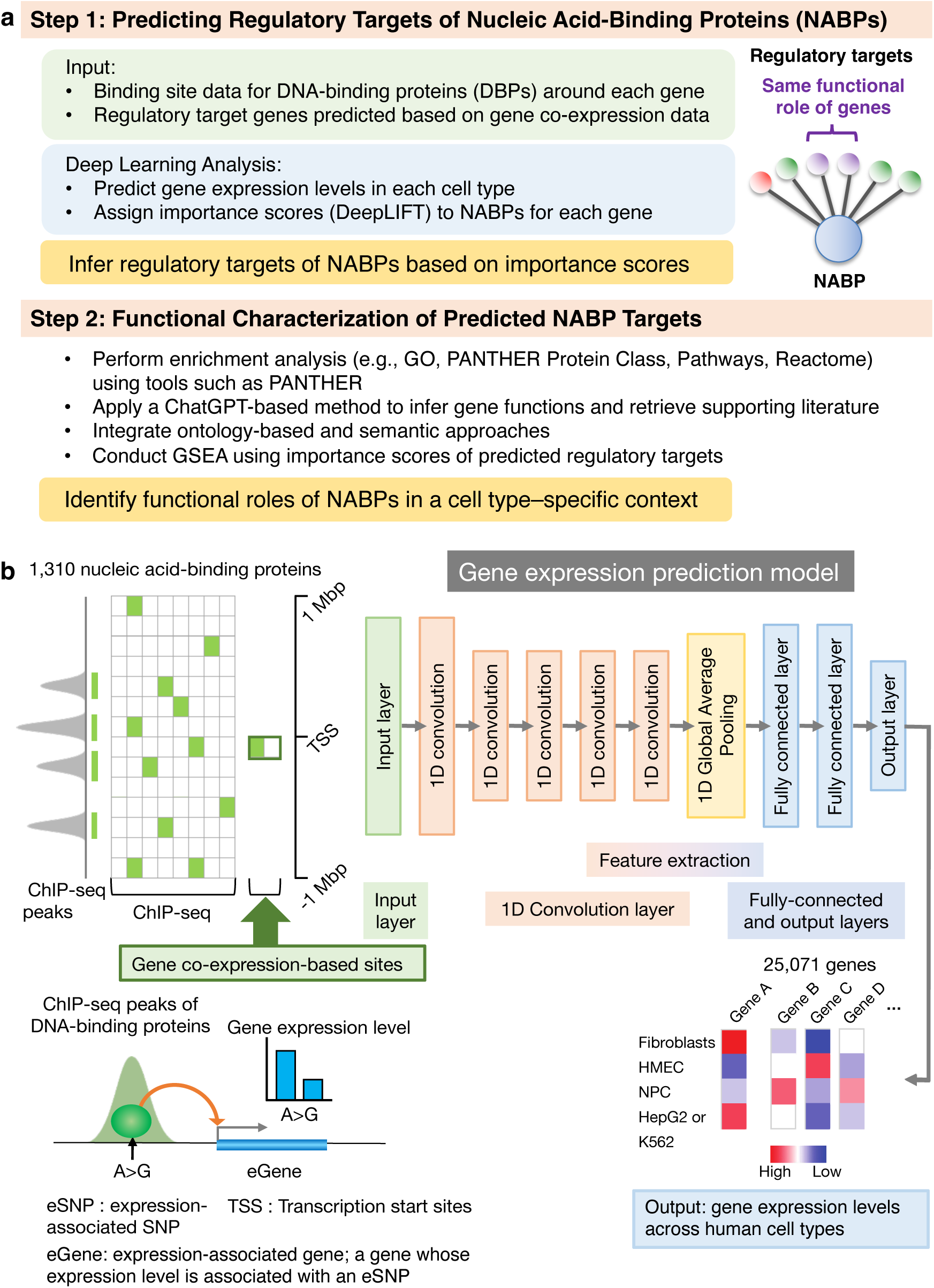
Integrated framework for predicting regulatory targets and dissecting functional organization of NABPs across cellular contexts. **a. Schematic overview of the analytical framework.** Gene expression profiles from multiple human cell types were integrated with NABP–gene co-expression relationships derived from COXPRESdb. These inferred regulatory associations were used to replace low-contribution DNA-binding protein features in a deep learning model originally trained on ChIP-seq–based binding site data. The modified model was trained to predict gene expression and to infer regulatory target genes using DeepLIFT-derived contribution scores. Functional characterization was performed through a three-layer analytical strategy: (i) statistical enrichment analysis using PANTHER to identify significantly associated biological processes, (ii) large language model–assisted interpretation to contextualize and integrate functional themes, and (iii) contribution score–based GSEA to quantify pathway redistribution across cellular contexts. **b. Gene expression prediction framework based on genomic distributions of DNA-binding sites across promoter-proximal and distal regulatory regions.** During model training, DNA-binding proteins associated with distal regulatory elements overlapping expression-associated SNPs (eSNPs) were linked to candidate target genes (eGenes) using cis-eQTL information. For each gene–protein pair, ChIP-seq peak distributions were binned (green boxes) and assembled into a unified feature matrix. In co-expression–based models, inferred regulatory interactions were represented as hypothetical binding sites near transcription start sites (TSSs). The deep learning architecture comprised one-dimensional convolutional layers, global average pooling, and fully connected layers, enabling prediction of gene expression across 25,071 genes in multiple human cell types.

**Figure 2.**
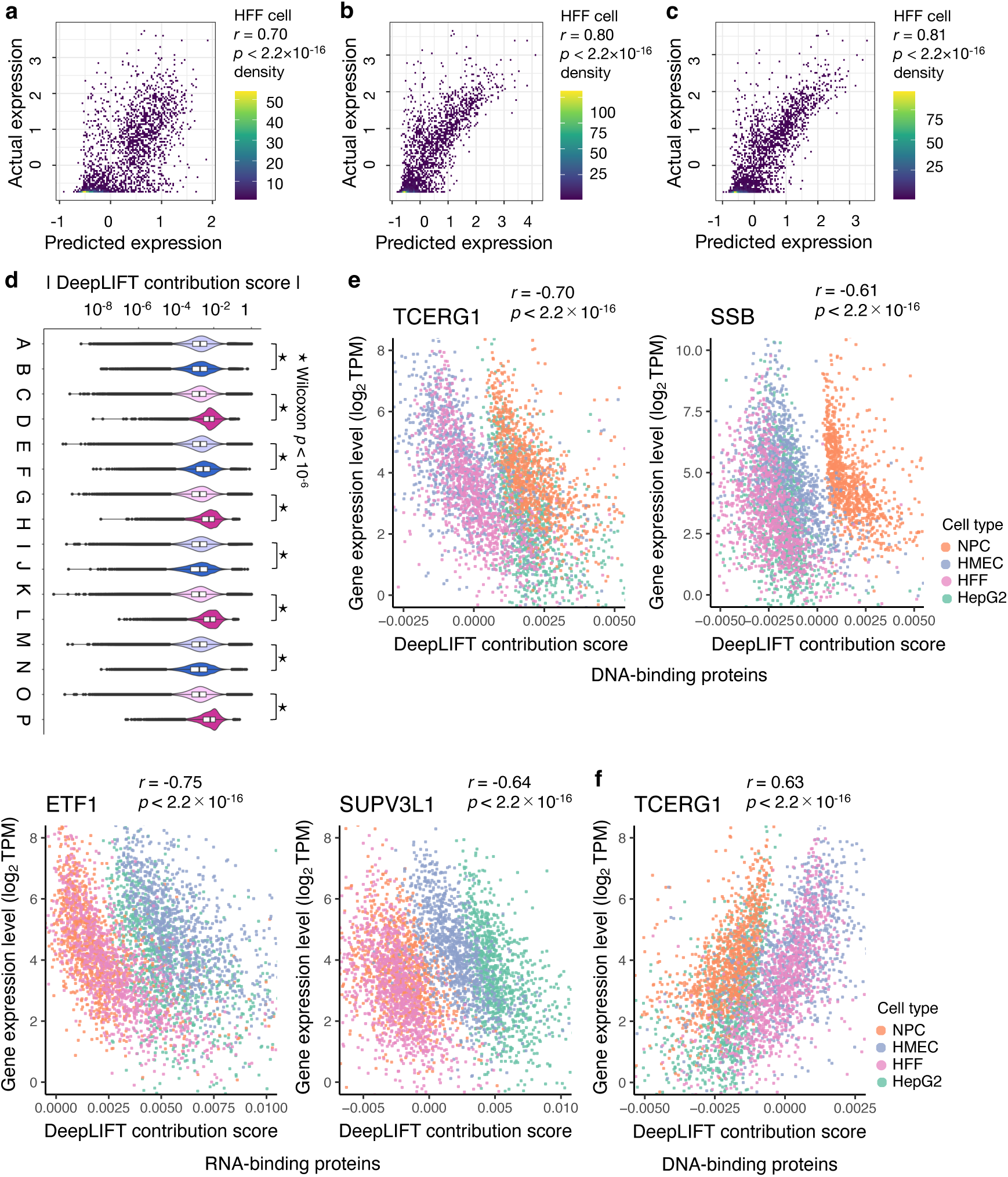
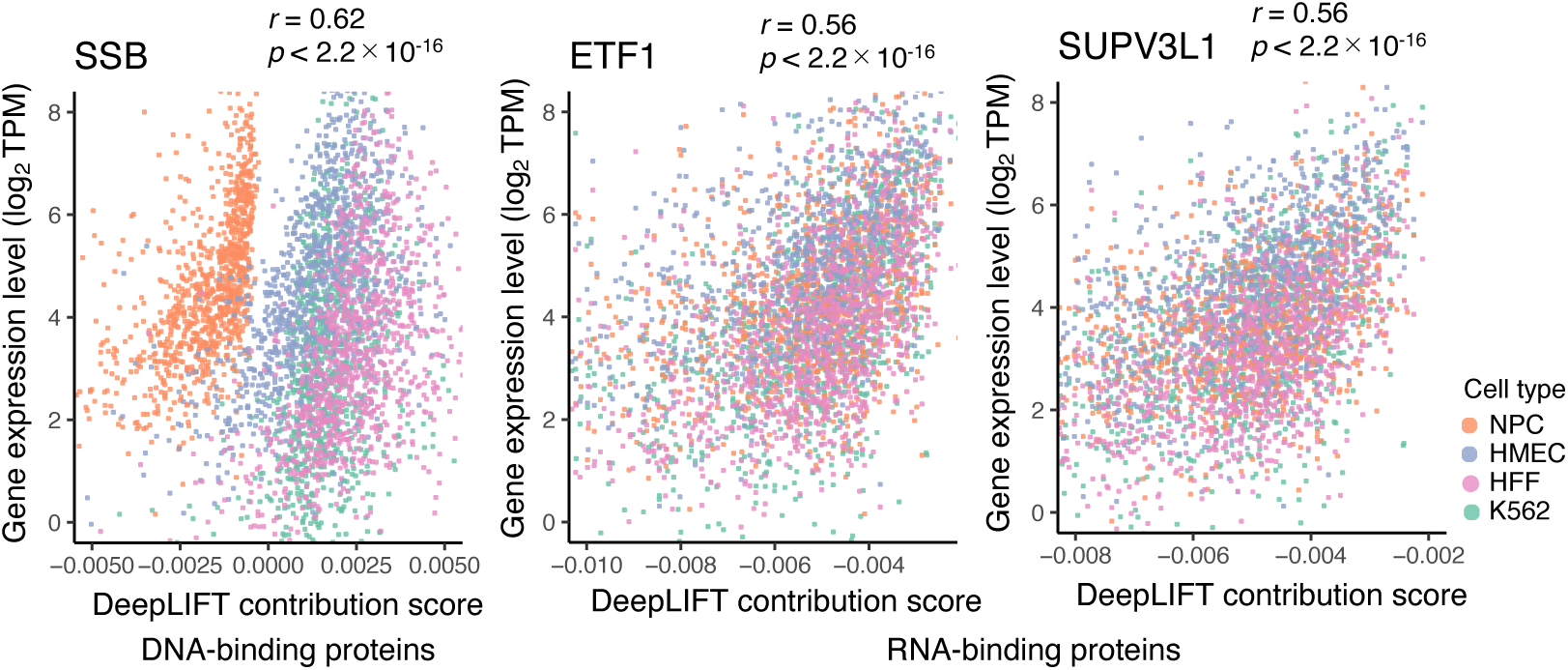
Relationship between gene expression levels and DeepLIFT contribution scores. Predicted gene expression levels generated by the deep learning model were compared with observed expression levels using Spearman’s rank correlation. **a–c. Schematic overview of input feature configurations used for gene expression prediction.** **a.** Baseline model using DNA-binding site features from 1,310 DNA-binding proteins derived from ChIP-seq data. **b.** Co-expression–based model in which 302 low-contribution DNA-binding proteins were replaced with alternative DNA-binding proteins inferred from gene co-expression relationships. **c.** RNA-binding protein–augmented model in which 325 low-contribution DNA-binding protein features were replaced with RNA-binding proteins identified through co-expression–based regulatory inference. **d. Comparison of absolute DeepLIFT contribution scores between predicted regulatory sites derived from gene co-expression and experimentally defined ChIP-seq binding sites, using a reduced DeepLIFT background reference (input × 0.5).** Each pair of violin plots represents co-expression–derived regulatory sites (right) and the corresponding excluded ChIP-seq sites (left). Across all examined conditions, contribution score magnitudes were significantly higher for co-expression–derived regulatory sites (Wilcoxon signed-rank test, *p*-value < 10⁻⁶). Equivalent analyses using an amplified background reference (input × 2) are shown in **Supplementary Figure S2c**. **e,f. Relationship between observed gene expression levels and DeepLIFT contribution scores for representative DNA-binding proteins and RNA-binding proteins across four human cell types.** Contribution scores were calculated using either a reduced background reference (input × 0.5; **e**) or an amplified background reference (input × 2; **f**). The same representative proteins are shown in both panels to enable direct comparison of background-dependent effects. Although background scaling altered the polarity of DeepLIFT–expression correlations, relative contribution score rankings and cell type–specific distributional structures were consistently preserved. For each protein, the displayed Spearman correlation coefficient and associated *p*-value correspond to the cell type exhibiting the strongest correlation among the four analyzed cell types. Data points are color-coded by cell type, highlighting reproducible cell type–specific regulatory patterns across background conditions.

Contribution scores derived from co-expression–based NABP–gene interactions exhibited broader distributions and higher overall magnitudes than those obtained from ChIP-seq–based inputs across all four examined cell types (Wilcoxon signed-rank test, *p* < 10⁻^2^; **Fig. 2d**, **Supplementary Fig. S2c,** and **Supplementary Table S1**). Each pair of violin plots represents the absolute contribution scores of co-expression–based regulatory sites (right plot in each pair) and the corresponding ChIP-seq-derived sites excluded from that co-expression set (left plot). **A**–**P** denote specific cell types and NABP classes as follows: **B**, DBP co-expression–based sites in neural progenitor cells (NPCs); **A**, NPC ChIP-seq sites excluding **B**. **D**, RBP co-expression–based sites in NPCs; **C**, NPC ChIP-seq sites excluding **D**. **F**, DBP co-expression–based sites in HepG2 cells; **E**, HepG2 ChIP-seq sites excluding **F**. **H**, RBP co-expression–based sites in K562 cells; **G**, K562 ChIP-seq sites excluding **H**. **J**, DBP co-expression–based sites in HMEC cells; **I,** HMEC ChIP-seq sites excluding **J**. **L**, RBP co-expression–based sites in HMEC cells; **K**, HMEC ChIP-seq sites excluding **L**. **N**, DBP co-expression–based sites in human foreskin fibroblasts (HFFs); **M**, HFF ChIP-seq sites excluding **N**. **P**, RBP co-expression–based sites in HFFs; **O**, HFF ChIP-seq sites excluding **P**. Corresponding violin plots generated using an amplified DeepLIFT background reference (input × 2) are shown in **Supplementary Figure S2c**. However, the nature of these differences varied by nucleic acid–binding protein class. For DNA-binding proteins (DBPs), contribution score distributions differed significantly between ChIP-seq–derived and co-expression–derived regulatory sites, although their mean and median values were often similar across cell types. This suggests that, for DBPs, co-expression–based feature selection primarily reshapes the distributional architecture of contribution scores without consistently altering their central tendency. In contrast, for RNA-binding proteins (RBPs), regulatory sites inferred from gene co-expression consistently exhibited substantially higher DeepLIFT contribution scores than those derived solely from experimental binding data. This pattern was observed across all examined cell types, including K562, NPCs, HMECs, and HFFs, and was reflected in pronounced increases in both mean and median contribution scores. These findings suggest that co-expression–based inference preferentially enriches for RBP-associated regulatory interactions that are more directly coupled to gene expression prediction, whereas experimental identified RBP binding sites may include a larger proportion of interactions with limited transcriptomic impact. These observations are consistent with previous reports demonstrating that the incorporation of RNA-binding protein– and miRNA-associated regulatory features can improve gene expression prediction accuracy, as exemplified by the DEcode framework^25^. Importantly, these trends were robust across DeepLIFT background definitions: comparable distinctions between ChIP-seq– and co-expression–derived regulatory targets were observed under both background scaling schemes (input × 0.5 and input × 2). Thus, although background scaling influences the sign and absolute magnitude of contribution scores, it does not alter the fundamental distinction between binding-based and co-expression–based regulatory inference.

To assess the influence of background selection on DeepLIFT-derived contribution scores, we compared results obtained using a reduced background reference (input × 0.5; **Fig. 2e**) and an amplified background reference (input × 2; **Fig. 2f**) for the same representative DNA- and RNA-binding proteins. Under the reduced background, contribution scores generally exhibited negative correlations with gene expression, whereas the amplified background condition produced predominantly positive correlations. These patterns were approximately mirrored around zero, consistent with DeepLIFT’s definition as a relative contribution metric defined with respect to the selected reference baseline. Importantly, despite this inversion in sign, the relative ranking of genes and the overall cell type–specific structure of contribution score distributions remained highly consistent across background settings. In both configurations, genes with larger-magnitude contribution scores were preferentially associated with stronger expression-related effects, indicating that DeepLIFT captures relative regulatory importance rather than absolute activation or repression. A conceptual model explaining how input definition and background scaling generate these patterns is provided in **Supplementary Figure S4**. Notably, background-dependent differences were more pronounced for RNA-binding proteins (RBPs) (**Fig. 2e,f**). For the same representative RBPs, scatter plots generated using the reduced background (input × 0.5) displayed more distinct and partially separated distributions across the four cell types, whereas under the amplified background (input × 2), data points from different cell types showed substantially greater overlap. This suggests that the reduced-background configuration enhances sensitivity to cell type–specific regulatory variation in RBP-associated contribution profile. In contrast, DNA-binding proteins (DBPs) generally exhibited stronger correlations with gene expression under both background conditions, consistent with their more direct roles in transcriptional regulation. Together, these findings underscore the importance of background-aware and protein class–specific interpretation when analyzing DeepLIFT contribution scores.

### Experimental binding data support model-inferred regulatory targets

To determine whether DeepLIFT-derived regulatory contribution scores reflect biologically meaningful nucleic acid–binding protein (NABP) activity, we systematically compared model-inferred regulatory targets with experimentally validated ChIP-seq and eCLIP binding data across multiple genomic regions, threshold definitions, and cellular contexts (**Fig. 3**). Experimentally validated binding data were broadly consistent with deep learning–derived regulatory targets. Genes containing ChIP-seq or eCLIP peaks for a given NABP disproportionately occupied high- or low-importance ranks, indicating that experimentally confirmed binding sites are associated with stronger regulatory influence in gene expression prediction models. This relationship supports the biological validity of DeepLIFT-based contribution scores, as quantitative measures of NABP-specific regulatory importance.

**Figure 3.**
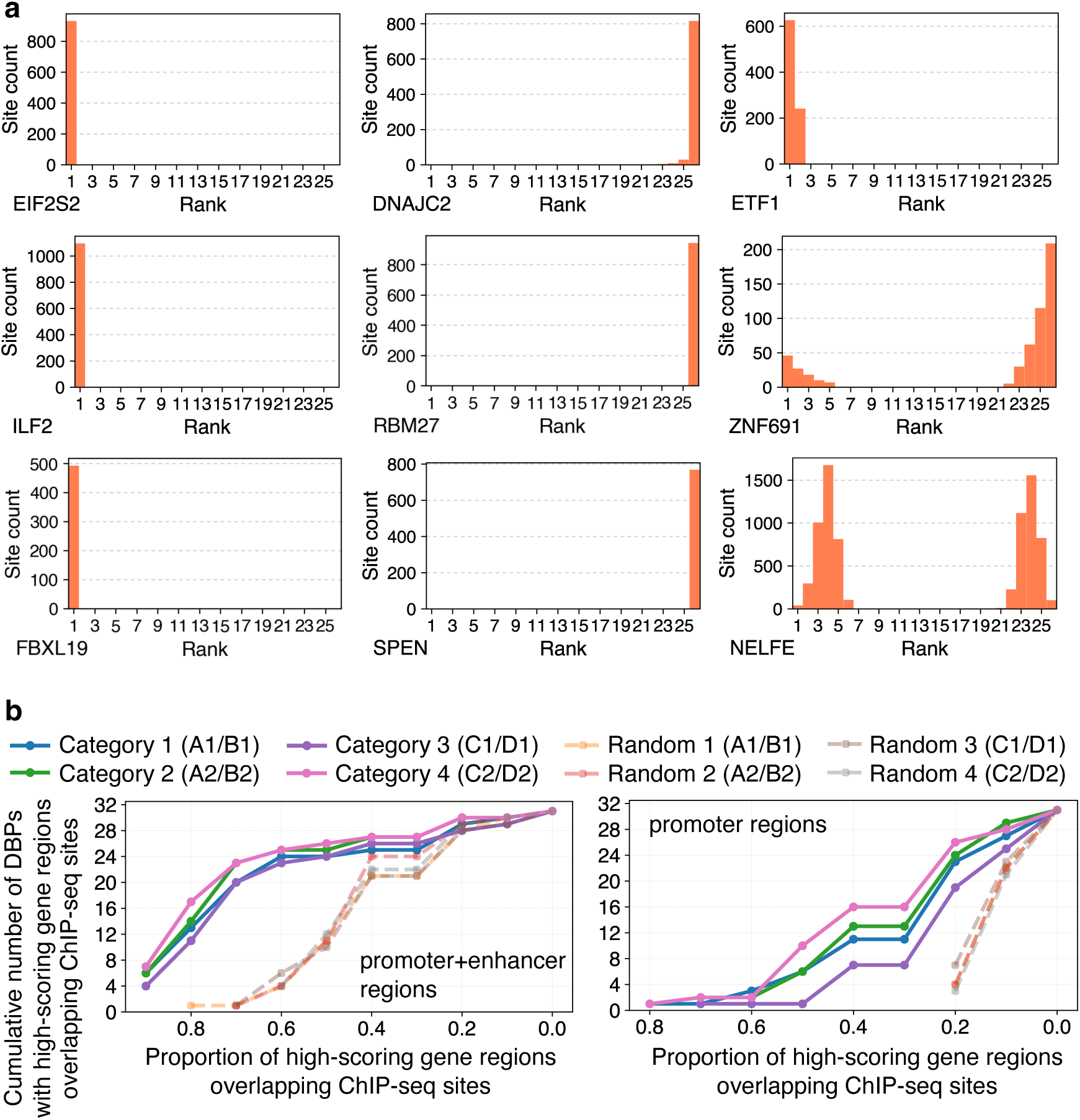
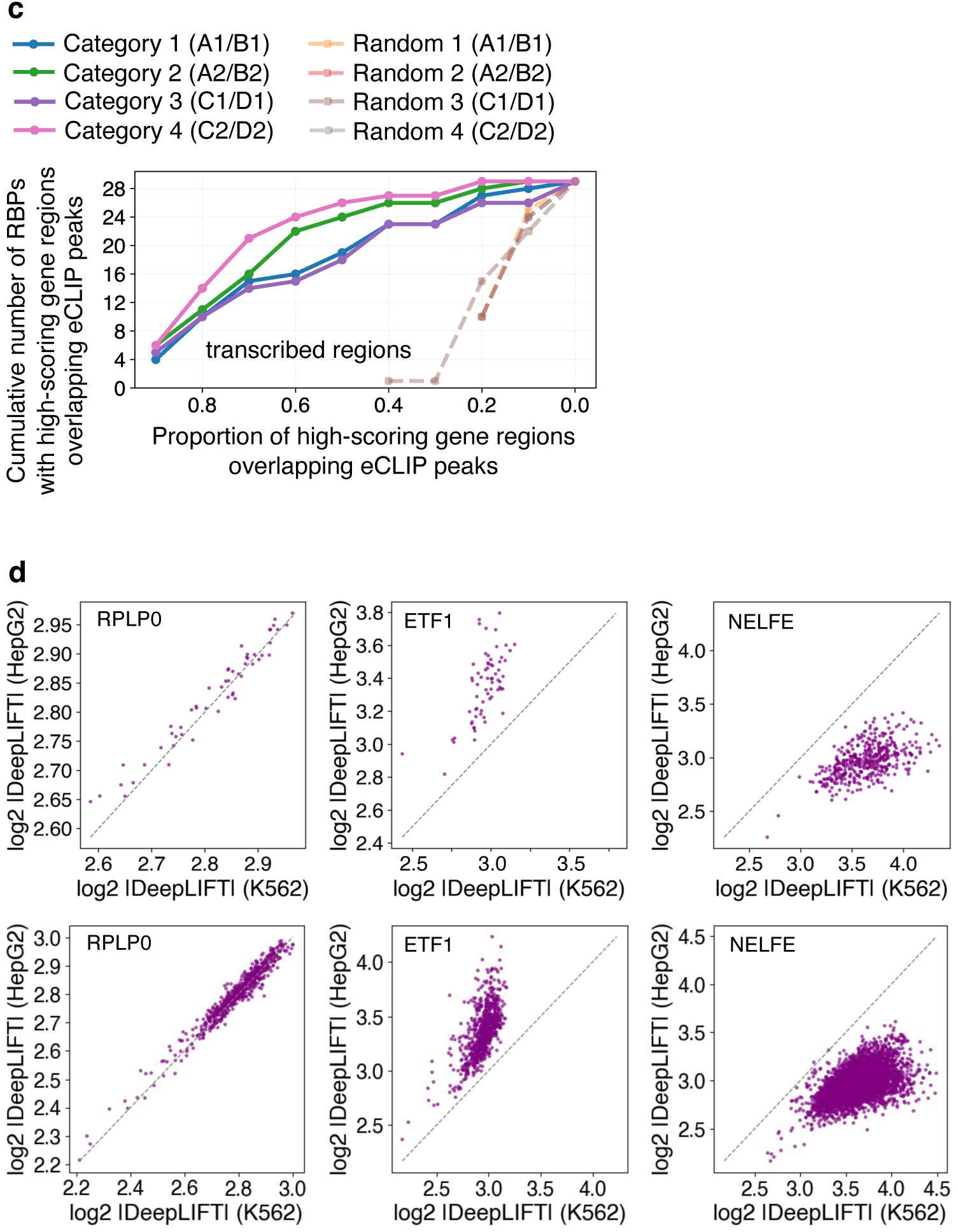
Comparison of predicted regulatory target genes with experimental ChIP-seq and eCLIP binding data. This figure illustrates the correspondence between gene-level regulatory contribution scores inferred by deep learning and experimentally validated nucleic acid–binding protein (NABP) binding sites, using DeepLIFT with a reduced background reference (input × 0.5). **a. Rank distribution of DeepLIFT contribution scores for experimentally validated NABP binding sites identified by ChIP-seq (DNA-binding proteins) or eCLIP (RNA-binding proteins).** Experimentally validated binding sites located within promoter or transcribed regions exhibited strong non-random enrichment toward extreme contribution score ranks (both high and low), indicating preferential association with genes predicted to have strong regulatory influence. **b. Proportion of high-contribution genomic regions overlapping ChIP-seq binding sites for DNA-binding proteins.** Cumulative distributions show overlap within promoter + enhancer regions (±100 kb from transcription start sites) and promoter-proximal regions alone (±5 kb). Multiple rank- and score-based thresholds were used to define high-contribution regions (see **Supplementary Note** and **Supplementary Tables S2 and S3**). Across all thresholds, overlap exceeded random expectations. **c. Overlap between high-contribution regulatory regions and eCLIP peaks for RNA-binding proteins.** Cumulative distributions show the proportion of contribution-ranked transcribed regions overlapping eCLIP peaks across gene bodies, compared with matched random controls. Across all evaluated thresholds, overlap with eCLIP peaks consistently exceeded matched random expectations. Similar patterns under amplified background conditions (input × 2) are shown in **Supplementary Figure S3**. **d. Comparison of DeepLIFT contribution score distributions between HepG2 and K562 cells under alternative background definitions.** Scatter plots show experimentally validated target genes of representative NABPs (RPLP0, ETF1, and NELFE), comparing K562 (x-axis) and HepG2 (y-axis). Absolute contribution scores were log₂-transformed. Reduced and amplified background configurations preserved relative regulatory structure despite polarity changes. RPLP0 showed conserved distributions across cell types, whereas ETF1 and NELFE exhibited stronger cell type–dependent divergence.

To evaluate this association, we analyzed ChIP-seq data for 32 DBPs in HepG2 cells and eCLIP data for 29 RBPs in K562 cells—two cell lines with the most comprehensive publicly available NABP binding datasets from ENCODE and related studies. A gene was considered associated with an NABP if a ChIP-seq peak was located within ±5 kb of its transcription start site (TSS) or if an eCLIP peak was mapped to its transcribed region. As illustrated in **Fig. 3a** and **Supplementary Fig. S1**, NABPs with experimentally validated binding sites consistently exhibited higher absolute contribution score magnitudes across predicted target genes, reinforcing their inferred regulatory relevance and aligning with prior interpretations of DeepLIFT scores^25,26^.

Model-derived importance scores also reliably predicted DBP-associated regulatory targets, as evidenced by significant overlap with ChIP-seq peaks (**Fig. 3b**; **Supplementary Table S2**, and **S3**; **Supplementary Fig. S3a**, and **Supplementary Note and Tables section: Gene co-expression-based predictions align with ChIP-seq-defined DNA-binding targets**). To quantify this correspondence, we analyzed ChIP-seq data for 32 DBPs and assessed how frequently high-scoring gene regions—defined by DeepLIFT contribution scores—coincided with experimental binding evidence. Overlap ratios were calculated as the proportion of high-contribution regions intersecting ChIP-seq peaks, considering both promoter-proximal (±5 kb) and distal (±100 kb) regions relative to TSSs. To account for the functional heterogeneity and uneven abundance of ChIP-seq peaks, we applied a precision-at-K framework, prioritizing top-ranked or top-scoring gene regions most likely to represent biologically meaningful regulatory interactions. Four thresholds were used to define high-contribution regions: two based on contribution score rank (top/bottom 20 or 50 genes) and two based on absolute score magnitude (≥ 0.01 or ≤ –0.01; ≥ 0.001 or ≤ –0.001). A matched random control analysis using gene regions of equivalent size was performed in parallel. Across all tested conditions, overlap between ChIP-seq peaks and high-contribution regions consistently exceeded random expectations, supporting the model’s capacity to prioritize functionally relevant regulatory sites.

Similarly, as illustrated in **Fig. 3c**, **Supplementary Table S4**, **Supplementary Fig. S3b**, and the **Supplementary Note and Tables section: Gene co-expression-based predictions align with ChIP-seq-defined DNA-binding targets**, model-derived importance scores consistently prioritize biologically relevant RBP–target interactions. This was evidenced by the cumulative overlap between high-contribution gene regions and eCLIP peaks, which was substantially greater than that observed in matched random controls. To evaluate this relationship, we analyzed eCLIP data for 29 RBPs in K562 cells, comparing high-contribution regions—defined using the same thresholds applied for DBPs—to experimentally identified eCLIP peaks mapped across transcribed gene regions (from transcription start site [TSS] to transcription end site [TES]). Collectively, these findings demonstrate that the model effectively captures functionally meaningful NABP–target associations for both DNA- and RNA-binding proteins.

Significant differences in predicted regulatory target distributions were observed for several NABPs between K562 and HepG2 cells based on DeepLIFT contribution scores (**Fig. 3d**). To assess cell type–specific regulatory activity, we compared DeepLIFT contribution score distributions for genes harboring experimentally validated NABP binding sites identified by ChIP-seq or eCLIP analyses. Because the deep learning framework identifies target candidates from NABP-associated co-expressed gene sets, predicted targets do not necessarily represent direct physical binding alone, but may also reflect genes participating in shared regulatory programs or downstream functional networks. For example, RPLP0 exhibited nearly identical contribution score distributions across the two cell types, whereas ETF1 and NELFE displayed pronounced divergence. These differences indicate that the model captures context-dependent NABP regulatory relationships that extend beyond static binding information alone.

### Deep learning uncovers functionally enriched NABP target genes beyond co-expression networks

Deep learning predictions revealed that nucleic acid–binding proteins (NABPs) are associated with regulatory target gene sets exhibiting coherent and biologically specific functional enrichment, extending substantially beyond patterns detectable from co-expression alone (**Tables 1,2** and **Supplementary Note 2**). Across twelve representative NABPs^27,28^, predicted target sets consistently showed significant enrichment for biologically relevant Gene Ontology (GO) terms, whereas randomly sampled gene sets of matched size exhibited no comparable enrichment (**Fig. 4**). Notably, several enriched functional annotations were absent from the broader underlying co-expressed gene pools (Pearson’s *z*-score > 2), indicating that the model selectively extracts biologically meaningful regulatory signals from large co-expression neighborhoods rather than merely recapitulating preexisting co-expression structure. Functional enrichment analyses across GO Slim, Reactome, and PANTHER Pathways further reinforced this specificity (**Table 1**)^29^. An independent large language model–based gene set interpretation approach further supported these findings by generating functional summaries and literature associations consistent with GO enrichment results and UniProt annotations^30^ (**Table 2** and **Supplementary Note 2**)^31^.

**Figure 4.**
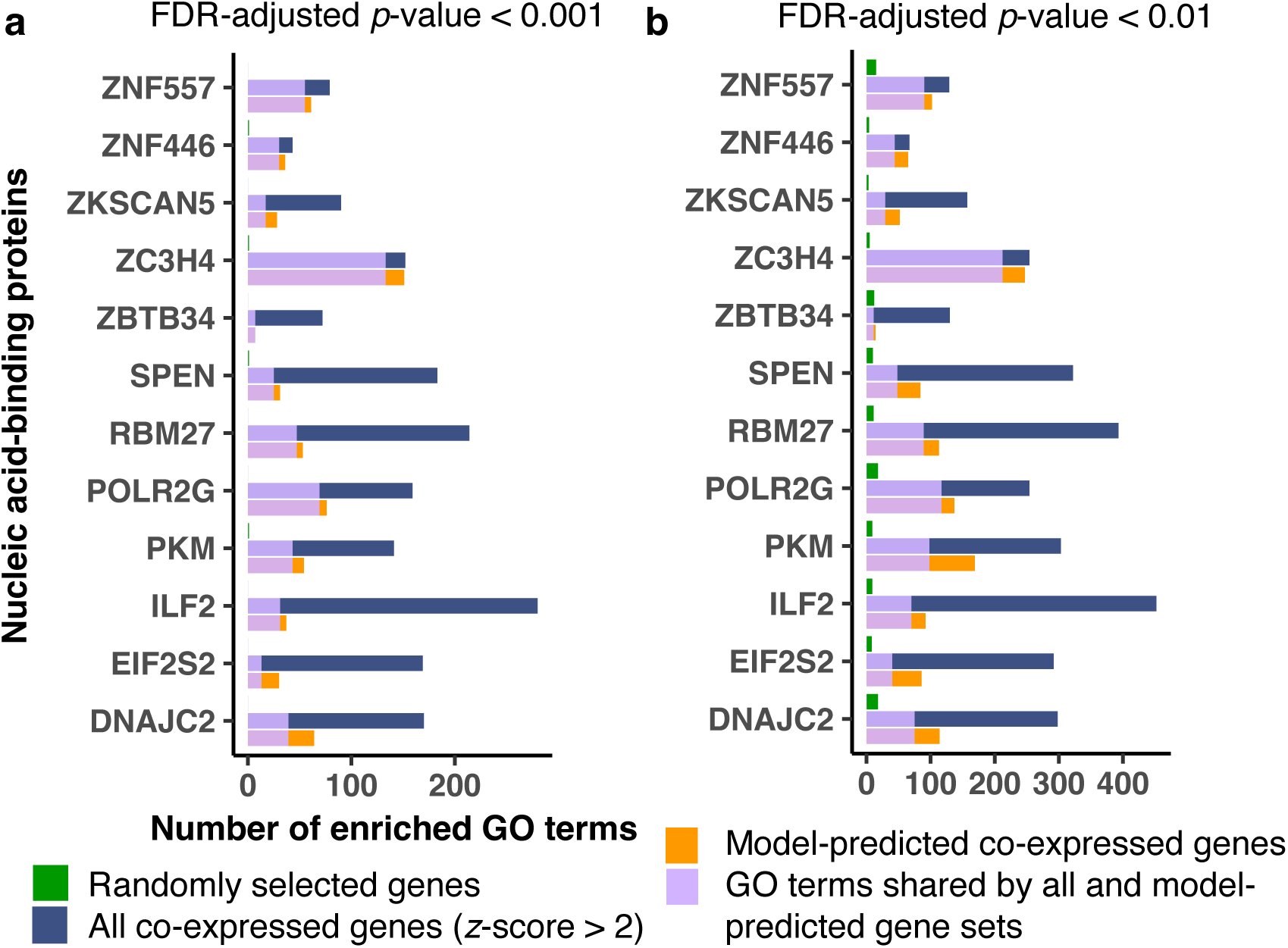
Overrepresented functional annotation terms among predicted regulatory target genes of nucleic acid-binding proteins (NABPs). **a,b. Functional enrichment analysis of predicted NABP regulatory target genes compared with broader co-expression networks and matched random controls.** Panels **a** and **b** display significantly enriched functional annotation terms filtered at false discovery rate (FDR) thresholds of < 0.001 and < 0.01, respectively. For each NABP, functional enrichment was evaluated across three gene sets: (1) deep learning model–predicted primary regulatory target genes derived from co-expression–refined NABP interactions, (2) all co-expressed genes associated with the NABP (Pearson’s z-score > 2), and (3) randomly selected gene sets of matched size. Gene Ontology (GO) terms spanning biological process, molecular function, and cellular component categories were combined to assess overall biological coherence. Enrichment significance was calculated using Fisher’s exact test with Benjamini–Hochberg correction for multiple hypothesis testing. Across multiple NABPs, predicted regulatory target gene sets consistently exhibited substantially greater functional enrichment specificity than broader co-expression networks or random controls, supporting the capacity of deep learning–based refinement to prioritize biologically coherent regulatory targets beyond co-expression structure alone.

**Table 1.**
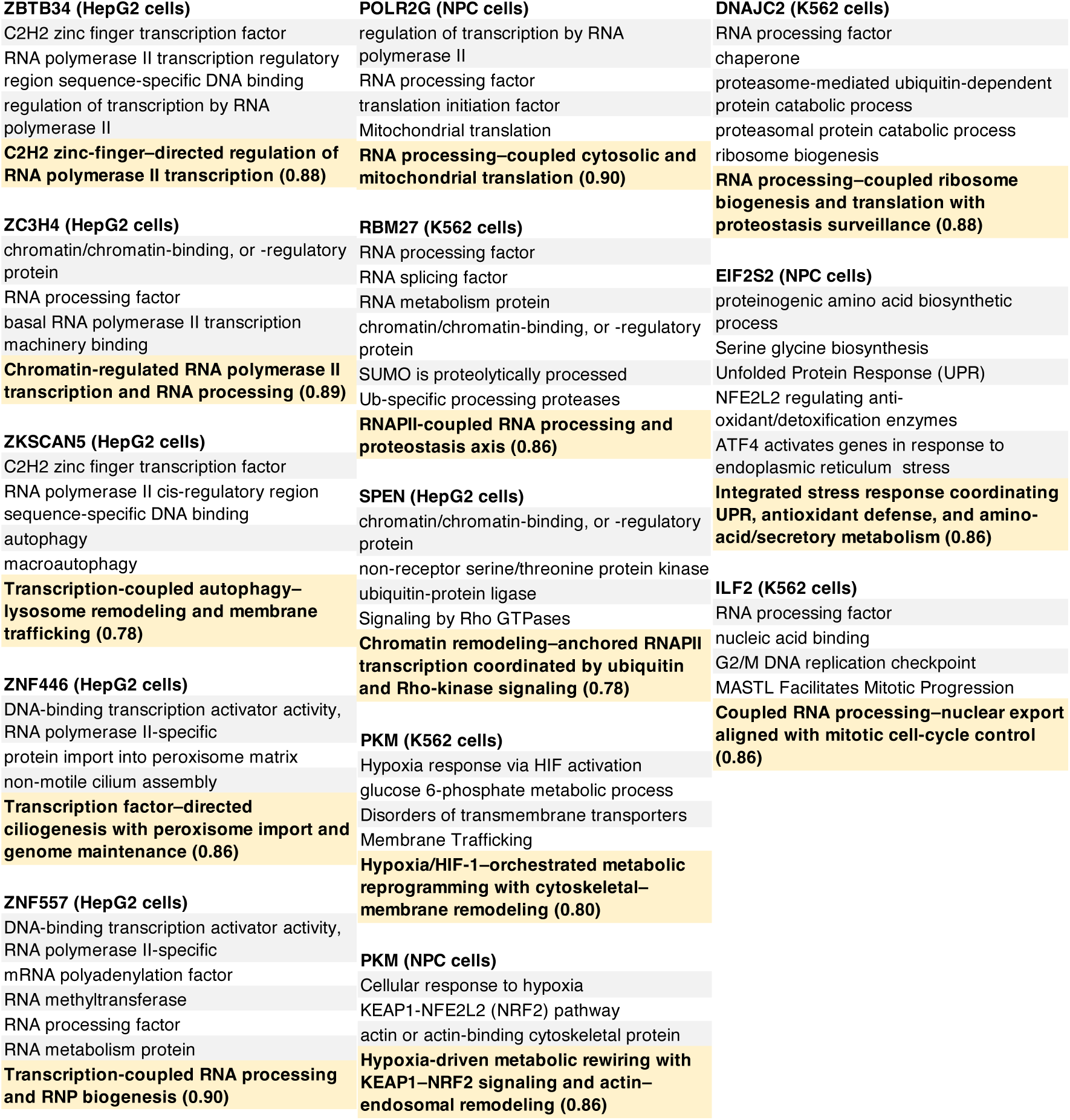
Functional roles inferred for nucleic acid–binding proteins through integration of UniProt annotations, large language model–based predictions, and confidence scores. Functional annotations derived from UniProt entries were systematically compared with significantly enriched biological processes among model-predicted NABP regulatory targets, as identified using the PANTHER Overrepresentation Test (see **Supplementary Table S5**). Shared, overlapping, or enrichment-associated biological functions between each nucleic acid–binding protein (NABP) and its predicted regulatory targets are indicated. Functional interpretations generated using a large language model–based gene set inference framework are highlighted in yellow. Confidence scores (shown in parentheses) represent the proportion of genes within each predicted target set that support the inferred functional annotation. Confidence categories were defined as follows: high (0.87–1.00), medium (0.82–0.86), low (0.01–0.81), or none (0). This integrative framework enables systematic comparison of known NABP functions with predicted context-specific regulatory programs, providing orthogonal validation of biologically coherent target inference. For methodological details, see Hu M. et al., *Nat. Methods*, 2025 ^31^.

**Table 2.**
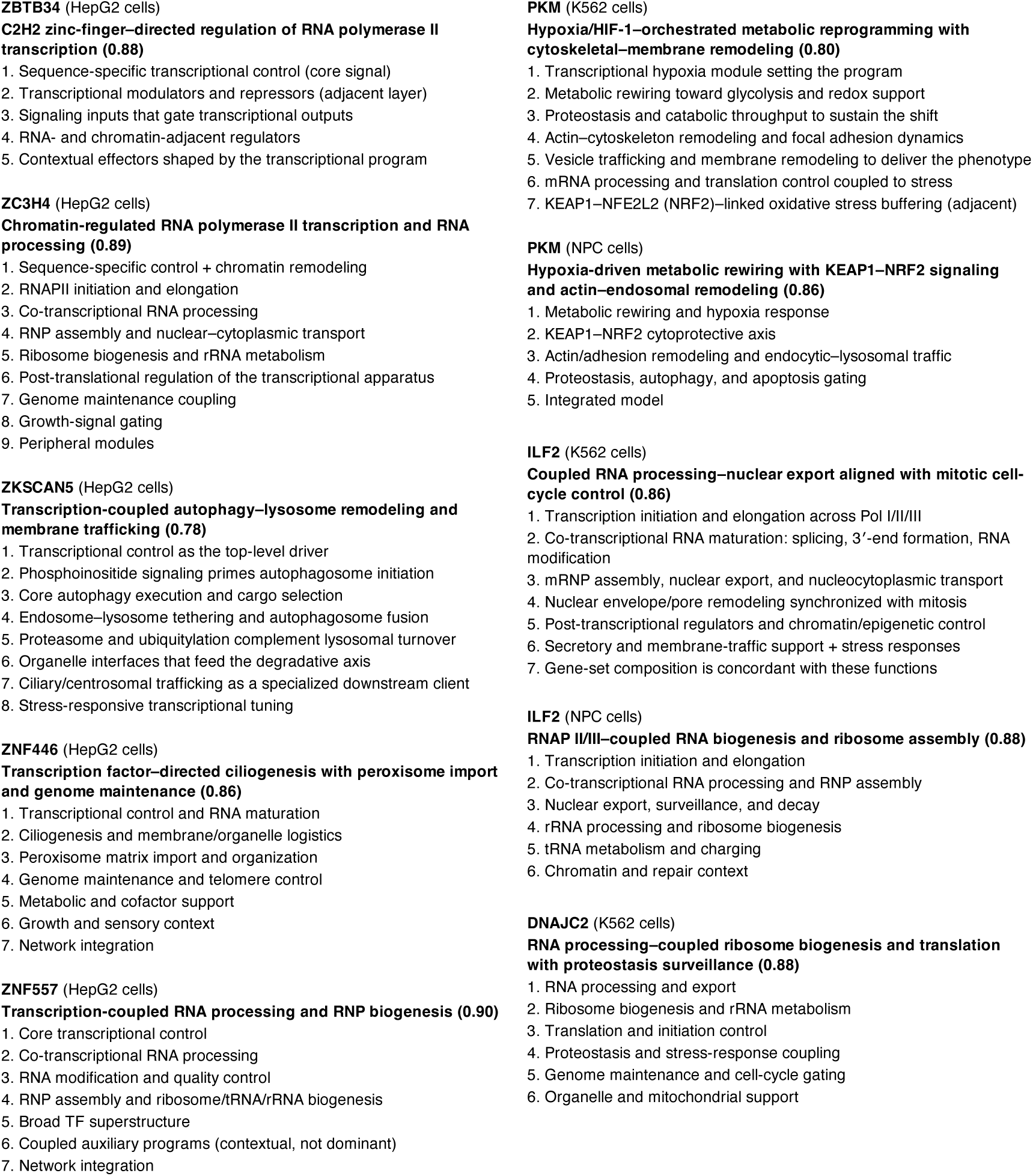
Functional characterization of predicted regulatory targets of nucleic acid-binding proteins using a large language model–based inference framework. Representative biological functions associated with predicted regulatory target genes were inferred for each nucleic acid–binding protein (NABP) using a large language model–based gene set interpretation framework. Major functional themes are summarized in bold for each NABP, while numbered subcategories indicate distinct biological processes, regulatory programs, or functional subdivisions associated with specific subsets of predicted target genes. This framework enables systematic interpretation of predicted NABP regulatory architectures by organizing complex gene set functions into hierarchically structured biological themes, thereby complementing conventional enrichment analyses with literature-guided functional synthesis.

### State–dependent functional specificity revealed by deep learning–refined NABP targets

Deep learning-based refinement further revealed cell type–specific biological functions associated with the regulatory targets of multiple RBPs. For example, the predicted targets of PKM exhibited distinct pathway enrichments profiles between K562 leukemia cells and neural progenitor cells (NPCs). Pathways including the *Pentose phosphate pathway*, *Glycolysis*, and *Hypoxia response mediated by HIF activation* were significantly enriched in K562 cells but not in NPCs, whereas the *Integrin signaling pathway*, *Glycogen metabolism*, and *Glutamate neurotransmitter release cycle* were specifically enriched in NPCs under a DeepLIFT background reference defined as 0.5× the original input feature values^32–41^ (**Fig. 5**, **Table 3**, and **Supplementary Table S5**). Large language model–based interpretation produced consistent functional assignments (**Supplementary Note 2**). Notably, these cell type–dependent associations closely align with established biological characteristics of cancer cells and neural progenitor cells.

**Figure 5.**
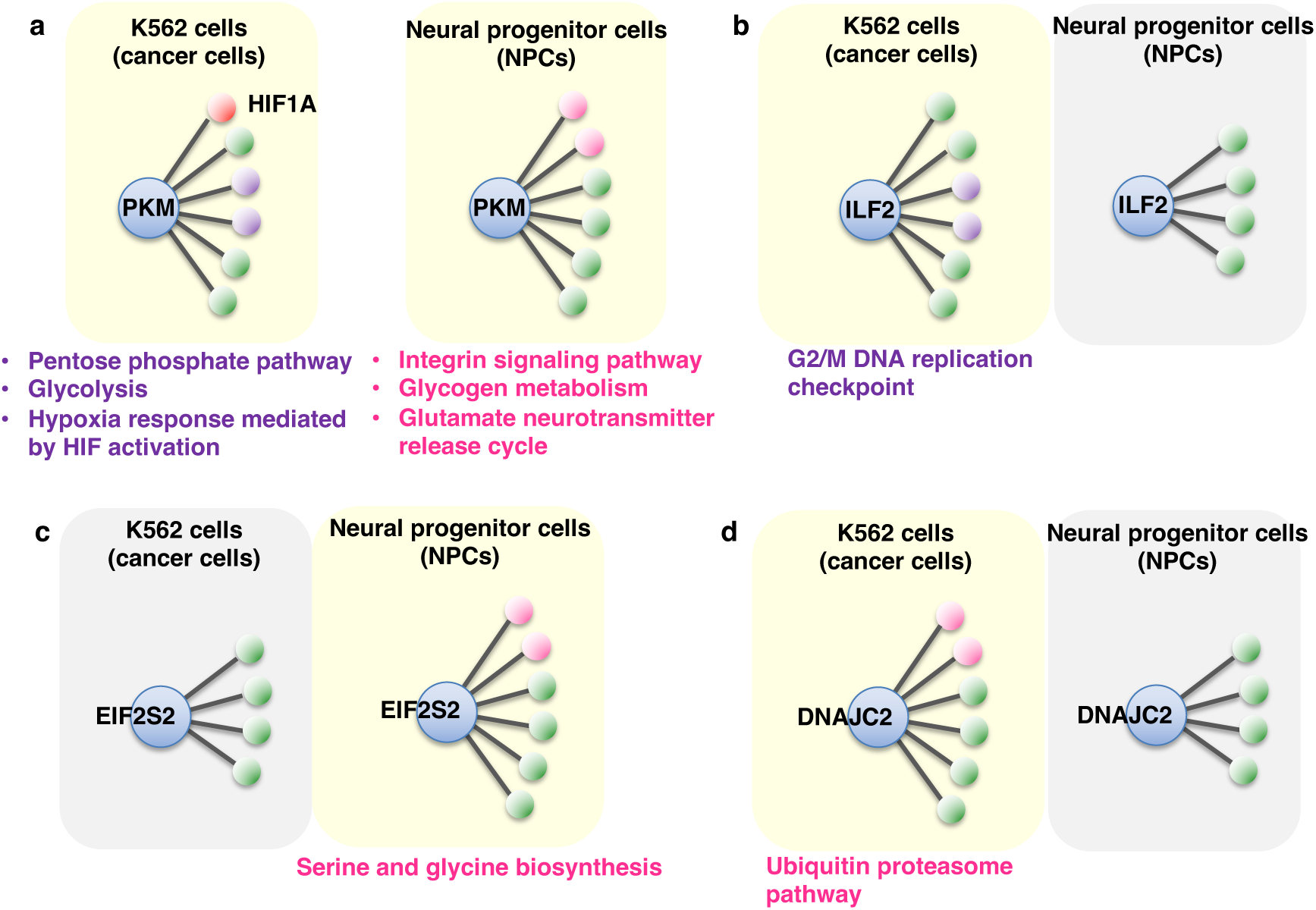
Functional mapping of nucleic acid-binding proteins reveals context-dependent regulatory roles in cancer and neural progenitor cells. A deep learning framework integrating transcriptomic co-expression relationships was used to infer the regulatory influence of nucleic acid–binding proteins (NABPs) on gene expression across distinct cellular contexts. DeepLIFT-derived contribution scores quantified the relative regulatory impact of each NABP across predicted target genes, and top-ranked targets were designated as primary regulatory candidates. Functional characterization of these predicted targets was performed using PANTHER-based enrichment analyses together with large language model–based functional inference, enabling identification of biologically coherent, cell type–specific regulatory pathways associated with individual NABPs. Representative examples illustrate distinct context-dependent regulatory specialization across K562 leukemia cells and neural progenitor cells (NPCs), including PKM-associated metabolic and hypoxia-related pathways in K562 cells versus neuronal and metabolic pathways in NPCs, ILF2-associated cell cycle checkpoint regulation in K562 cells, EIF2S2-associated amino acid biosynthesis in NPCs, and DNAJC2-associated ubiquitin–proteasome pathway enrichment in K562 cells. Together, these results demonstrate that deep learning–refined NABP target prediction enables systematic identification of cell type–dependent regulatory functions, revealing biologically meaningful context-specific regulatory architectures beyond co-expression alone.

**Table 3.**
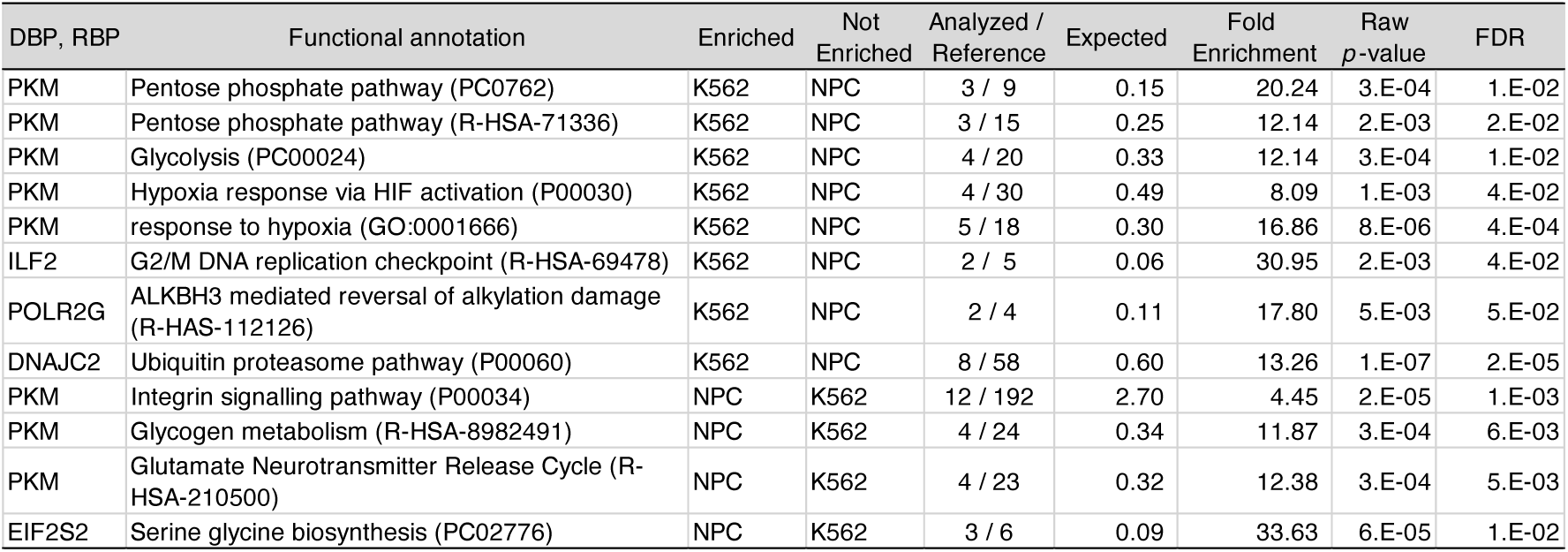
Functional enrichments of predicted regulatory target genes for nucleic acid-binding proteins. Functional enrichment analysis was performed using the PANTHER database to identify biologically significant annotations among regulatory target genes predicted by the deep learning framework incorporating gene co-expression–based NABP inference. This table presents representative examples of significantly enriched functional annotations associated with predicted regulatory targets for individual DNA-binding proteins (DBPs) and RNA-binding proteins (RBPs). For each annotation: ‘**Reference**’ indicates the total number of genes in the background reference dataset; ‘**Analyzed**’ denotes the number of predicted regulatory target genes tested for enrichment; ‘**Expected**’ represents the number of genes expected to contain the annotation by chance; ‘**Fold Enrichment**’ reflects the ratio of observed to expected gene counts; ‘**Raw *p*-value**’ corresponds to Fisher’s exact test significance; and ‘**FDR**’ indicates the Benjamini–Hochberg false discovery rate–adjusted significance. Together, these enrichments provide statistical validation that predicted NABP regulatory targets are functionally coherent and biologically relevant.

| DBP, RBP | Functional annotation | Enriched | Not Enriched | Analyzed / Reference | Expected | Fold Enrichment | Raw *p*-value | FDR |
| --- | --- | --- | --- | --- | --- | --- | --- | --- |
| PKM | Pentose phosphate pathway (PC0762) | K562 | NPC | 3 / 9 | 0.15 | 20.24 | 3.E-04 | 1.E-02 |
| PKM | Pentose phosphate pathway (R-HSA-71336) | K562 | NPC | 3 / 15 | 0.25 | 12.14 | 2.E-03 | 2.E-02 |
| PKM | Glycolysis (PC00024) | K562 | NPC | 4 / 20 | 0.33 | 12.14 | 3.E-04 | 1.E-02 |
| PKM | Hypoxia response via HIF activation (P00030) | K562 | NPC | 4 / 30 | 0.49 | 8.09 | 1.E-03 | 4.E-02 |
| PKM | response to hypoxia (GO:0001666) | K562 | NPC | 5 / 18 | 0.30 | 16.86 | 8.E-06 | 4.E-04 |
| ILF2 | G2/M DNA replication checkpoint (R-HSA-69478) | K562 | NPC | 2 / 5 | 0.06 | 30.95 | 2.E-03 | 4.E-02 |
| POLR2G | ALKBH3 mediated reversal of alkylation damage (R-HAS-112126) | K562 | NPC | 2 / 4 | 0.11 | 17.80 | 5.E-03 | 5.E-02 |
| DNAJC2 | Ubiquitin proteasome pathway (P00060) | K562 | NPC | 8 / 58 | 0.60 | 13.26 | 1.E-07 | 2.E-05 |
| PKM | Integrin signalling pathway (P00034) | NPC | K562 | 12 / 192 | 2.70 | 4.45 | 2.E-05 | 1.E-03 |
| PKM | Glycogen metabolism (R-HSA-8982491) | NPC | K562 | 4 / 24 | 0.34 | 11.87 | 3.E-04 | 6.E-03 |
| PKM | Glutamate Neurotransmitter Release Cycle (R-HSA-210500) | NPC | K562 | 4 / 23 | 0.32 | 12.38 | 3.E-04 | 5.E-03 |
| EIF2S2 | Serine glycine biosynthesis (PC02776) | NPC | K562 | 3 / 6 | 0.09 | 33.63 | 6.E-05 | 1.E-02 |

Additional RBPs exhibited similar distinct regulatory specificity. Predicted target genes of ILF2 were enriched for the *G2/M DNA replication checkpoint* in K562 cells but not in NPCs^42–44^; EIF2S2 target genes were specifically associated with *Serine and glycine biosynthesis* in NPCs but not in K562 cells^45,46^; and DNAJC2 target genes were enriched for the *Ubiquitin proteasome pathway* in K562 cells but not in NPCs^47,48^ (**Fig. 5, Table 3**, and **Supplementary Table S5**). Large language model–based functional inference again recapitulated these cell type–dependent patterns (**Supplementary Note 2**), further supporting consistency with prior biological knowledge.

Collectively, these results demonstrate that deep learning–based refinement of NABP target predictions enables systematic identification of cell type–dependent regulatory functions, revealing biologically meaningful regulatory specificity beyond that detectable from co-expression alone.

### Functional enrichment analysis of NABP target genes using contribution score–based GSEA

Gene set enrichment analysis (GSEA) based on DeepLIFT-derived contribution scores revealed robust, cell type–dependent functional redistribution among NABP regulatory targets. Rather than ranking genes by expression level, predicted targets were ranked according to their relative regulatory contribution within each cellular context, enabling pathway-level interpretation through shifts in the positional distribution of functionally related genes along contribution-ranked lists, independent of absolute score magnitude or sign (**Fig 6a**). Within this framework, PKM-associated pathways exhibited pronounced redistribution between neural progenitor cells (NPCs) and K562 leukemia cells. Multiple Reactome pathways displayed substantial differences in normalized enrichment scores (ΔNES = NES_K562 − NES_NPC), indicating marked divergence in inferred regulatory coupling across cellular contexts. Under the input × 0.5 background configuration, DeepLIFT contribution scores derived from co-expression–based inputs exhibited a negative correlation with gene expression levels. Consistent with this relationship, Neural System–associated pathways were significantly overrepresented among high ΔNES terms, whereas Immune System–associated pathways were preferentially enriched among low ΔNES terms, reflecting systematic context-dependent pathway redistribution (**Fig. 6b**, **Table 4**, and **Supplementary Fig. S5**). Importantly, ΔNES reflects redistribution in contribution-ranked pathway positioning rather than direct activation or repression of lineage-specific programs. Conceptually, ΔNES quantifies the relative positional shift of pathway-associated genes along regulatory contribution landscapes between cellular states.

**Figure 6.**
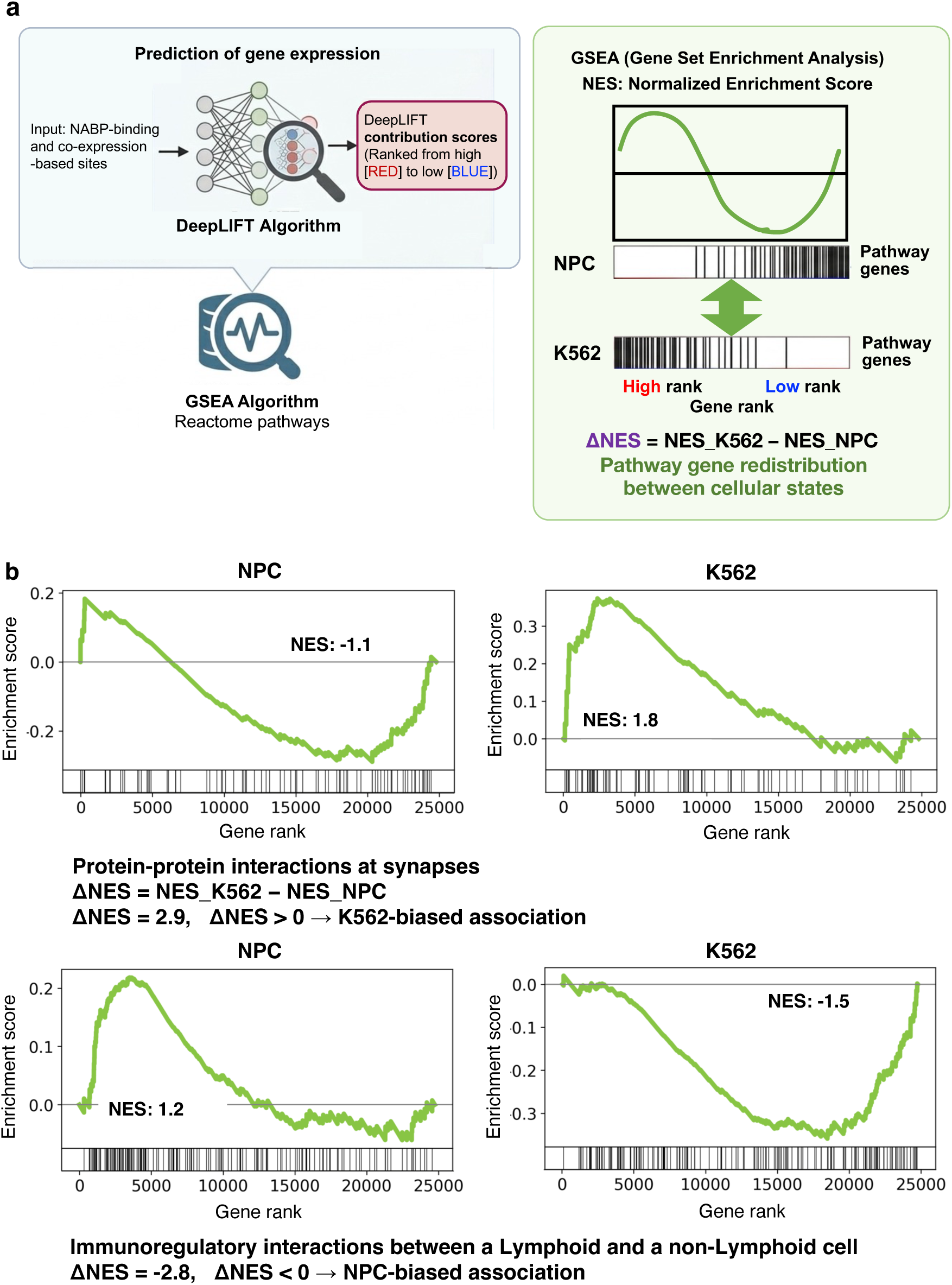
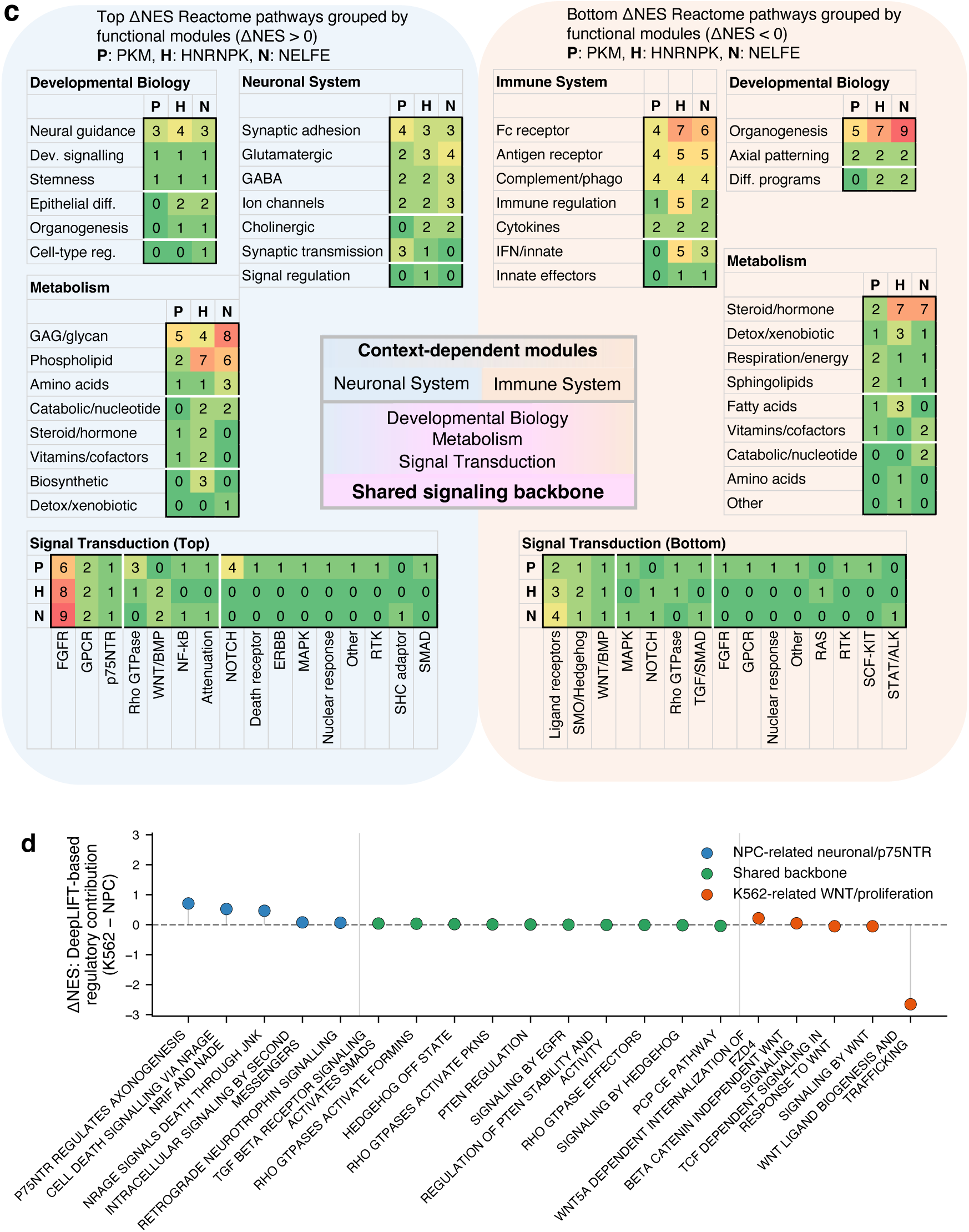
Pathway redistribution reveals a shared signaling backbone and context-dependent functional modules across RNA-binding proteins. **a. Conceptual framework of ΔNES-based pathway redistribution analysis.** DeepLIFT-derived gene contribution scores were used to rank genes for each RNA-binding protein (RBP) across biological states. Gene set enrichment analysis (GSEA) of Reactome pathways was then performed on contribution-ranked gene lists, and pathway redistribution between K562 leukemia cells and neural progenitor cells (NPCs) was quantified as ΔNES (NES_K562 − NES_NPC). This framework captures shifts in pathway-associated gene positioning along regulatory contribution landscapes rather than direct transcriptional activation or repression. **b. Representative GSEA enrichment plots illustrating context-dependent pathway redistribution.** Positive and negative ΔNES values indicate opposing positional shifts of pathway-associated genes between K562 and NPC regulatory landscapes. These redistribution patterns reflect changes in inferred regulatory coupling across cellular contexts rather than binary pathway activation states. Additional representative examples are provided in **Supplementary Figure S5**. **c. Functional module organization of ΔNES-ranked pathways across representative RBPs (PKM, HNRNPK, and NELFE).** Pathways were grouped into higher-order functional modules to facilitate biological interpretation. Across all three RBPs, Signal Transduction pathways consistently formed a conserved shared signaling backbone, whereas Neural System, Immune System, Developmental Biology, and metabolic modules exhibited pronounced context-dependent redistribution. These results support a hierarchical regulatory model in which RBPs modulate common upstream signaling architectures while redistributing downstream functional outputs across cellular states. **d. Distribution of Signal Transduction pathways along the ΔNES axis for PKM.** Shared signaling backbone pathways were defined as strongly enriched Signal Transduction pathways with near-zero ΔNES values (max |NES| ≥ 1.5), indicating conserved regulatory influence across cellular contexts. In contrast, pathways with large absolute ΔNES values represented context-specific signaling extensions associated with NPC-related neuronal modules or K562-associated proliferative/WNT-related programs. Ten representative shared backbone pathways and five representative context-specific pathways from both NPC- and K562-associated regulatory modules were selected based on ΔNES ranking and enrichment magnitude to illustrate the continuum from conserved backbone signaling to cell type–biased functional specialization.

**Table 4.**
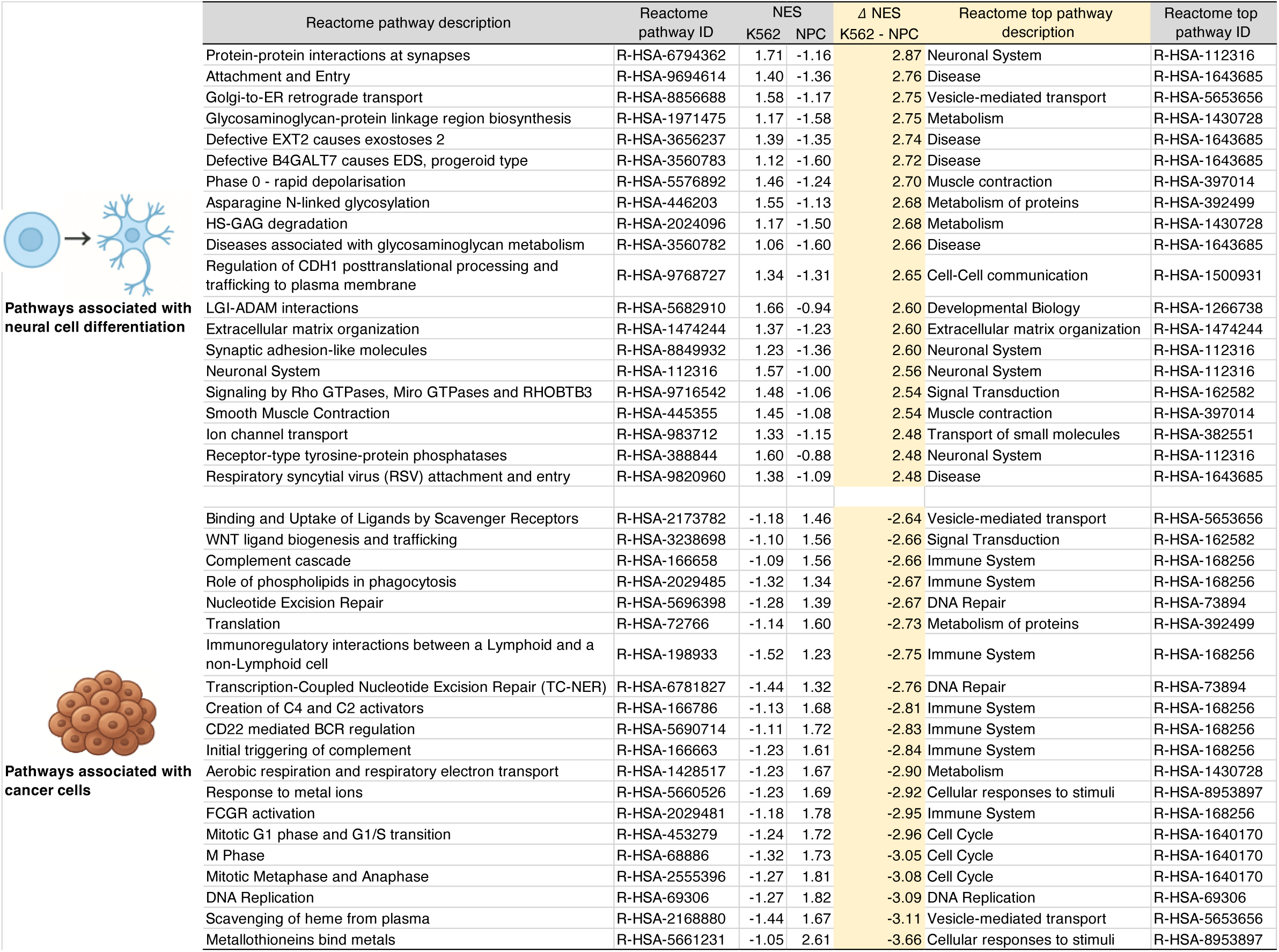
Cell type–dependent pathway redistribution identified by contribution score–based GSEA. Normalized enrichment score (NES) values were derived from gene set enrichment analysis (GSEA) performed on DeepLIFT contribution score–ranked gene lists. ΔNES represents the difference in pathway enrichment between K562 leukemia cells and neural progenitor cells (NPCs), calculated as ΔNES = NES_K562 − NES_NPC. This table summarizes representative pathway redistribution results for PKM, shown as a model RNA-binding protein consistent with the analyses presented in **Figure 6**. The twenty Reactome pathways with the highest positive and most negative ΔNES values are listed in the upper and lower sections, respectively. Pathways with strongly positive ΔNES values were predominantly enriched for Neural System– and developmental biology–associated functional categories, whereas pathways with strongly negative ΔNES values were primarily associated with immune regulation, proliferative signaling, cell cycle control, and genome maintenance. These opposing distributions reflect substantial context-dependent redistribution of inferred regulatory influence between NPC and K562 cellular states. Importantly, ΔNES values reflect shifts in pathway-associated gene positioning along regulatory contribution rankings rather than direct pathway activation or repression. Corresponding analyses using an amplified DeepLIFT background reference (input × 2) produced qualitatively consistent pathway redistribution patterns, with generally larger NES magnitudes due to increased numbers of non-zero predicted PKM-associated target genes (**Supplementary Tables S8 and S9**).

Because DeepLIFT contribution scores depend on background definition, we further evaluated alternative reference configurations (input × 0.5 and input × 2). Although background scaling altered contribution–expression polarity and the absolute direction of individual NES values, relative ΔNES-based pathway differences between NPC and K562 were preserved across configurations (**Fig. 6**; **Supplementary Tables S7–S9**). The amplified background reference (input × 2) increased the number of genes with non-negligible contribution scores and generally produced larger NES magnitudes, reflecting enhanced sensitivity without altering relative redistribution structure. These observations support the robustness of ΔNES as a comparative metric driven by stable rank-structure differences rather than background-dependent score polarity. Extending this analysis beyond PKM, similar category-level redistribution patterns were observed for additional RNA-binding proteins, including HNRNPK and NELFE (**Supplementary Tables S10–S13**). Across all three RBPs, Neural System–annotated pathways were significantly overrepresented among high ΔNES terms, whereas Immune System–annotated pathways were enriched among low ΔNES terms (one-sided Fisher’s exact test, *p* < 0.001; **Supplementary Table S14**). This reproducible enrichment pattern indicates systematic redistribution of regulatory influence between NPC and K562 contexts that is conserved across multiple RBPs and remains consistent at both pathway and higher-order functional module levels.

Notably, Signal Transduction emerged as the dominant top-level Reactome category across the ΔNES spectrum for all three RBPs (**Supplementary Table S14**). Subpathway analysis demonstrated that this enrichment was primarily driven by broadly utilized receptor-proximal signaling modules, including FGFR/RTK signaling, phospholipase C–mediated cascades, WNT/BMP pathways, and receptor-associated second-messenger systems. At the level of exact Reactome pathway identity, overlap across RBPs was limited, with only one pathway (FGFR1C ligand binding and activation) shared among PKM, HNRNPK, and NELFE, alongside a small number of pairwise overlaps such as NF-κB activation and signaling between PKM and NELFE (**Supplementary Table S16**). Consistent with this observation, pairwise comparison using the Jaccard similarity index revealed strong pathway-level similarity between HNRNPK and NELFE (J ≈ 0.65), whereas PKM shared comparatively limited exact pathway overlap with either protein (PKM–HNRNPK: J ≈ 0.13; PKM–NELFE: J ≈ 0.15), indicating substantial divergence at the level of individual pathway identity. However, functional module–level grouping revealed a markedly different organizational structure. Despite limited exact pathway overlap, all three RBPs converged on shared receptor-proximal signaling architectures, particularly involving FGFR signaling, phospholipase C cascades, ligand–receptor systems, and MAPK-related signaling processes. These findings indicate that RBPs share a conserved signaling backbone at the functional systems level, even when individual pathway identities diverge. Furthermore, PKM exhibited a broader and more diversified signaling architecture, incorporating additional regulatory modules related to Rho GTPase signaling, Notch signaling, and neuronal-associated pathways. In contrast, bottom-ranked pathways were predominantly restricted to more general signaling modules such as GPCR, WNT/BMP, and ligand-binding receptor pathways, consistent with leukemia- or hematopoietic-associated signaling states in K562 cells. Together, these findings support a hierarchical regulatory model in which a shared signaling backbone is differentially extended by cell type–biased functional modules, rather than being defined solely by exact pathway overlap.

To enable systematic interpretation, ΔNES-ranked pathways were grouped into functional modules based on shared biological processes (see **Methods** and **Supplementary Table S15**). This module-level organization revealed a clear separation between pathways with positive ΔNES values, which were predominantly associated with NPC-enriched functional pathways, and pathways with low negative ΔNES values, which were primarily associated with K562-enriched functional programs (**Fig. 6c**). Positive ΔNES modules were largely enriched for signaling- and interaction-related biological functions, including neuronal system–associated signaling, metabolic regulation, and stemness-related programs. In contrast, negative ΔNES modules were predominantly associated with immune system–related functions and developmental processes, including organogenesis and lineage specification. Despite this broad functional divergence, a subset of Signal Transduction pathways—including receptor tyrosine kinase (RTK), MAPK, and WNT-related signaling modules—was consistently detected across multiple RBPs, suggesting the presence of a broadly conserved signaling architecture. However, analysis along the ΔNES axis demonstrated that these signaling pathways were not uniformly redistributed (**Fig. 6d**; **Supplementary Fig. S6**; **Supplementary Table S18** and **S19**). Signal Transduction pathways with near-zero ΔNES values formed a shared signaling backbone, reflecting conserved regulatory contributions across cellular contexts, whereas pathways with large absolute ΔNES values represented context-dependent signaling extensions. Notably, neuronal system–associated pathways—including P75NTR- and NRAGE-related signaling for PKM^49,50^, RND3- and WNT-associated signaling for HNRNPK^51,52^, and NRIF-, TGFBR3-, and insulin-related signaling for NELFE^53–55^—generally exhibited more moderate ΔNES shifts, indicating shared directional redistribution with protein-specific variation. In contrast, K562-associated proliferative pathways, particularly WNT-related signaling modules, consistently displayed substantially larger ΔNES shifts, reflecting stronger context-dependent regulatory redistribution. This pattern suggests that modulation or repression of lineage-associated pathways may require comparatively limited pathway-level redistribution, whereas activation of proliferative or oncogenic programs may involve broader regulatory reorganization. Across RBPs, shared backbone pathways exhibited relatively stable functional specificity distributions, whereas context-dependent signaling modules displayed protein-specific shifts in leading-edge gene composition. Leading-edge genes represent the subset of pathway-associated genes that contribute most strongly to enrichment signals by appearing preferentially near one end of the DeepLIFT score–ranked gene list, thereby contributing to higher normalized enrichment scores (NES) (**Supplementary Fig. S7**). PKM exhibited highly specific positive ΔNES modules, whereas HNRNPK and NELFE showed distinct negative-module specialization patterns, indicating partially conserved yet protein-dependent regulatory redistribution. Together, these findings indicate that RBPs do not primarily regulate entirely distinct sets of cell-type-specific genes; rather, they modulate a common signaling framework while redistributing functional regulatory outputs across cellular contexts.

Consistent with established roles of PKM2 in metabolic rewiring, nuclear transcriptional regulation, and proliferative state maintenance ^56,57^, the observed ΔNES redistribution is compatible with context-dependent repositioning of metabolic and neuronal regulatory modules between leukemia and neural progenitor states ^36^. HNRNPK and NELFE, which coordinate transcriptional and post-transcriptional regulatory processes during development and malignancy, displayed parallel redistribution patterns, further supporting a shared regulatory architecture modulated across cellular contexts (**Supplementary Note and Tables section: Functional interpretation of ΔNES-based pathway redistribution across PKM, HNRNPK, and NELFE**). Collectively, these findings demonstrate that contribution score–based GSEA captures reproducible, category-level functional redistribution across cell types while revealing a conserved upstream signaling backbone shared among multiple RNA-binding proteins. This framework provides an interpretable and scalable strategy for dissecting context-dependent regulatory organization beyond expression-centric or threshold-dependent analytical approaches.

To further investigate the gene-level composition of enriched pathways, we examined leading-edge genes identified by GSEA for ΔNES-ranked Reactome pathways. Notably, many leading-edge genes were classified as shared functional genes rather than strictly cell-type-specific genes, even within pathways associated with neuronal or immune systems (**Supplementary Table S17**). These findings further support the model that RBPs predominantly regulate broadly shared gene networks while redistributing pathway-level functional outputs in a context-dependent manner.

## Discussion

Our findings suggest that RNA-binding proteins (RBPs) regulate gene expression programs through redistribution of regulatory influence across a shared signaling backbone, rather than by controlling entirely distinct pathway sets. Across multiple RBPs, including PKM, HNRNPK, and NELFE, Signal Transduction consistently forms a conserved signaling backbone, while functional modules associated with neuronal and immune processes are differentially positioned depending on cellular context. This hierarchical organization suggests that regulatory rewiring occurs through redistribution across common signaling networks.

Importantly, ΔNES captures this redistribution as shifts in the positional distribution of pathway-associated genes along contribution-ranked lists, rather than reflecting direct pathway activation or repression. Because gene expression represents the combined effects of multiple regulatory inputs, contribution scores quantify relative regulatory influence, enabling detection of subtle but systematic differences in regulatory architecture across biological states.

At the pathway level, the shared signaling backbone is characterized by broadly utilized receptor-mediated and growth factor–associated pathways, including RTK, MAPK, and WNT-related signaling. Although individual pathway identities vary across RBPs, functional module–level analysis reveals convergence on common upstream signaling processes. This structural convergence differentiates our findings from conventional enrichment-based analyses, which typically emphasize pathway identity rather than organization.

In contrast, modules associated with neuronal functions and immune responses exhibit context-dependent redistribution, indicating that RBPs modulate how shared gene networks are functionally deployed across cellular states. This model differs from typical transcription factor–mediated regulation, which often determine cellular identity through selective activation of more restricted gene sets. Instead, RBPs appear to regulate functional allocation within broadly shared signaling and metabolic networks.

The robustness of these patterns across alternative DeepLIFT background settings further supports the interpretation that pathway-level contrasts arise from stable rank-structure differences rather than background-dependent artifacts. While co-expression–based inference does not directly establish causal regulatory relationships, it captures functional coupling at the systems level, providing a scalable framework for analyzing regulatory architecture in the absence of experimentally defined binding information.

Looking forward, expanding this framework to a broader range of cell types and regulatory factors will enable systematic characterization of pathway redistribution across diverse biological contexts. Comparative analysis across transcription factors and RNA-binding proteins may further clarify their distinct and complementary roles in shaping regulatory architectures. Extending the framework to inter-individual variation and disease-associated transcriptomes, as demonstrated in related deep learning studies of gene expression regulation, may provide insight into the regulatory basis of phenotypic variation and disease susceptibility. Such analyses could facilitate the identification of early disease-associated shifts in regulatory architecture and support applications in diagnosis and preventive medicine.

Furthermore, by linking predicted regulatory targets to pathway-level redistribution patterns, this framework may contribute to the reconstruction of gene regulatory networks and enable in silico prediction of transcriptional responses to perturbations. These capabilities could support the identification of candidate therapeutic targets and guide strategies for modulating gene expression programs in a controlled manner.

Together, our results suggest that contribution score–based pathway redistribution provides a framework for uncovering conserved signaling architectures alongside context-dependent functional modules. Consistency between inferred redistribution patterns and previously reported biological functions of PKM, HNRNPK, and NELFE further supports the biological relevance of this approach. In contrast to conventional expression-centric analyses, which primarily capture changes in gene abundance, our framework emphasizes redistribution of regulatory influence across cellular contexts. By integrating interpretable deep learning with pathway-level analysis, this strategy enables systems-level investigation of regulatory organization beyond direct binding- or expression-based analyses. More broadly, pathway redistribution may provide a general framework for investigating regulatory reorganization across diverse biological state transitions, including development, disease progression, metabolic adaptation, and cellular responses to environmental or signaling stimuli.

## Methods

### Predictive modeling of gene expression across diverse cell types using deep learning and NABP binding information

To predict gene expression across multiple human cell types, we adapted a previously established deep learning framework that integrates nucleic acid–binding protein (NABP) binding site information, including DNA-binding proteins (DBPs) and RNA-binding proteins (RBPs), as regulatory input features. In this implementation, regulatory elements located both proximally and distally (up to ±1 Mb from transcription start sites [TSSs]) were incorporated as input features (**Fig. 1**)^25,58^. To minimize the confounding effects of cellular heterogeneity inherent to bulk tissue datasets, RNA-seq data were selected at the individual cell-type level. Publicly available RNA-seq datasets were obtained for four functionally distinct cell types: human foreskin fibroblasts (HFFs), mammary epithelial cells (HMECs), neural progenitor cells (NPCs), and either HepG2 hepatoma cells or K562 leukemia cells, depending on the regulatory protein class under investigation^59,60^. Gene expression values (FPKM) were calculated using the RSeQC toolkit based on GENCODE v19 annotations, and subsequently log₂-transformed following the addition of a small pseudocount to avoid undefined values. Consistent with the framework described by Tasaki et al.^25,58^, gene expression fold changes were calculated relative to the median expression level across the four cell types. For each primary model iteration, three common reference cell types (HFFs, HMECs, and NPCs) were included, together with either HepG2 (for DNA-binding protein analyses) or K562 (for RNA-binding protein analyses), depending on the availability of corresponding ChIP-seq or eCLIP validation datasets. These cell types were selected to capture diverse biological lineages and regulatory environments while maximizing compatibility with publicly available experimental binding datasets. For analyses specifically requiring direct comparison of DeepLIFT contribution score distributions across multiple cell types for the same NABP—such as the evaluation of cell type–dependent regulatory differences (e.g., **Fig. 3d**)—an alternative four-cell-type configuration was employed, consisting of HFFs, NPCs, HepG2, and K562. This unified modeling framework enabled direct cross-cell-type comparison of contribution score distributions between HepG2 and K562 while preserving a consistent shared biological reference context.

### Extended regulatory features and DeepLIFT-compatible model design

To capture distal regulatory signals beyond promoter-proximal regions, we integrated DNA-binding site information from putative enhancer regions into a deep learning framework for gene expression prediction. Building upon our previous work^25,58^, this implementation extends the architecture originally proposed by Tasaki et al. ^25,58^, which primarily focused on promoter-associated regions and incorporated RNA-binding protein and miRNA features. In our earlier study, aimed to identify insulator-associated DBPs, RBPs and miRNAs were intentionally excluded to prioritize DNA-centered regulatory architecture. Accordingly, we constructed a unified feature matrix combining DNA-binding site information from both promoter regions (-2 kb to +1 kb) and enhancer-associated regions (up to ±1 Mb from the transcription start site [TSS]), thereby enabling DeepLIFT analysis under the high-accuracy “genomics-default” configuration (**Supplementary Figure S8**). In this study, DeepLIFT was used to calculate attribution values for each input feature associated with gene expression prediction. These values are referred to throughout this manuscript as DeepLIFT scores, contribution scores, or importance scores, depending on analytical context.

Binding site coordinates were obtained from the GTRD database (v20.06)^61^, and regulatory associations between distal binding sites and target genes were inferred using cis-expression quantitative trait loci (cis-eQTLs) from GTEx v8. A ChIP-seq peak overlapping a 100-bp window centered on an expression-associated SNP was considered functionally linked to the corresponding transcript. Trans-eQTLs were excluded, and genomic coordinates were converted to the hg19 reference assembly using UCSC LiftOver^62^. To maximize regulatory coverage while controlling feature dimensionality, peak lengths for each DBP were calculated in 100-bp bins across ±1 Mb surrounding each TSS, and the maximum peak value within each 10-kb interval was retained, yielding up to 200 distal regulatory bins. These distal bins were concatenated with 30 promoter-region bins to generate a unified 230-bin input layer. This single-layer input architecture was required because DeepLIFT’s genomics-default configuration (“RevealCancel” rule for dense layers and “Rescale” rule for convolutional layers) does not support models with multiple independent input layers. Consolidating all DBP-related features into a single matrix therefore preserved DeepLIFT compatibility while enhancing model interpretability.

Model training followed the protocol established by Tasaki et al. and applied in our prior studies^25,58^. Specifically, 2,705 genes located on chromosome 1 were reserved as an independent test set, while 22,251 genes from all other chromosomes were randomly assigned to the training set and 2,472 genes to the validation set. Models were implemented using TensorFlow 2.6.0 and trained on an NVIDIA RTX 3080 GPU, with typical training times ranging from approximately 2 to 4 hours. Feature attribution was assessed using DeepLIFT, which assigned contribution scores to each input feature based on its relative influence on predicted gene expression.

### Predicting NABP regulatory targets using gene co-expression–informed substitutions

To infer regulatory targets of nucleic acid–binding proteins (NABPs) without relying exclusively on direct experimental binding assays, we implemented a gene co-expression–driven modification of the deep learning input architecture. Specifically, DBPs exhibiting low contribution scores in the original model were selectively replaced with NABPs whose putative regulatory interactions were inferred from gene co-expression relationships. Among the 1,310 DBPs initially included in the baseline framework, 302 were substituted with alternative DBPs in one model configuration, while 325 were replaced with RBPs identified from the ENPD database^24^ in a separate RBP-focused analysis. Putative NABP–target associations were established using gene co-expression profiles from COXPRESdb^15^, where a Pearson’s z-score > 2 between a NABP-encoding gene and a non-NABP gene was used to define a strong co-expression relationship. These co-expression pairs were treated as proxies for regulatory interactions and incorporated into the model as synthetic regulatory input features. To enable integration into the existing DeepLIFT-compatible framework, hypothetical NABP binding sites were assigned to the transcription start site (TSS) and the 2-kb upstream promoter region of each co-expressed target gene. The modified model was subsequently retrained, and DeepLIFT was used to calculate feature-level attribution values (hereafter referred to as contribution scores), quantifying the relative influence of each inferred NABP–gene interaction on predicted gene expression outputs. Importantly, these contribution scores represent model-derived feature attributions rather than direct measurements of gene expression, which reflects the integrated effects of multiple regulatory inputs. This strategy enables functional characterization of NABPs lacking experimentally defined binding motifs or high-quality binding datasets by leveraging co-expression structure as an interpretable proxy for regulatory targeting.

### Validation of predicted NABP targets using experimentally determined binding

To assess the biological relevance of predicted regulatory targets, deep learning–inferred NABP–gene associations were systematically compared with experimentally validated binding datasets. Specifically, spatial overlap was evaluated between predicted target genes and ChIP-seq peaks for 32 DBPs (within ±5 kb of transcription start sites [TSSs]) as well as eCLIP peaks for 29 RBPs (across transcribed gene regions spanning TSS to transcription end site [TES]). Gene structure annotations were derived from GENCODE v19, and experimentally validated binding site datasets were obtained from the ENCODE portal and relevant prior studies (**Supplementary Table S6**) ^63,64^. This orthogonal validation framework provided an independent assessment of predicted regulatory target accuracy and supported the utility of gene co-expression–based deep learning models for inferring biologically meaningful NABP–target regulatory relationships.

### Prioritization of NABP regulatory targets based on importance score rankings

To identify the primary regulatory targets of each NABP, genes were prioritized according to DeepLIFT-derived contribution score rankings assigned to NABP-associated regulatory features. Genes whose associated binding sites were consistently ranked among the top or bottom scoring targets for a given NABP were selected as putative primary regulatory targets, representing strong positive or negative inferred regulatory influence. In most analyses, genes associated with the top five or bottom five ranked contribution scores were designated as high-confidence targets. To maintain sufficient statistical power for downstream functional enrichment analyses, these rank thresholds were adaptively adjusted for individual NABPs when necessary. For example, in cases where fewer high-confidence targets were initially identified, thresholds were relaxed (e.g., to the top or bottom ten genes) to ensure adequate target set size while preserving biological specificity.

### Functional enrichment analysis of NABP regulatory targets

To evaluate the biological coherence of predicted NABP regulatory targets, functional enrichment analyses were performed using the Statistical Overrepresentation Test implemented in the PANTHER database, together with custom in-house analytical pipelines^29^. Gene Ontology (GO) terms spanning biological process, molecular function, and cellular component categories were assessed across three gene sets: (1) primary NABP regulatory target genes identified by the deep learning model; (2) all co-expressed genes (Pearson’s z-score > 2); and (3) randomly selected gene sets of matched size. Enrichment significance was evaluated using Fisher’s exact test with Benjamini–Hochberg false discovery rate (FDR) correction, with FDR < 0.05 considered statistically significant. Human gene symbols were mapped to Ensembl Gene IDs using HGNC resources, and GO annotations were obtained from the Gene Ontology database^27,28^. Predicted target genes served as the test set, while the full gene expression dataset used for model training served as the reference background. To further contextualize these enrichment results and derive integrative functional interpretations, we subsequently applied a large language model–based gene set interpretation framework (see below).

### Large language model–assisted gene set interpretation

To complement conventional statistical analyses, we applied a large language model–based framework to interpret predicted NABP target gene sets. Gene lists and PANTHER-derived functional annotation summaries were provided as input to ChatGPT, which generated concise functional descriptions and candidate process labels with associated confidence scores using curated prompts (detailed in Extended Data Tables 1 and 5). A secondary prompt was used to identify genes supporting each inferred process based on explicit, non-speculative statements in the generated text. This workflow was implemented using a modified version of the “4.Reference search and validation.ipynb” pipeline^31^ and is publicly available via Code Ocean and GitHub (https://doi.org/10.24433/CO.7045777.v1; https://github.com/idekerlab/llm_evaluation_for_gene_set_interpretation). All model outputs were reviewed and curated by the authors. Conventional enrichment analyses (PANTHER and GSEA) served as the primary basis for biological interpretation, whereas the language model was used solely to assist functional summarization and provide an orthogonal layer of interpretability. Large language model–based interpretations were generally consistent with conventional enrichment analyses and yielded biologically coherent functional themes, supporting their use as a complementary interpretative approach for predicted NABP regulatory targets. The overall analytical workflow comprised three complementary steps: (i) statistical evaluation of functional enrichment using PANTHER, (ii) large language model–assisted interpretation to contextualize and extend enrichment results, including identification of functionally related signals beyond strict statistical thresholds, and (iii) gene set enrichment analysis (GSEA) to characterize pathway-level redistribution and global regulatory patterns across cellular contexts.

### Contribution score–based gene set enrichment analysis (GSEA)

Gene set enrichment analysis (GSEA) was performed using the preranked GSEA algorithm implemented in GSEApy (v1.1.11). For each DNA-binding protein (DBP) and RNA-binding protein (RBP), DeepLIFT contribution scores were calculated across all genes from trained deep learning models. Because raw DeepLIFT contribution scores vary substantially in scale and frequently include zero-valued features, scores were normalized on a per-gene basis by scaling values across all 1,310 DBPs/RBPs to a continuous range from −1 to 1 using min–max normalization. Genes were then ranked according to normalized contribution scores specific to the NABP of interest. These ranked gene lists were analyzed using the Reactome (v2026) gene set library with the following parameters: [1] 10,000 permutations; [2] weighted enrichment statistic (weight = 1); [3] Minimum gene set size of 10; and [4] at least three genes per pathway with non-zero contribution scores. In this framework, GSEA was used primarily to calculate normalized enrichment scores (NES) as continuous quantitative measures of pathway-associated regulatory contribution, rather than as formal significance tests of pathway enrichment. NES values were subsequently used for comparative downstream analyses, including ΔNES calculations across cellular contexts. This strategy enabled systematic identification of pathways associated with high positive or negative inferred regulatory influence and facilitated pathway redistribution analysis across cell states. Because the primary objective was comparative interpretation rather than statistical inference, nominal *p*-values and FDR were not used as primary criteria for pathway prioritization. Unlike conventional pathway activity inference approaches such as GSVA^65^ or AUCell^66^, which rank genes based on expression levels, this framework ranks genes according to deep learning–derived regulatory contribution scores, thereby enabling pathway-level analysis of inferred regulatory architecture rather than transcriptional abundance. Pairwise similarity between Signal Transduction subpathway sets was quantified using the Jaccard similarity index, defined as the ratio of intersection size to union size between pathway sets.

### Definition of functional modules for Reactome pathways

To facilitate systematic interpretation of pathway-level results, individual Reactome pathways were grouped into higher-order functional modules based on shared biological processes. Functional modules were manually curated through integrated evaluation of Reactome pathway annotations, pathway nomenclature, hierarchical category structure, and established biological relationships among pathways. Initially, pathways within the same top-level Reactome categories were examined and subsequently subdivided into biologically coherent submodules, such as synaptic signaling, receptor tyrosine kinase signaling, immune receptor signaling, and metabolic regulatory programs. This hierarchical grouping strategy reduced redundancy among closely related pathways while improving interpretability and visualization of large-scale pathway enrichment patterns. Each Reactome pathway was assigned to a single curated functional module, and corresponding abbreviated module labels (“module short”) were defined for compact representation in figures and comparative analyses. The complete mapping between Reactome pathways and functional modules is provided in **Supplementary Table S15**. Although this module classification involved manual curation, assignments were guided by established biological knowledge, Reactome ontology structure, and cross-protein consistency, thereby ensuring reproducibility and conceptual coherence across pathway categories and RBPs.

### Statistical analysis

Functional enrichment analyses were performed using Fisher’s exact test, with multiple hypothesis correction applied using the Benjamini–Hochberg false discovery rate (FDR) procedure. Statistical significance was defined as FDR < 0.05. Distributions of DeepLIFT contribution scores between gene co-expression–derived regulatory sites and experimentally validated ChIP-seq or eCLIP binding sites were compared using the Mann–Whitney U test (Wilcoxon rank-sum test), according to the relevant nucleic acid–binding protein class. A nominal *p*-value < 0.05 was considered statistically significant. Pairwise similarity between Signal Transduction subpathway sets was quantified using the Jaccard similarity index, defined as the ratio of the intersection size to the union size of compared pathway sets.

## Data availability

RNA-seq data for human foreskin fibroblast (HFF) cells were obtained from the Gene Expression Omnibus (GEO) under accession number GSM941744. RNA-seq data for human mammary epithelial cells (HMECs) were retrieved from ENCODE via the UCSC Genome Browser (http://genome.ucsc.edu/encode/) under the file name *wgEncodeCshlLongRnaSeqHmecCellPapAlnRep1.bam*. RNA-seq data for human neural progenitor cells (NPCs) were obtained from GEO under accession number GSM915326. RNA-seq data for HepG2 and K562 cells were obtained from the ENCODE portal under accession numbers ENCFF670LIE and ENCFF584WML, respectively. mRNA co-expression data were downloaded from COXPRESdb v7 (https://zenodo.org/record/6861444). The list of nucleic acid-binding proteins was obtained from the Eukaryotic Nucleic acid Binding Protein (ENPD) database (http://qinlab.sls.cuhk.edu.hk/ENPD/). ChIP-seq data for DNA-binding proteins were downloaded from the GTRD database (v20.06 release; https://gtrd.biouml.org). cis-eQTL data (GTEx_Analysis_v8_eQTL.tar) were obtained from the GTEx portal (v8 release; https://gtexportal.org). eCLIP data for RNA-binding proteins were retrieved from the ENCODE portal (accession ENCSR456FVU) as well as from Supplementary Data 2 of Van Nostrand et al., *Nature* (2020), https://doi.org/10.1038/s41586-020-2077-3.

## Code availability

The deep learning model implemented in this study is available at https://github.com/reposit2/coexgene.

## Acknowledgements

Supercomputing resources were provided by the Human Genome Center, the Institute of Medical Science, the University of Tokyo, and the ROIS National Institute of Genetics.

## Author contributions

N.O. designed and conducted the research, and wrote and revised the manuscript. K.S. revised the manuscript.

## Supplementary data

Supplementary data are available at

## References

1 Belz, G. T. & Nutt, S. L. Transcriptional programming of the dendritic cell network. Nature reviews. Immunology 12, 101‒113 (2012). 10.1038/nri3149

2 Miller, J. C. et al. Deciphering the transcriptional network of the dendritic cell lineage. Nature immunology 13, 888‒899 (2012). 10.1038/ni.2370

3 Ravasi, T. et al. An atlas of combinatorial transcriptional regulation in mouse and man. Cell 140, 744‒752 (2010). 10.1016/j.cell.2010.01.044

4 Gertz, J. et al. Distinct properties of cell-type-specific and shared transcription factor binding sites. Molecular cell 52, 25‒36 (2013). 10.1016/j.molcel.2013.08.037

5 Lambert, S. A. et al. The Human Transcription Factors. Cell 172, 650‒665 (2018). 10.1016/j.cell.2018.01.029

6 Vaquerizas, J. M., Kummerfeld, S. K., Teichmann, S. A. & Luscombe, N. M. A census of human transcription factors: function, expression and evolution. Nature reviews. Genetics 10, 252‒263 (2009). 10.1038/nrg2538

7 Pierson, E. et al. Sharing and Specificity of Co-expression Networks across 35 Human Tissues. PLoS Comput Biol 11, e1004220 (2015). 10.1371/journal.pcbi.1004220

8 Sonawane, A. R. et al. Understanding Tissue-Specific Gene Regulation. Cell Rep 21, 1077‒1088 (2017). 10.1016/j.celrep.2017.10.001

9 Saha, A. et al. Co-expression networks reveal the tissue-specific regulation of transcription and splicing. Genome research 27, 1843‒1858 (2017). 10.1101/gr.216721.116

10 Boyle, A. P. et al. Comparative analysis of regulatory information and circuits across distant species. Nature 512, 453‒456 (2014). 10.1038/nature13668

11 Kribelbauer, J. F., Rastogi, C., Bussemaker, H. J. & Mann, R. S. Low-Affinity Binding Sites and the Transcription Factor Specificity Paradox in Eukaryotes. Annu Rev Cell Dev Biol 35, 357‒379 (2019). 10.1146/annurev-cellbio-100617-062719

12 Ray, D. et al. RNA-binding proteins that lack canonical RNA-binding domains are rarely sequence-specific. Sci Rep 13, 5238 (2023). 10.1038/s41598-023-32245-9

13 Zeke, A. et al. Deep structural insights into RNA-binding disordered protein regions. Wiley Interdiscip Rev RNA 13, e1714 (2022). 10.1002/wrna.1714

14 Zaharias, S. et al. Intrinsically disordered electronegative clusters improve stability and binding specificity of RNA-binding proteins. J Biol Chem 297, 100945 (2021). 10.1016/j.jbc.2021.100945

15 Obayashi, T., Kodate, S., Hibara, H., Kagaya, Y. & Kinoshita, K. COXPRESdb v8: an animal gene coexpression database navigating from a global view to detailed investigations. Nucleic acids research 51, D80‒D87 (2023). 10.1093/nar/gkac983

16 Yang, S. et al. COEXPEDIA: exploring biomedical hypotheses via co-expressions associated with medical subject headings (MeSH). Nucleic acids research 45, D389‒D396 (2017). 10.1093/nar/gkw868

17 Zhu, Q. et al. Targeted exploration and analysis of large cross-platform human transcriptomic compendia. Nature methods 12, 211‒214, 213 p following 214 (2015). 10.1038/nmeth.3249

18 Miller, H. E. & Bishop, A. J. R. Correlation AnalyzeR: functional predictions from gene co-expression correlations. BMC Bioinformatics 22, 206 (2021). 10.1186/s12859-021-04130-7

19 Raina, P. et al. GeneFriends: gene co-expression databases and tools for humans and model organisms. Nucleic acids research 51, D145‒D158 (2023). 10.1093/nar/gkac1031

20 Stuart, J. M., Segal, E., Koller, D. & Kim, S. K. A gene-coexpression network for global discovery of conserved genetic modules. Science (New York, N.Y.) 302, 249‒255 (2003). 10.1126/science.1087447

21 Lemoine, G. G., Scott-Boyer, M. P., Ambroise, B., Perin, O. & Droit, A. GWENA: gene co-expression networks analysis and extended modules characterization in a single Bioconductor package. BMC Bioinformatics 22, 267 (2021). 10.1186/s12859-021-04179-4

22 Warde-Farley, D. et al. The GeneMANIA prediction server: biological network integration for gene prioritization and predicting gene function. Nucleic acids research 38, W214‒220 (2010). 10.1093/nar/gkq537

23 Wong, A. K., Krishnan, A. & Troyanskaya, O. G. GIANT 2.0: genome-scale integrated analysis of gene networks in tissues. Nucleic acids research 46, W65‒W70 (2018). 10.1093/nar/gky408

24 Tak Leung, R. W., Jiang, X., Chu, K. H. & Qin, J. ENPD - A Database of Eukaryotic Nucleic Acid Binding Proteins: Linking Gene Regulations to Proteins. Nucleic acids research 47, D322‒D329 (2019). 10.1093/nar/gky1112

25 Tasaki, S., Gaiteri, C., Mostafavi, S. & Wang, Y. Deep learning decodes the principles of differential gene expression. Nat Mach Intell 2, 376‒386 (2020). 10.1038/s42256-020-0201-6

26 Shrikumar, A., Greenside, P. & Kundaje, A. in Proceedings of the 34th International Conference on Machine Learning Vol. 70 (eds Precup Doina & Teh Yee Whye) 3145‒‒3153 (PMLR, Proceedings of Machine Learning Research, 2017).

27 Ashburner, M. et al. Gene ontology: tool for the unification of biology. The Gene Ontology Consortium. Nature genetics 25, 25‒29 (2000). 10.1038/75556

28 Gene Ontology, C., et al. The Gene Ontology knowledgebase in 2023. Genetics 224 (2023). 10.1093/genetics/iyad031

29 Thomas, P. D. et al. PANTHER: Making genome-scale phylogenetics accessible to all. Protein Sci 31, 8‒22 (2022). 10.1002/pro.4218

30 UniProt, C. UniProt: the Universal Protein Knowledgebase in 2025. Nucleic acids research 53, D609‒D617 (2025). 10.1093/nar/gkae1010

31 Hu, M. et al. Evaluation of large language models for discovery of gene set function. Nature methods 22, 82‒91 (2025). 10.1038/s41592-024-02525-x

32 Zahra, K., Dey, T., Ashish, Mishra, S. P. & Pandey, U. Pyruvate Kinase M2 and Cancer: The Role of PKM2 in Promoting Tumorigenesis. Front Oncol 10, 159 (2020). 10.3389/fonc.2020.00159

33 Zhang, Z. et al. PKM2, function and expression and regulation. Cell Biosci 9, 52 (2019). 10.1186/s13578-019-0317-8

34 Alquraishi, M. et al. Pyruvate kinase M2: A simple molecule with complex functions. Free Radic Biol Med 143, 176‒192 (2019). 10.1016/j.freeradbiomed.2019.08.007

35 Azoitei, N. et al. PKM2 promotes tumor angiogenesis by regulating HIF-1alpha through NF-kappaB activation. Mol Cancer 15, 3 (2016). 10.1186/s12943-015-0490-2

36 He, P. et al. PKM2 is a key factor to regulate neurogenesis and cognition by controlling lactate homeostasis. Stem Cell Reports 20, 102381 (2025). 10.1016/j.stemcr.2024.11.011

37 Bonar, N. A. & Petersen, C. P. Integrin suppresses neurogenesis and regulates brain tissue assembly in planarian regeneration. Development 144, 784‒794 (2017). 10.1242/dev.139964

38 Porcheri, C., Suter, U. & Jessberger, S. Dissecting integrin-dependent regulation of neural stem cell proliferation in the adult brain. J Neurosci 34, 5222‒5232 (2014). 10.1523/JNEUROSCI.4928-13.2014

39 Zheng, X. et al. Metabolic reprogramming during neuronal differentiation from aerobic glycolysis to neuronal oxidative phosphorylation. eLife 5 (2016). 10.7554/eLife.13374

40 Agostini, M. et al. Metabolic reprogramming during neuronal differentiation. Cell Death Differ 23, 1502‒1514 (2016). 10.1038/cdd.2016.36

41 Saez, I. et al. Neurons have an active glycogen metabolism that contributes to tolerance to hypoxia. J Cereb Blood Flow Metab 34, 945‒955 (2014). 10.1038/jcbfm.2014.33

42 Matthews, H. K., Bertoli, C. & de Bruin, R. A. M. Cell cycle control in cancer. Nat Rev Mol Cell Biol 23, 74‒88 (2022). 10.1038/s41580-021-00404-3

43 Dominguez-Brauer, C. et al. Targeting Mitosis in Cancer: Emerging Strategies. Molecular cell 60, 524‒536 (2015). 10.1016/j.molcel.2015.11.006

44 Levine, M. S. & Holland, A. J. The impact of mitotic errors on cell proliferation and tumorigenesis. Genes Dev 32, 620‒638 (2018). 10.1101/gad.314351.118

45 Huang, X. et al. D-Serine regulates proliferation and neuronal differentiation of neural stem cells from postnatal mouse forebrain. CNS Neurosci Ther 18, 4‒13 (2012). 10.1111/j.1755-5949.2011.00276.x

46 El-Hattab, A. W. Serine biosynthesis and transport defects. Mol Genet Metab 118, 153‒159 (2016). 10.1016/j.ymgme.2016.04.010

47 Kaida, A. & Iwakuma, T. Regulation of p53 and Cancer Signaling by Heat Shock Protein 40/J-Domain Protein Family Members. Int J Mol Sci 22 (2021). 10.3390/ijms222413527

48 Sterrenberg, J. N., Blatch, G. L. & Edkins, A. L. Human DNAJ in cancer and stem cells. Cancer Lett 312, 129‒142 (2011). 10.1016/j.canlet.2011.08.019

49 Salehi, A. H. et al. NRAGE, a novel MAGE protein, interacts with the p75 neurotrophin receptor and facilitates nerve growth factor-dependent apoptosis. Neuron 27, 279‒288 (2000). 10.1016/s0896-6273(00)00036-2

50 Zuccaro, E. et al. Polarized expression of p75(NTR) specifies axons during development and adult neurogenesis. Cell Rep 7, 138‒152 (2014). 10.1016/j.celrep.2014.02.039

51 Pacary, E., Azzarelli, R. & Guillemot, F. Rnd3 coordinates early steps of cortical neurogenesis through actin-dependent and -independent mechanisms. Nat Commun 4, 1635 (2013). 10.1038/ncomms2614

52 Hirabayashi, Y. et al. The Wnt/beta-catenin pathway directs neuronal differentiation of cortical neural precursor cells. Development 131, 2791‒2801 (2004). 10.1242/dev.01165

53 Linggi, M. S. et al. Neurotrophin receptor interacting factor (NRIF) is an essential mediator of apoptotic signaling by the p75 neurotrophin receptor. J Biol Chem 280, 13801‒13808 (2005). 10.1074/jbc.M410435200

54 Kuwabara, T. et al. Insulin biosynthesis in neuronal progenitors derived from adult hippocampus and the olfactory bulb. EMBO Mol Med 3, 742‒754 (2011). 10.1002/emmm.201100177

55 Hori, Y., Gu, X., Xie, X. & Kim, S. K. Differentiation of insulin-producing cells from human neural progenitor cells. PLoS Med 2, e103 (2005). 10.1371/journal.pmed.0020103

56 Yang, W. et al. ERK1/2-dependent phosphorylation and nuclear translocation of PKM2 promotes the Warburg effect. Nat Cell Biol 14, 1295‒1304 (2012). 10.1038/ncb2629

57 Israelsen, W. J. & Vander Heiden, M. G. Pyruvate kinase: Function, regulation and role in cancer. Semin Cell Dev Biol 43, 43‒51 (2015). 10.1016/j.semcdb.2015.08.004

58 Osato, N. & Hamada, M. Systematic discovery of directional regulatory motifs associated with human insulator sites. bioRxiv (2025).

59 Barrett, T. et al. NCBI GEO: archive for functional genomics data sets--update. Nucleic acids research 41, D991‒995 (2013). 10.1093/nar/gks1193

60 Edgar, R., Domrachev, M. & Lash, A. E. Gene Expression Omnibus: NCBI gene expression and hybridization array data repository. Nucleic acids research 30, 207‒210 (2002). 10.1093/nar/30.1.207

61 Kolmykov, S. et al. GTRD: an integrated view of transcription regulation. Nucleic acids research 49, D104‒D111 (2021). 10.1093/nar/gkaa1057

62 Hinrichs, A. S. et al. The UCSC Genome Browser Database: update 2006. Nucleic acids research 34, D590‒598 (2006). 10.1093/nar/gkj144

63 Mudge, J. M. et al. GENCODE 2025: reference gene annotation for human and mouse. Nucleic acids research 53, D966‒D975 (2025). 10.1093/nar/gkae1078

64 Van Nostrand, E. L. et al. A large-scale binding and functional map of human RNA-binding proteins. Nature 583, 711‒719 (2020). 10.1038/s41586-020-2077-3

65 Hanzelmann, S., Castelo, R. & Guinney, J. GSVA: gene set variation analysis for microarray and RNA-seq data. BMC Bioinformatics 14, 7 (2013). 10.1186/1471-2105-14-7

66 Aibar, S. et al. SCENIC: single-cell regulatory network inference and clustering. Nature methods 14, 1083‒1086 (2017). 10.1038/nmeth.4463

