## Supplementary Figures S1-S8 for "Pathway redistribution across cellular states reveals a shared signaling backbone and context-dependent regulatory modules in RNA-binding protein networks"

**a.**

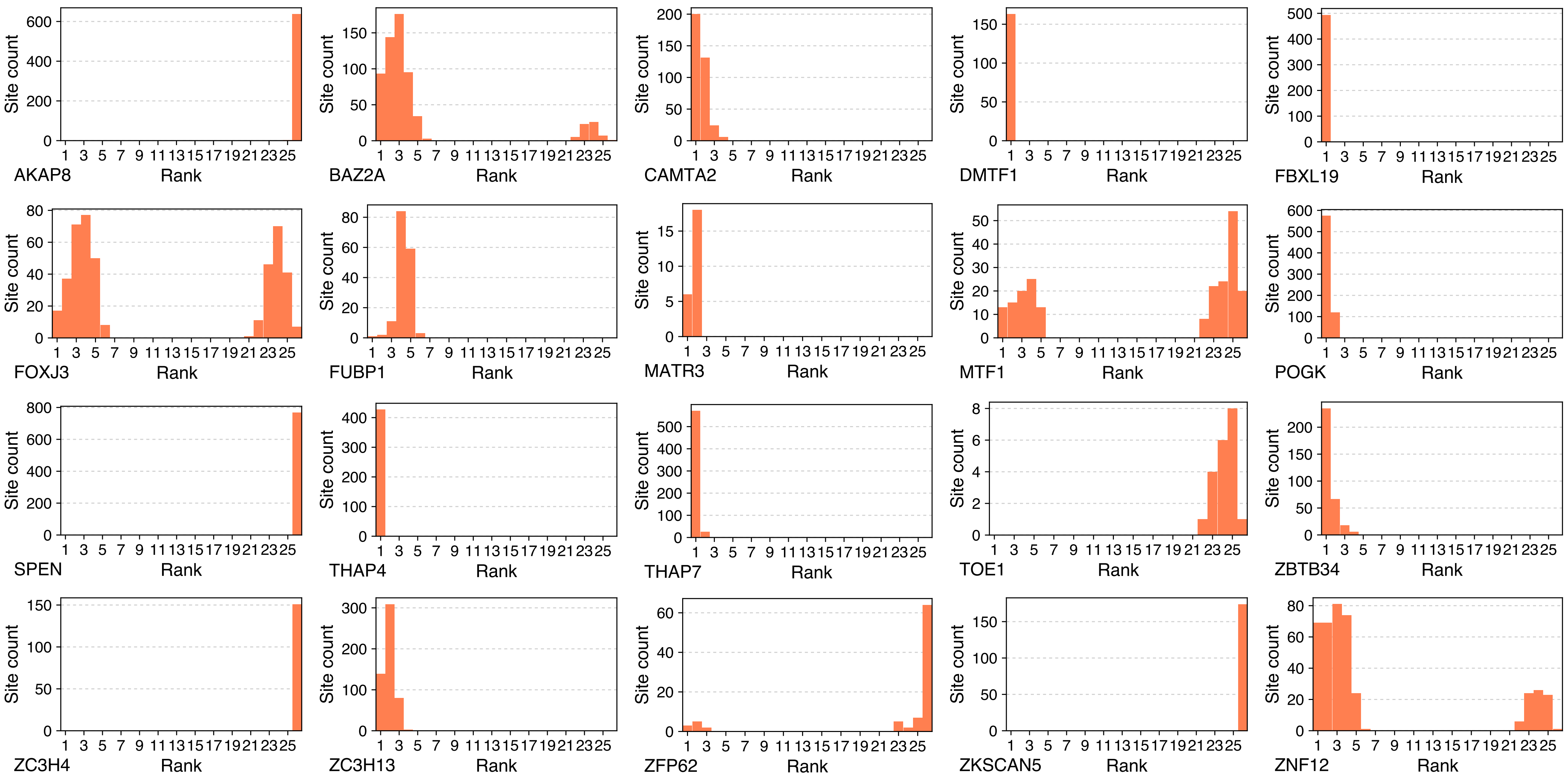
AKAP8
BAZ2A
CAMTA2
DMTF1
FBXL19
FOXJ3
FUBP1
MATR3
MTF1
POGK
SPEN
THAP4
THAP7
TOE1
ZBTB34
ZC3H4
ZC3H13
ZFP62
ZKSCAN5
ZNF12
Site count
Rank

**a.**

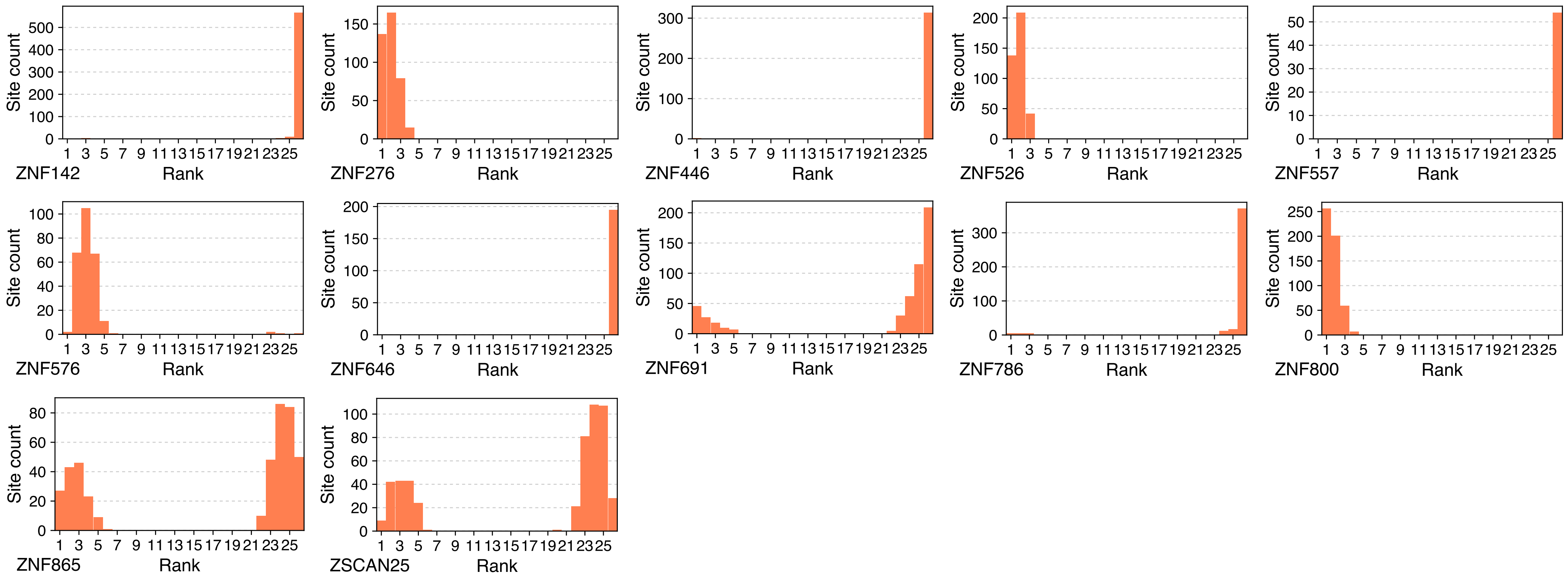
Site count
Rank
ZNF142
ZNF276
ZNF446
ZNF526
ZNF557
ZNF576
ZNF646
ZNF691
ZNF786
ZNF800
ZNF865
ZSCAN25

**b.**

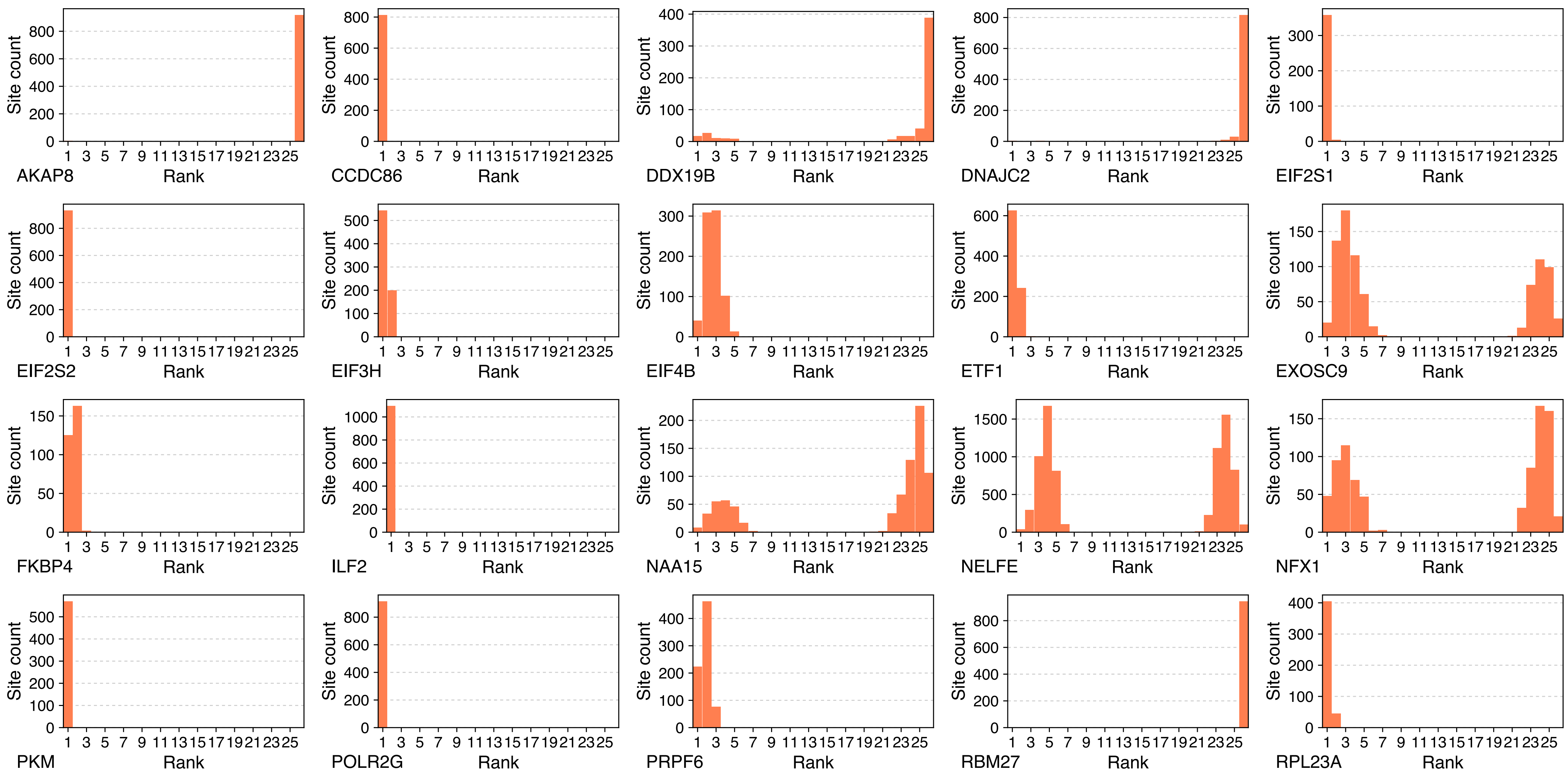
Site count
Rank
AKAP8
CCDC86
DDX19B
DNAJC2
EIF2S1
EIF2S2
EIF3H
EIF4B
ETF1
EXOSC9
FKBP4
ILF2
NAA15
NELFE
NFX1
PKM
POLR2G
PRPF6
RBM27
RPL23A

**b.**

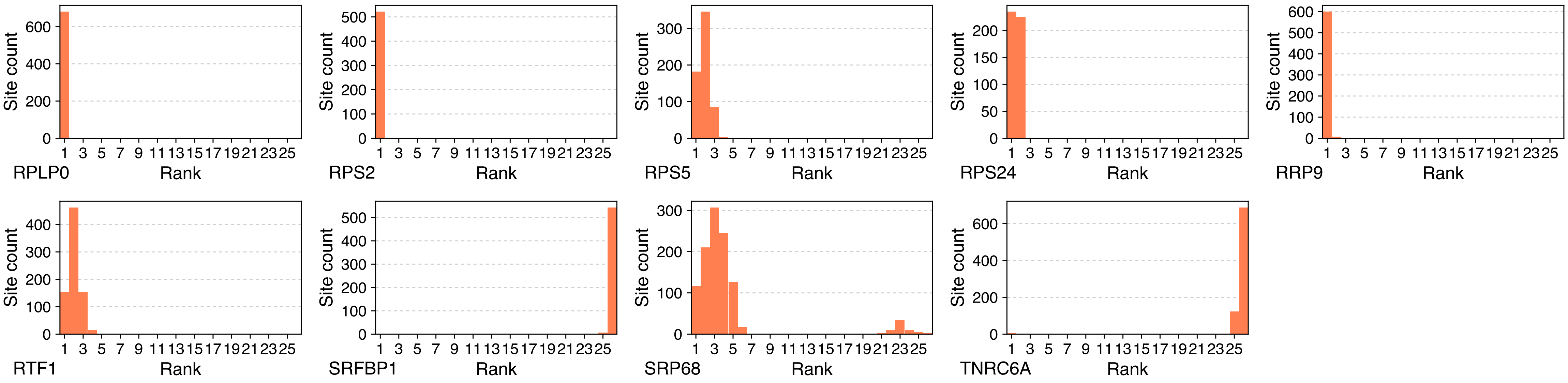


**Supplementary Figure S1. Rank distribution of DeepLIFT contribution scores using a reduced background configuration.**

The rank distribution of DeepLIFT contribution scores is shown for predicted nucleic acid-binding protein (NABP) regulatory targets with experimentally validated binding evidence, identified by ChIP-seq (DNA-binding proteins) or eCLIP (RNA-binding proteins). Contribution scores were computed using a reduced DeepLIFT background reference (input $\times$ 0.5).

NABPs capable of binding both DNA and RNA are classified according to the experimental assay used for validation. The x-axis represents rank bins, where rank 1 corresponds to positions 1–50, rank 2 to ranks 51–100, and rank 26 to ranks 1,251–1,310 among all candidate NABPs.

**a.** DNA-binding proteins; **b**. RNA-binding proteins.

† For example, AKAP8 is known to interact with both DNA and RNA, but is shown here according to the assay (ChIP-seq or eCLIP) from which its binding sites were derived. ChIP-seq data were obtained from HepG2 cells, and eCLIP data from K562 cells.

a.

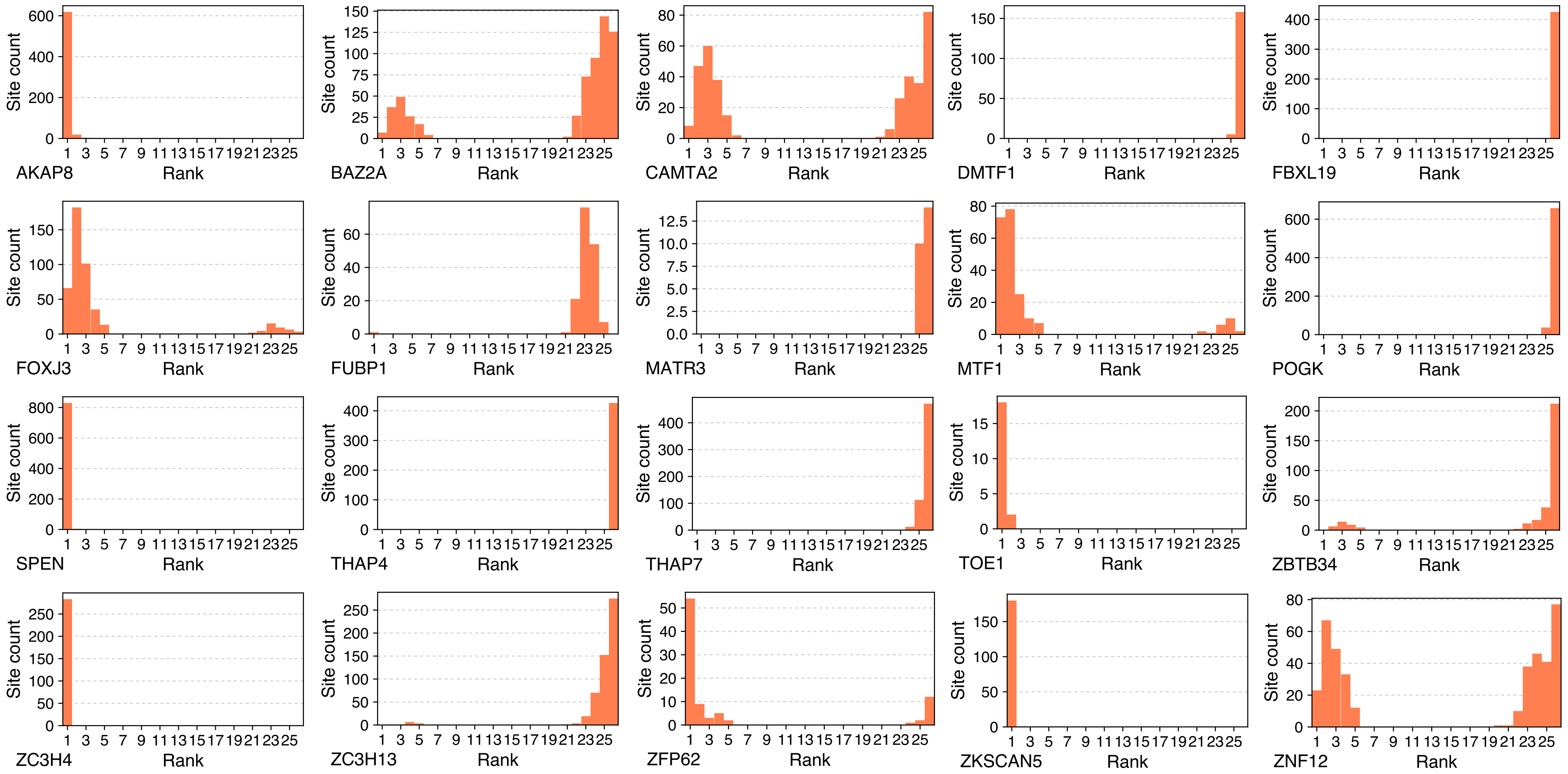
AKAP8
BAZ2A
CAMTA2
DMTF1
FBXL19
FOXJ3
FUBP1
MATR3
MTF1
POGK
SPEN
THAP4
THAP7
TOE1
ZBTB34
ZC3H4
ZC3H13
ZFP62
ZKSCAN5
ZNF12
Site count
Rank

a.

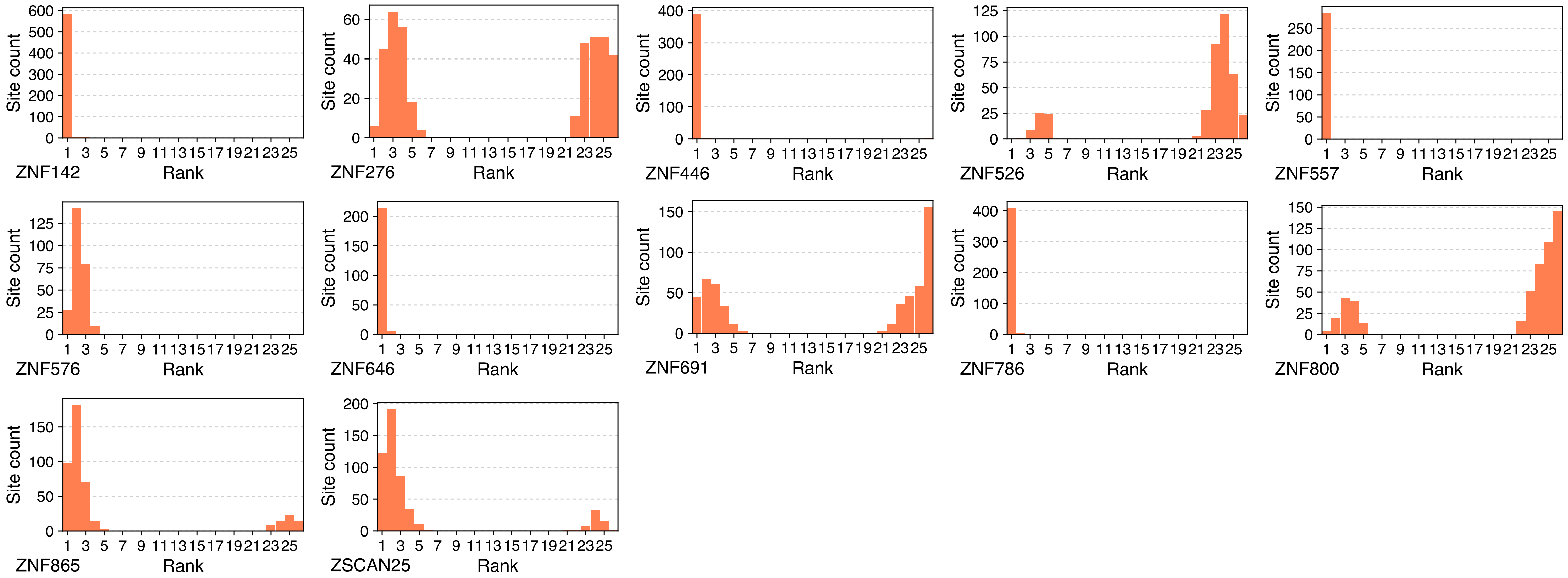
ZNF142
ZNF276
ZNF446
ZNF526
ZNF557
ZNF576
ZNF646
ZNF691
ZNF786
ZNF800
ZNF865
ZSCAN25
Site count
Rank

**b.**

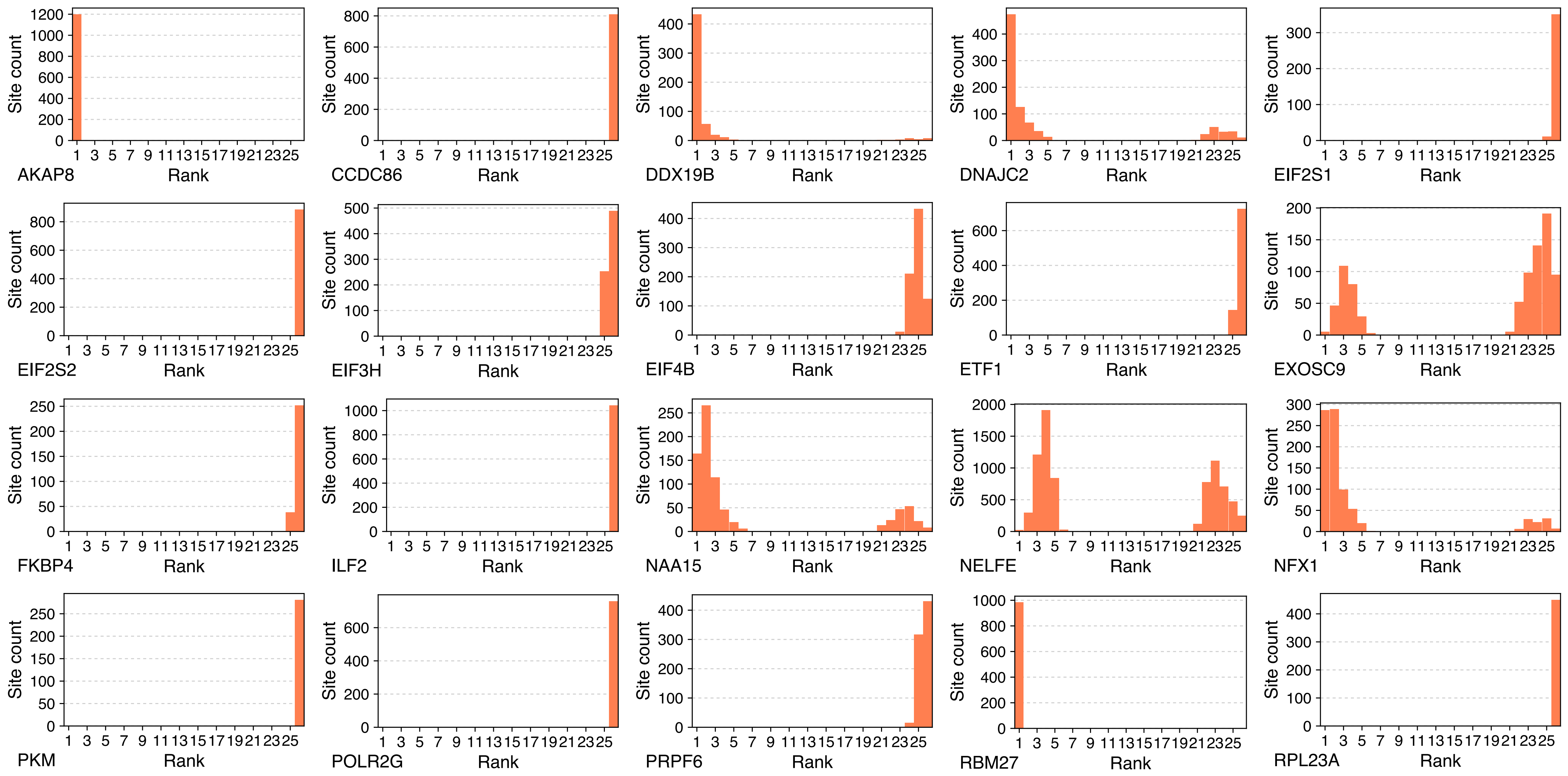
Site count
Rank
AKAP8
CCDC86
DDX19B
DNAJC2
EIF2S1
EIF2S2
EIF3H
EIF4B
ETF1
EXOSC9
FKBP4
ILF2
NAA15
NELFE
NFX1
PKM
POLR2G
PRPF6
RBM27
RPL23A

**b.**

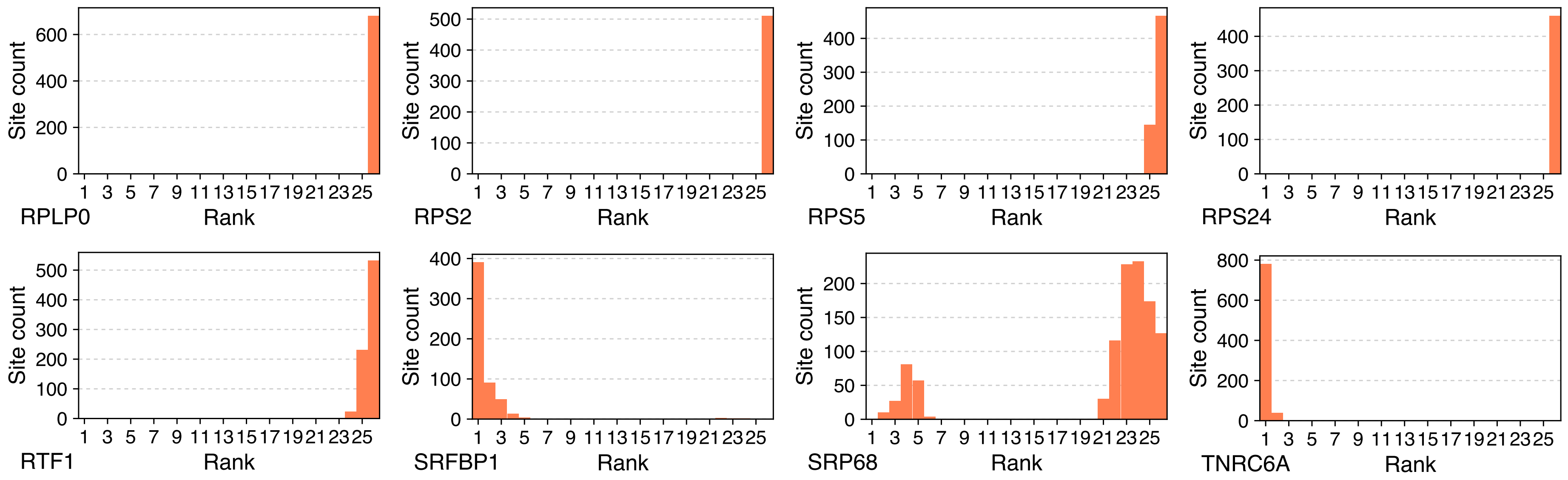


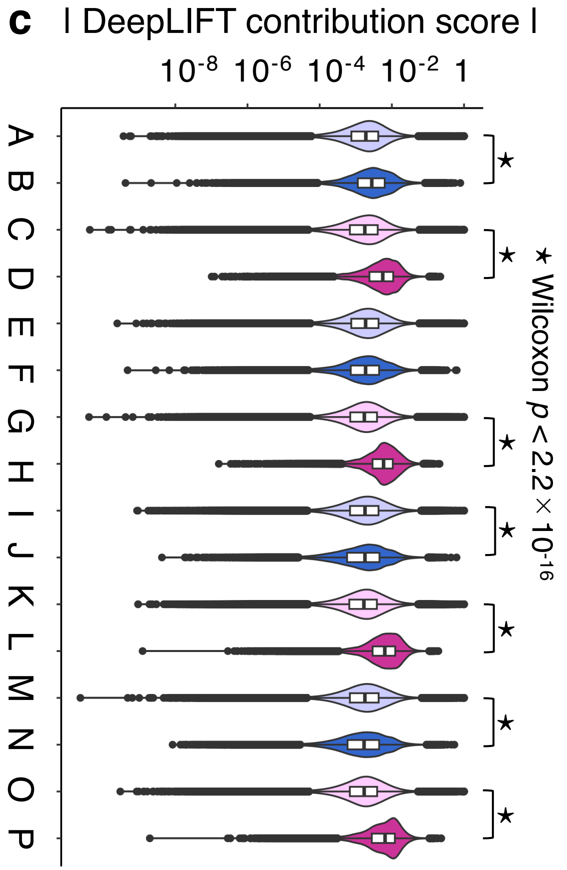


**Supplementary Figure S2. Effects of an amplified DeepLIFT background on contribution score distributions. Rank distribution of DeepLIFT contribution scores using an amplified background.**

**a. b**. Rank distributions of DeepLIFT contribution scores for predicted nucleic acid-binding protein (NABP) regulatory targets using an amplified background reference (input × 2). As in **Supplementary Figure S1**, genes with experimentally validated binding sites identified by ChIP-seq (DNA-binding proteins) or eCLIP (RNA-binding proteins) are preferentially enriched at extreme ranks (high or low). Although background scaling alters the polarity of contribution scores relative to the reduced-background condition, the enrichment of experimentally validated targets at extreme ranks is preserved. **a.** DNA-binding proteins; **b**. RNA-binding proteins.

**c**. Violin plots showing absolute DeepLIFT contribution scores computed using the amplified background (input × 2). The same comparison strategy and labeling scheme as in **Figure 2d** were applied, contrasting co-expression–derived predicted binding sites with corresponding ChIP-seq–derived sites not included in those sets. Distribution patterns closely resemble those observed under the reduced-background condition (**Figure 2d**), indicating that relative enrichment of contribution score is robust to background scaling.

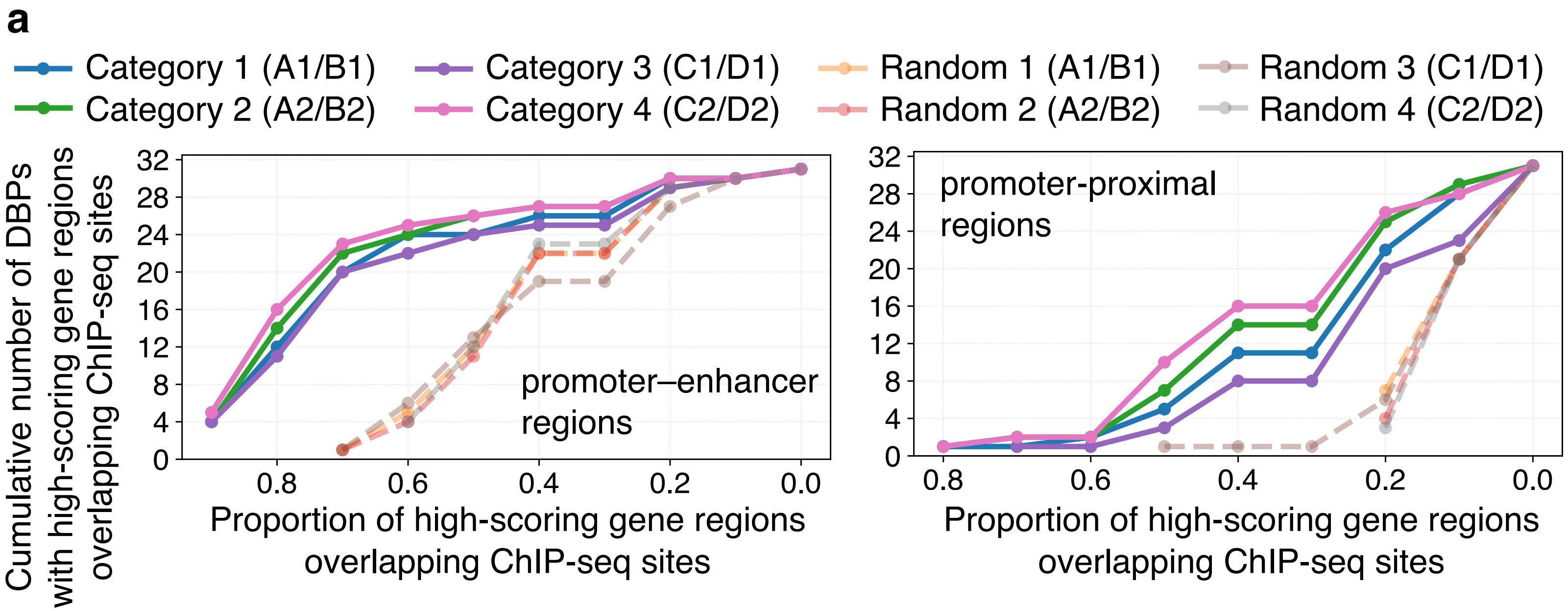


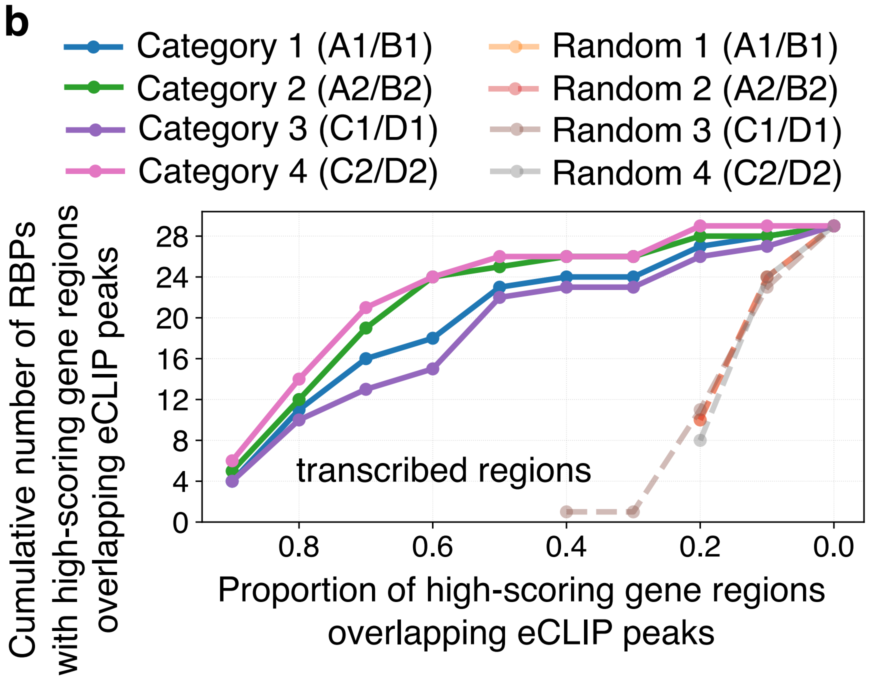


**Supplementary Figure S3. Robustness of overlap between high-contribution regions and experimental binding sites under an amplified DeepLIFT background.**

Analyses equivalent to those shown in **Figure 3b** and **3c** were performed using an amplified DeepLIFT background reference (input × 2). The proportions of high-contribution regions overlapping ChIP-seq binding sites in promoter–enhancer (±100 kb) and promoter-proximal (±5 kb) regions (**a**), as well as eCLIP peaks within transcribed regions (**b**), were calculated using the same rank-based and score-based thresholds as in the main figure.

Across all threshold definitions, overlap patterns closely mirrored those observed under the reduced-background condition (**Figure 3**), indicating that the concordance between model-predicted regulatory targets and experimentally validated binding sites is robust to substantial changes in DeepLIFT background definition.

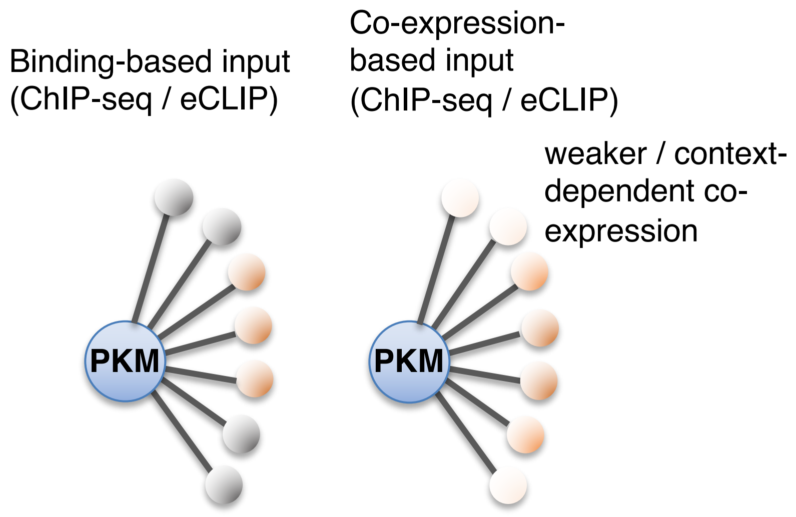


**Supplementary Figure S4. Conceptual model illustrating how input feature definition and background scaling influence DeepLIFT contribution score patterns.**

This schematic illustrates conceptual differences between binding-based and co-expression–based input features used to infer regulatory targets of nucleic acid-binding proteins (NABPs), exemplified here using PKM.

Left, when experimental binding data (ChIP-seq or eCLIP peaks) are used as model inputs, the set of candidate target genes includes both functionally regulated genes (orange nodes) and genes that are bound but do not directly contribute to expression regulation (gray nodes). As a result, DeepLIFT contribution scores are selectively elevated for functionally relevant targets, whereas many bound but non-regulatory genes receive low or negligible scores.

Right, when co-expression–based inputs are used, candidate target genes are preselected based on transcriptomic correlation with PKM expression. Consequently, most input genes are expected to participate in PKM-associated regulatory programs (orange nodes). However, variation in co-expression strength across cellular contexts means that genes with weaker or context-dependent associations (light orange nodes) contribute less to expression prediction, resulting in lower contribution scores.

Together, differences in input composition and background reference can lead to distinct—and in some cases inverted—DeepLIFT contribution score patterns between binding-based and co-expression–based analyses. Importantly, in both settings, genes with higher contribution scores are enriched for experimentally supported targets and biologically relevant pathways, indicating that DeepLIFT captures relative regulatory importance rather than direct binding or absolute expression levels.

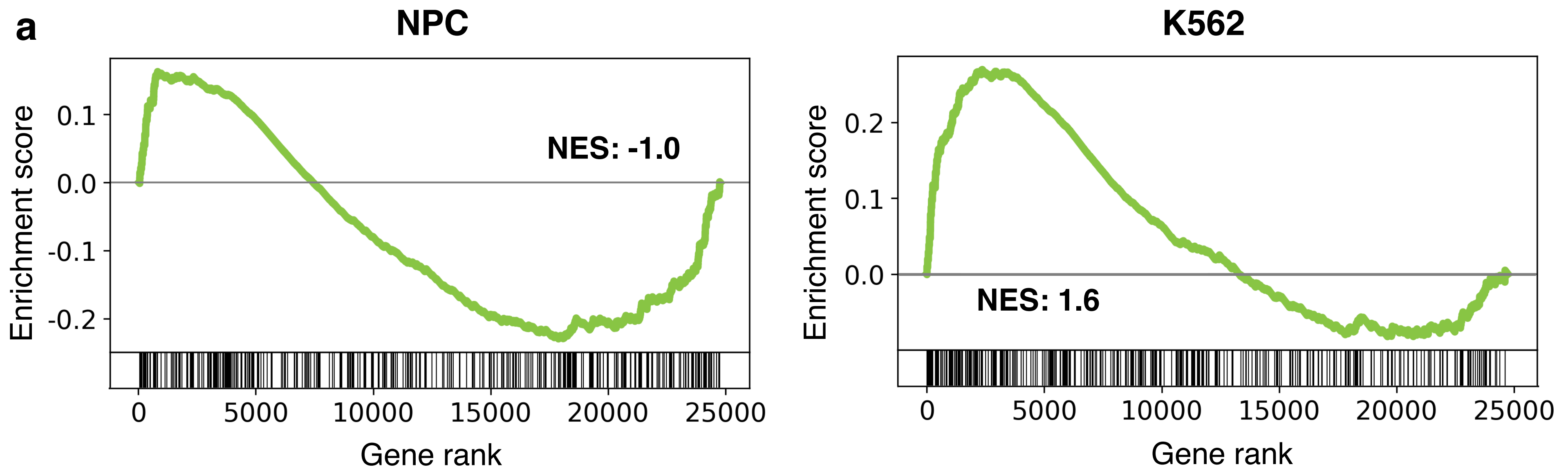


**Neuronal System (ΔNES = 2.6)**

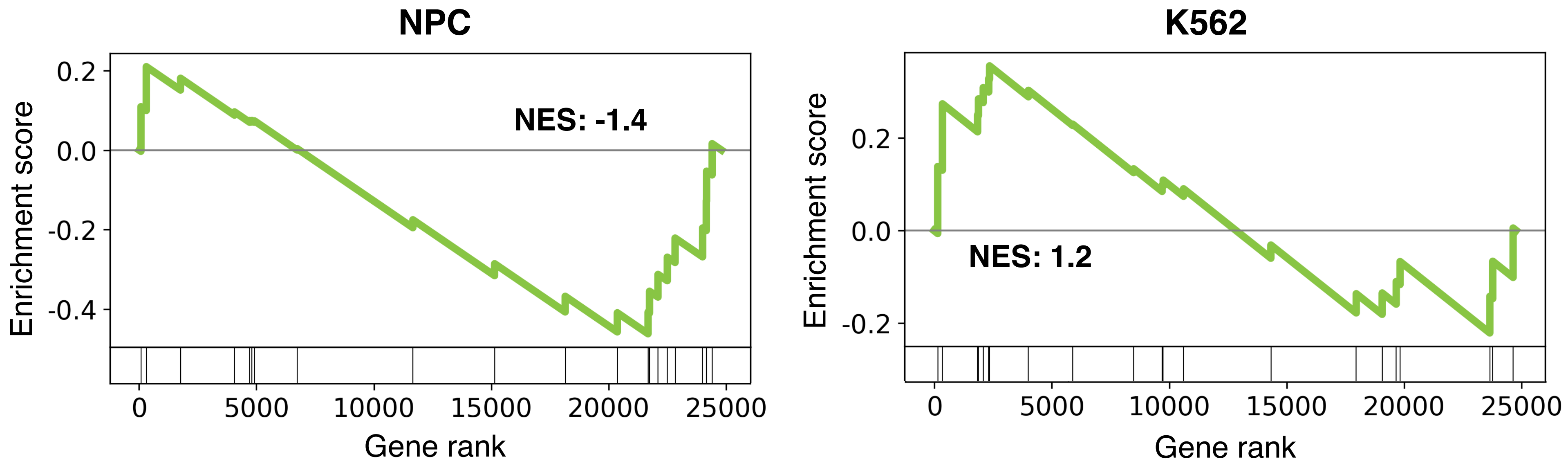


**Synaptic adhesion-like molecules (ΔNES = 2.6)**

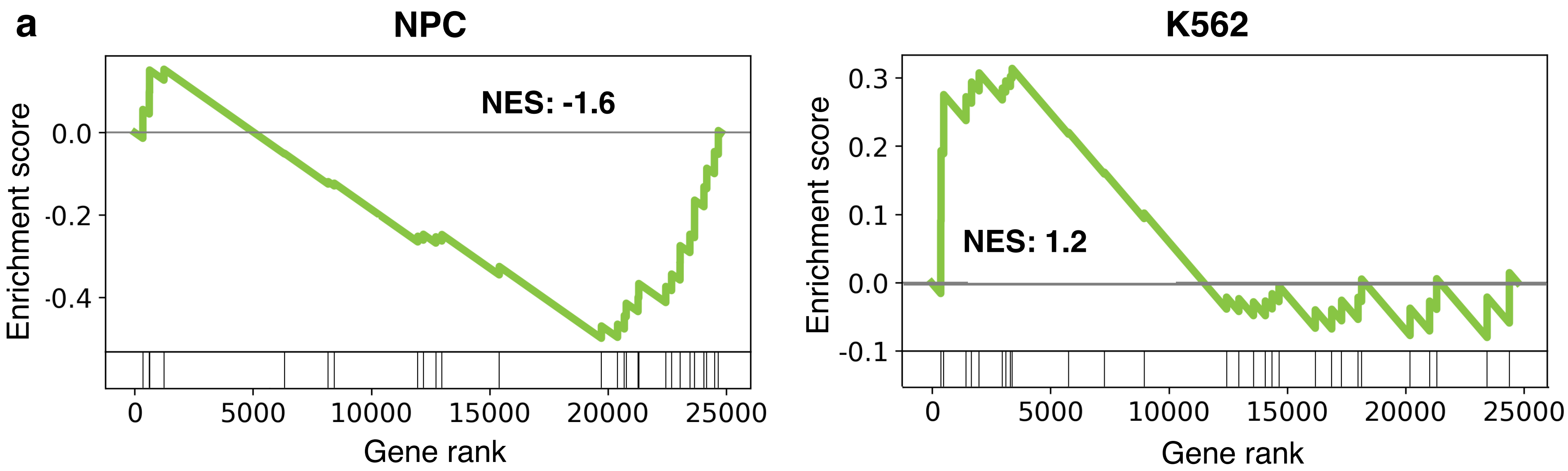


**Glycosaminoglycan-protein linkage region biosynthesis (ΔNES = 2.8)**

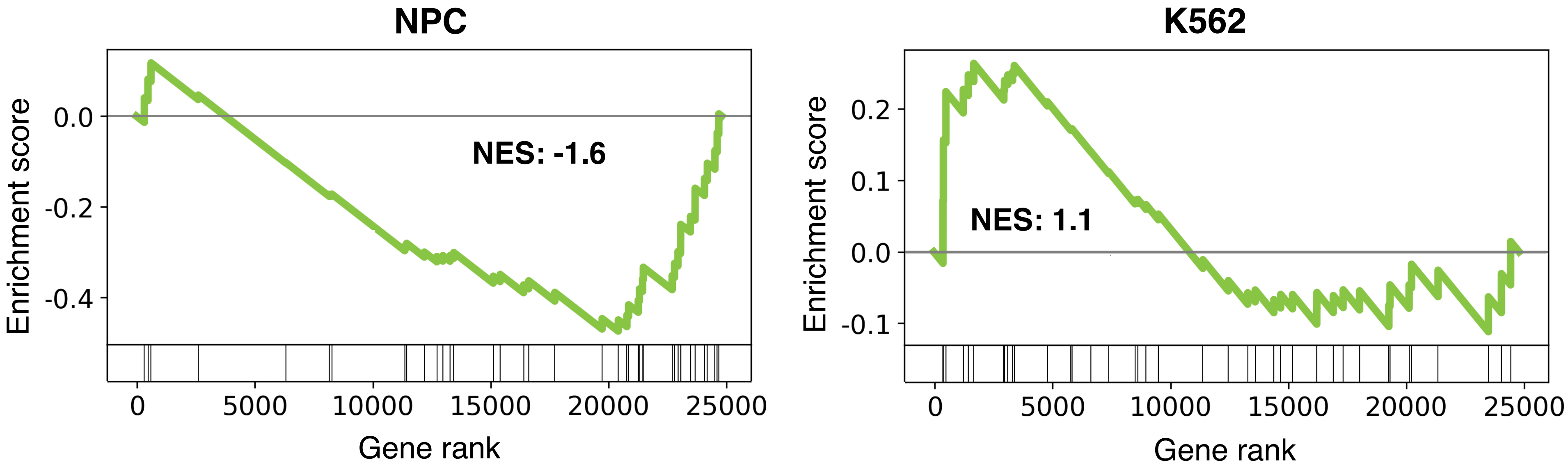


**Diseases associated with glycosaminoglycan metabolism (ΔNES = 2.7)**

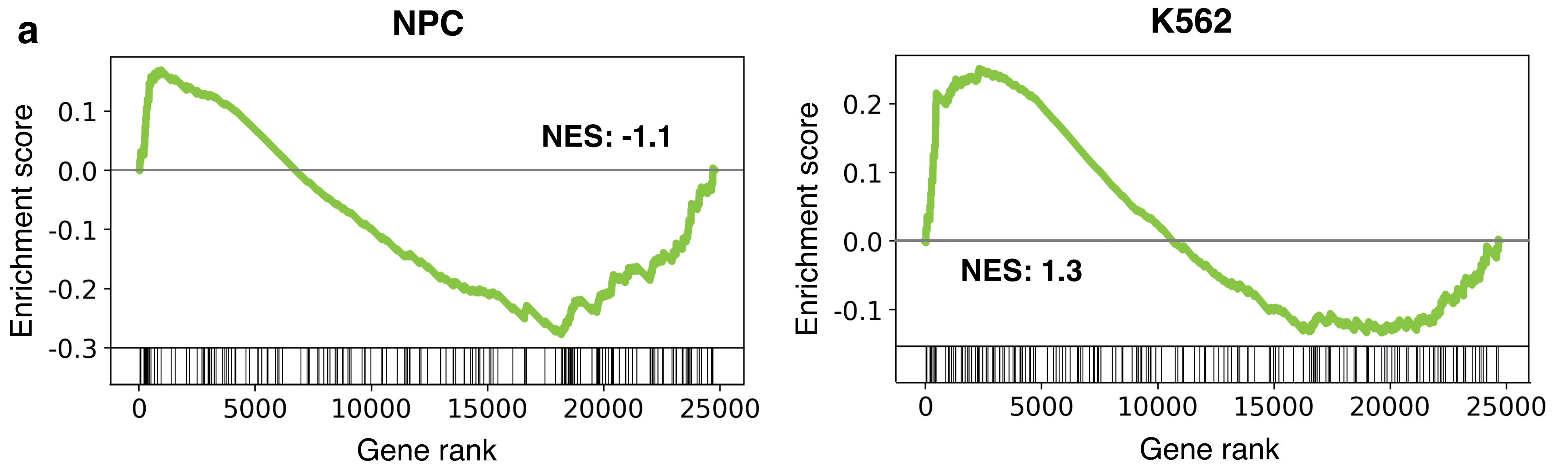


**Ion channel transport (ΔNES = 2.4)**

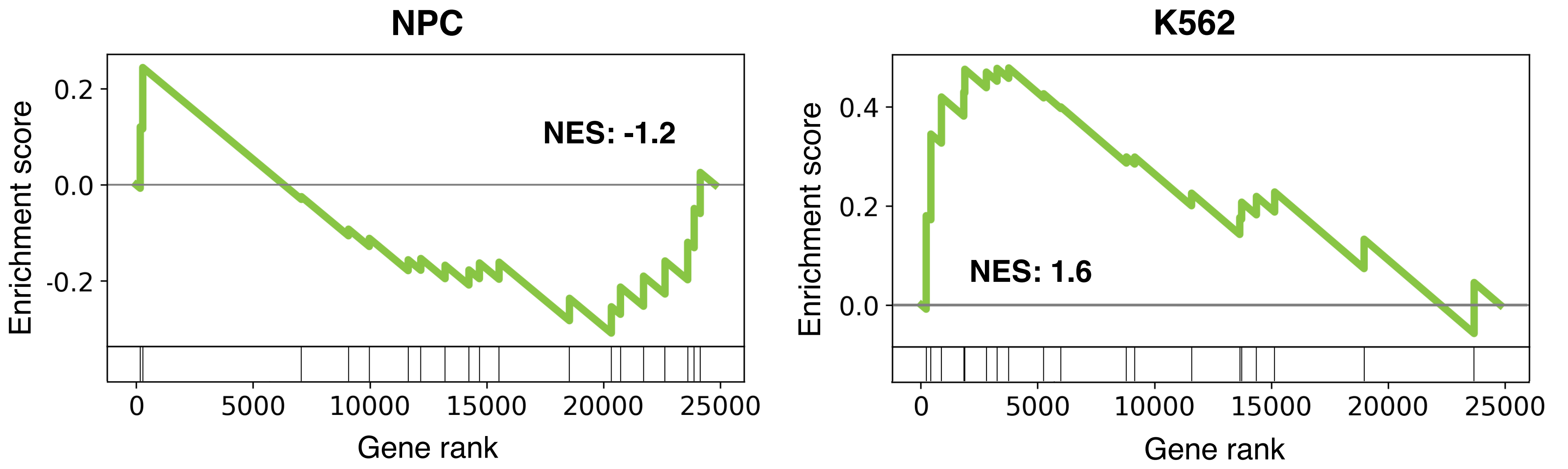


**Receptor-type tyrosine-protein phosphatases (ΔNES = 2.5)**

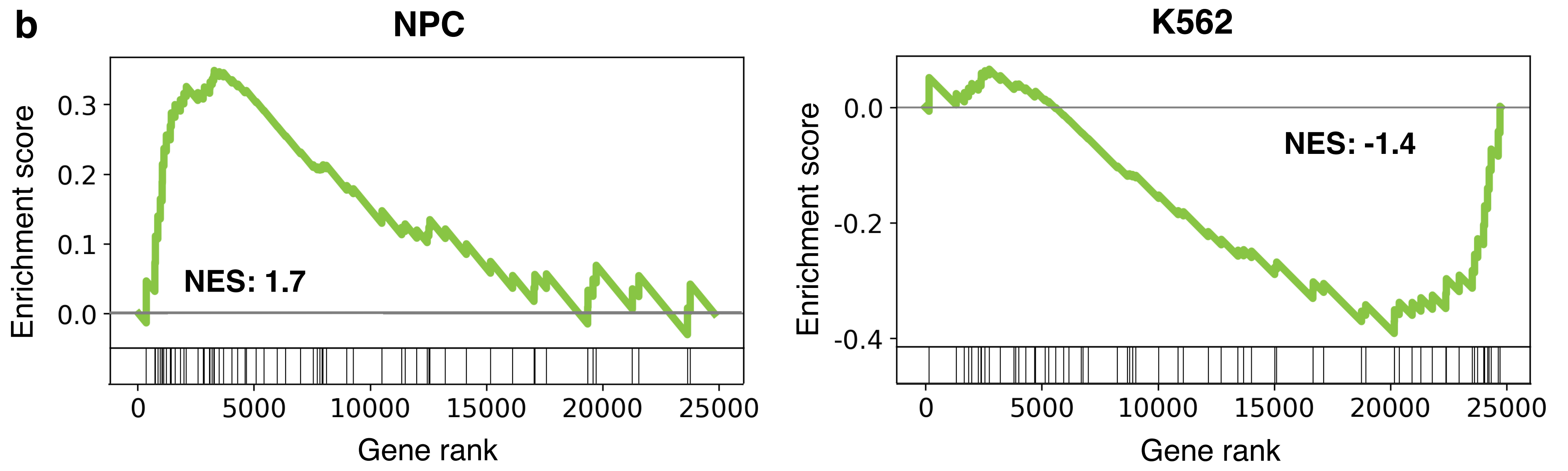


**Scavenging of heme from plasma (ΔNES = -3.1)**

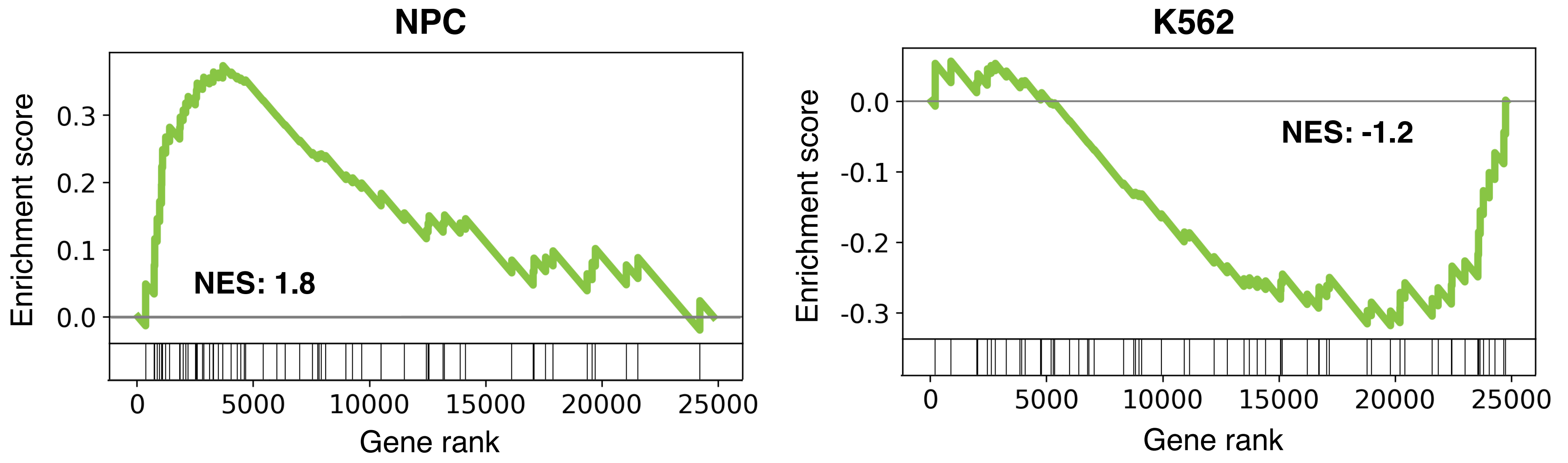


**FCGR activation (ΔNES = -3.0)**

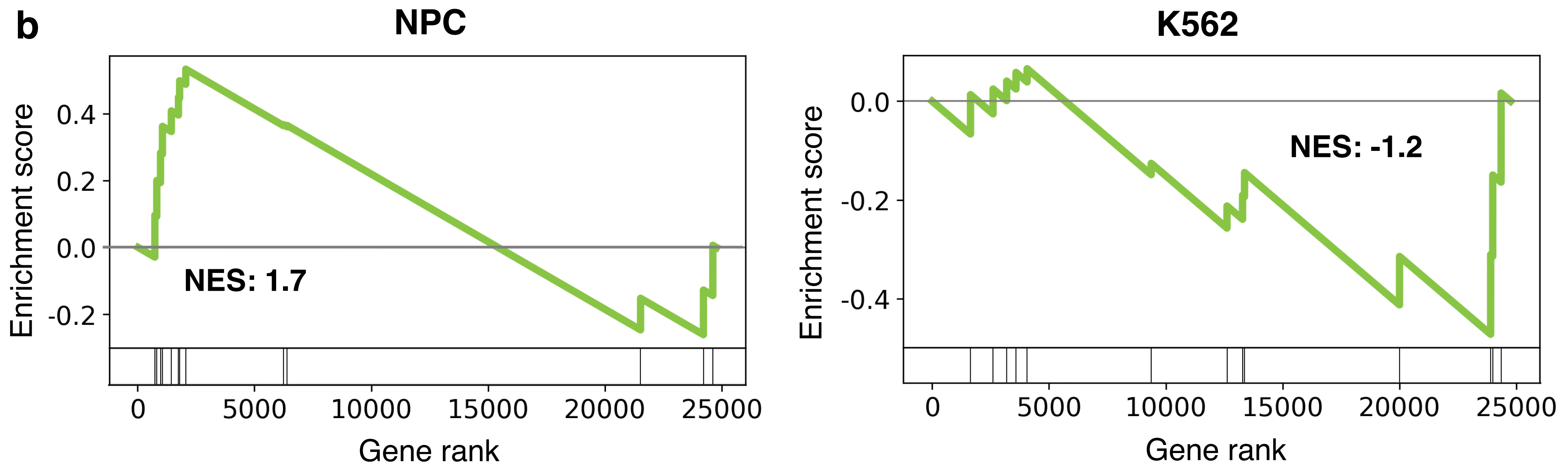


**Response to metal ions (ΔNES = -2.9)**

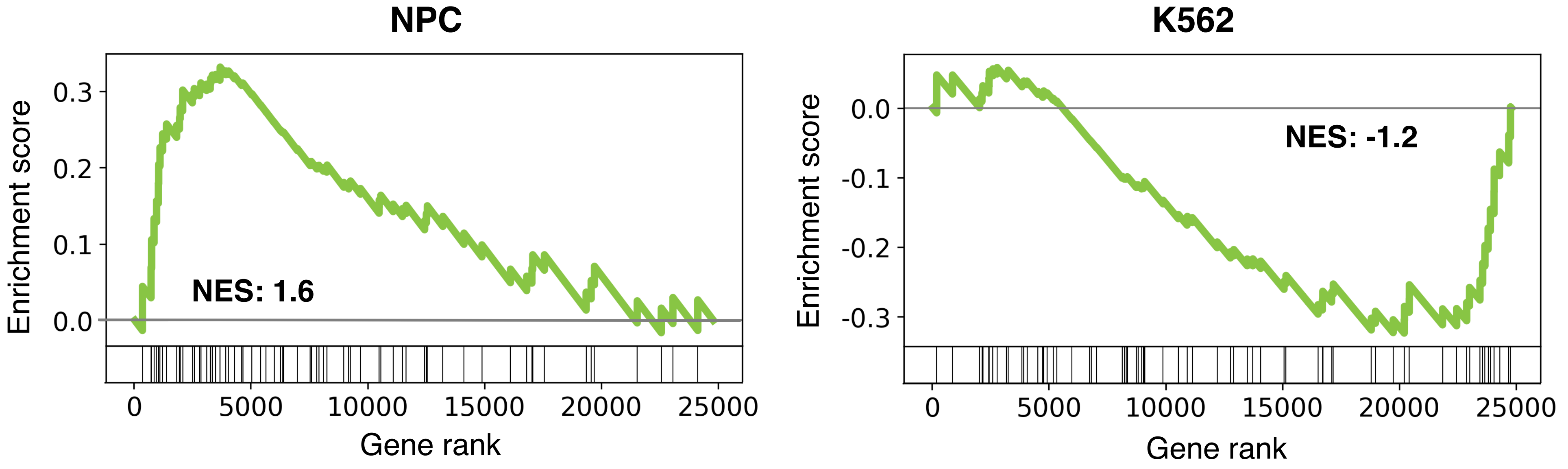


**Initial triggering of complement (ΔNES = -2.8)**

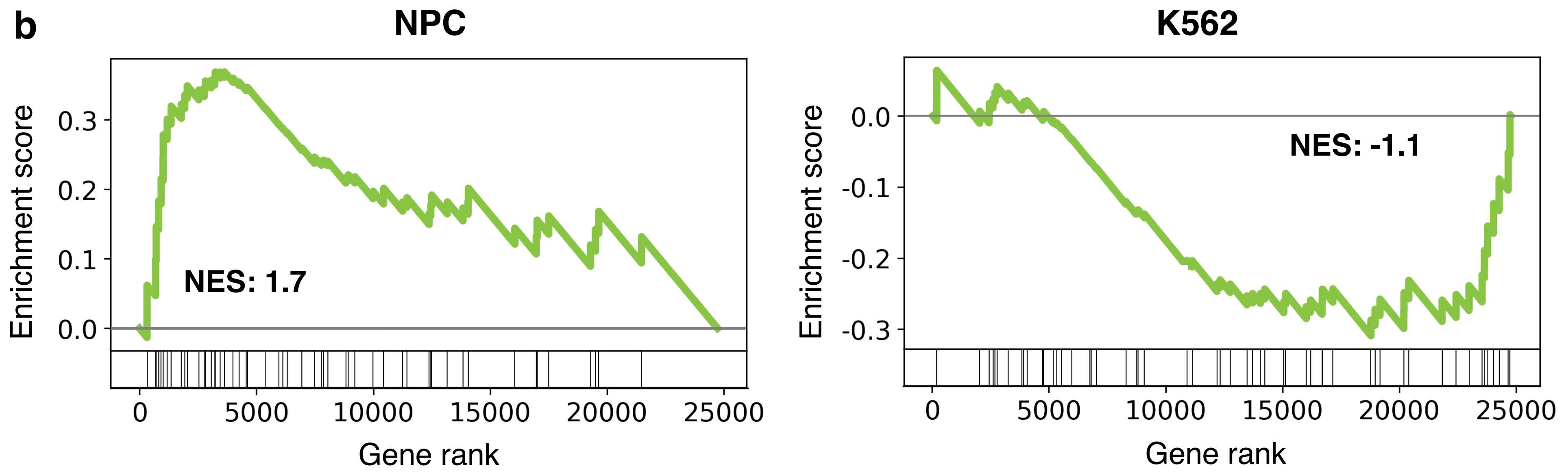


**CD22 mediated BCR regulation (ΔNES = -2.8)**

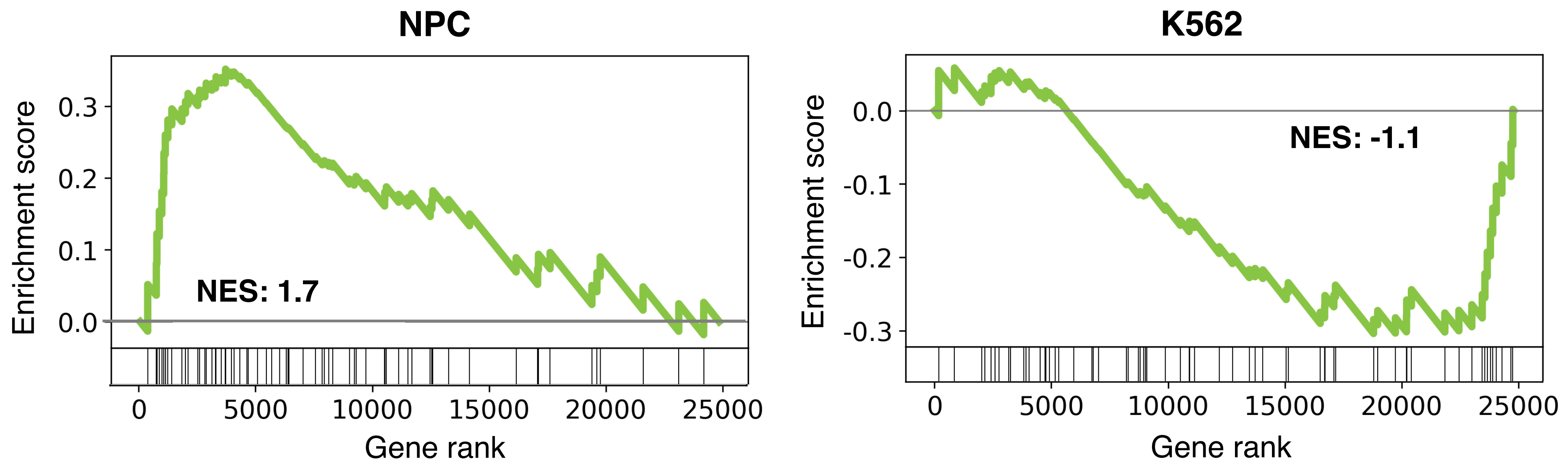


**Creation of C4 and C2 activators (ΔNES = -2.8)**

**Supplementary Figure S5. Representative GSEA plots illustrating ΔNES-based pathway redistribution.**

Representative examples of Reactome pathways showing contrasting enrichment patterns between neural progenitor cell (NPC) and K562 leukemia contexts are shown. Pathways associated with the Neural System are preferentially enriched among positive ΔNES values, whereas immune-associated pathways are more frequently observed among negative ΔNES values, These patterns reflect context-dependent redistribution of pathway-associated genes along contribution-ranked lists rather than direct pathway activation or repression.

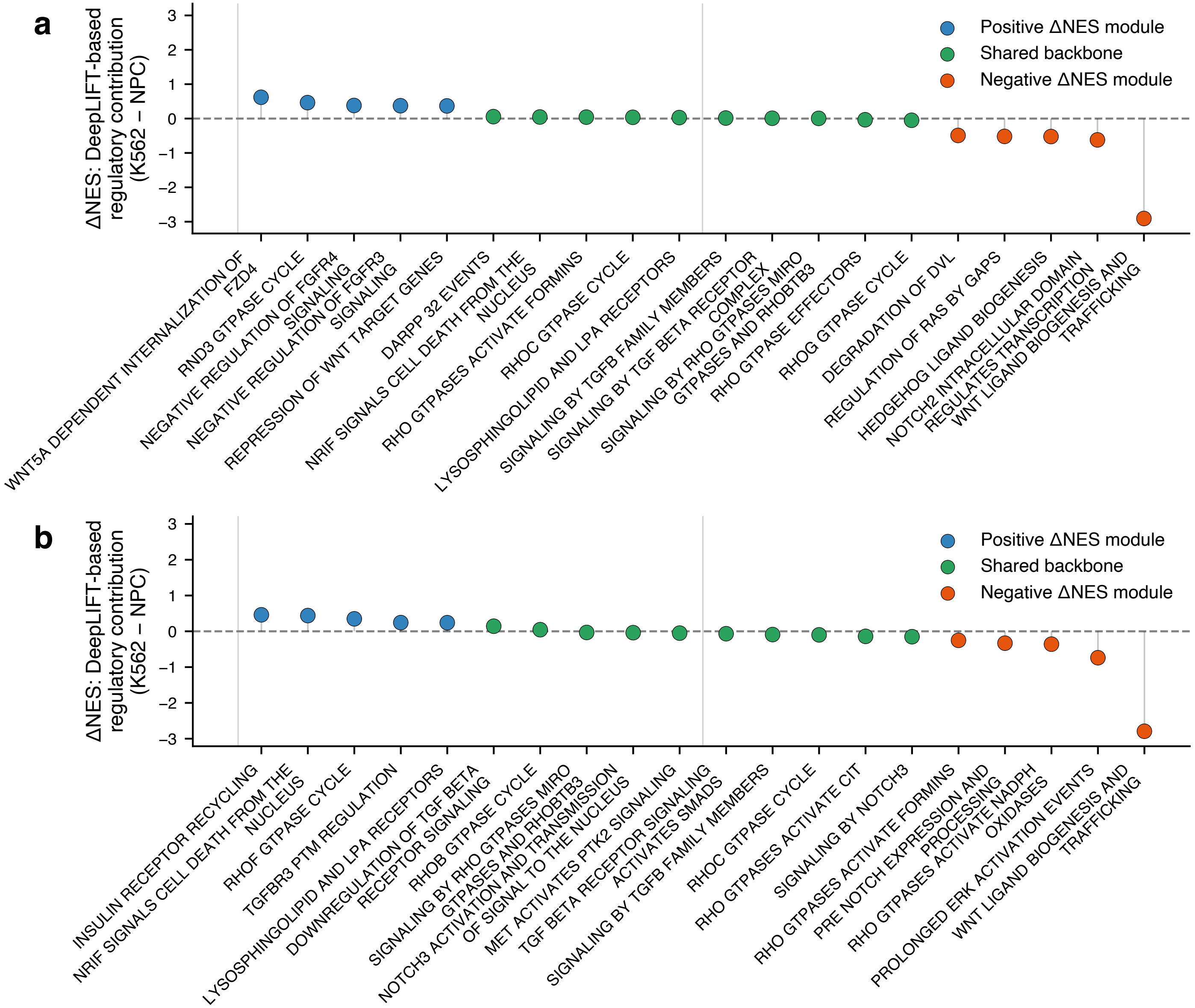


**Supplementary Figure S6. Signal Transduction pathways along the ΔNES axis for HNRNPK (a) and NELFE (b).** ΔNES values were derived from DeepLIFT-based GSEA using protein-specific contribution score rankings in K562 and neural progenitor cell (NPC) contexts. Shared signaling backbone pathways were defined as strongly enriched Signal Transduction pathways with near-zero ΔNES values (max |NES| ≥ 1.5), indicating conserved regulatory contributions across cellular contexts. In contrast, context-specific signaling pathways were defined as strongly enriched pathways with large absolute ΔNES values, reflecting cell type–dependent redistribution of inferred regulatory influence. For each protein, ten representative shared backbone pathways and five representative context-specific pathways from both NPC- and K562-associated regulatory modules were selected based on ΔNES ranking and enrichment magnitude.

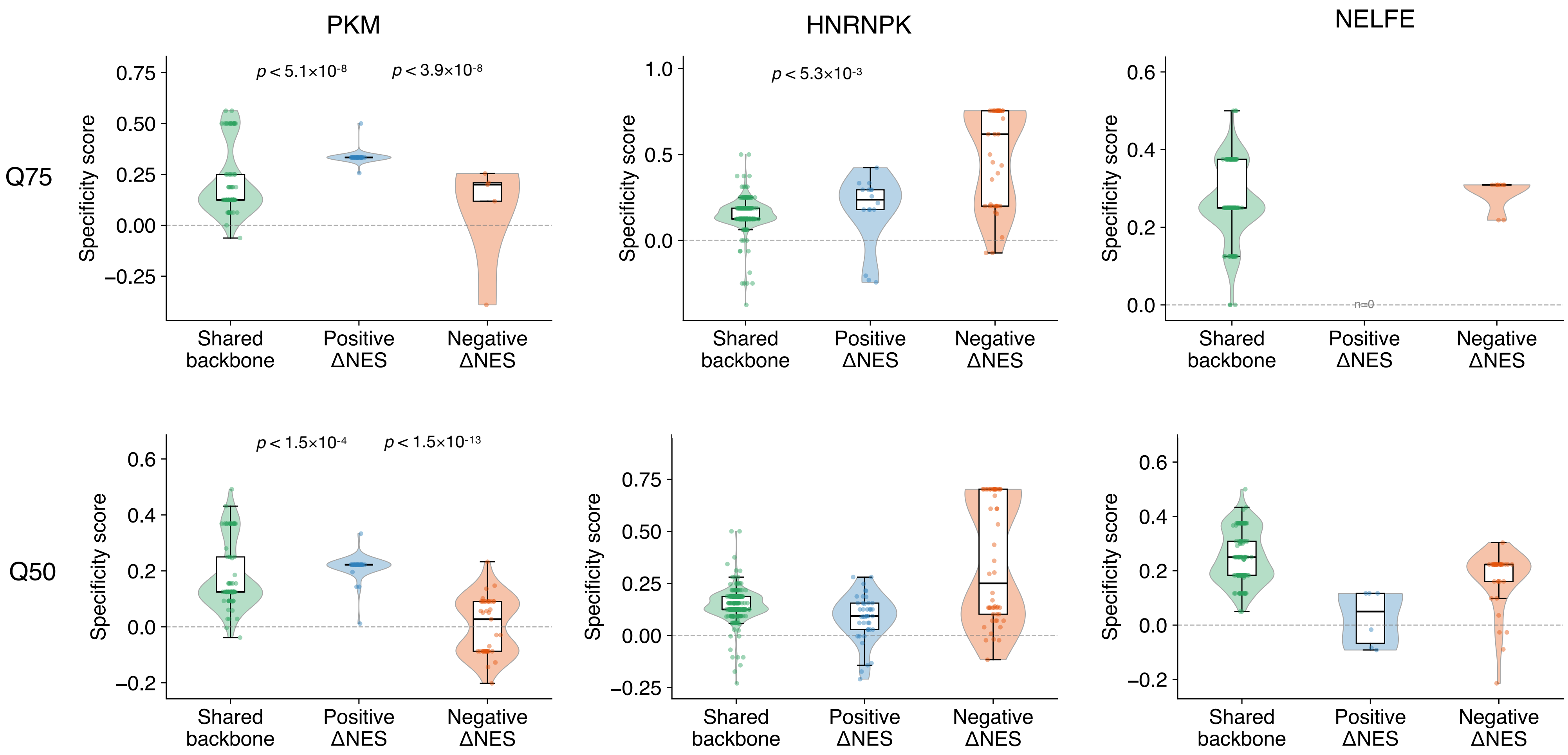


**Supplementary Figure S7. Conserved shared signaling backbone and context-dependent regulatory redistribution across RNA-binding proteins.** Signal Transduction pathways were classified into shared backbone pathways (lowest IΔNESI quartile) and context-dependent modules defined by positive or negative ΔNES values (Q75 in upper panels; Q50 in lower panels). Leading-edge genes identified by DeepLIFT-based GSEA were assigned specificity scores based on their relative enrichment within each signaling module compared with other Signal Transduction pathways. Higher specificity scores indicate preferential concentration of genes within module-specific pathways. Violin plots show the distributions of specificity scores for PKM, HNRNPK, and NELFE. Across RBPs, shared backbone pathways exhibited relatively stable specificity distributions, whereas context-dependent modules showed protein-specific shifts, consistent with selective redistribution of inferred regulatory influence. Statistical significance was assessed using two-sided Mann–Whitney U tests.

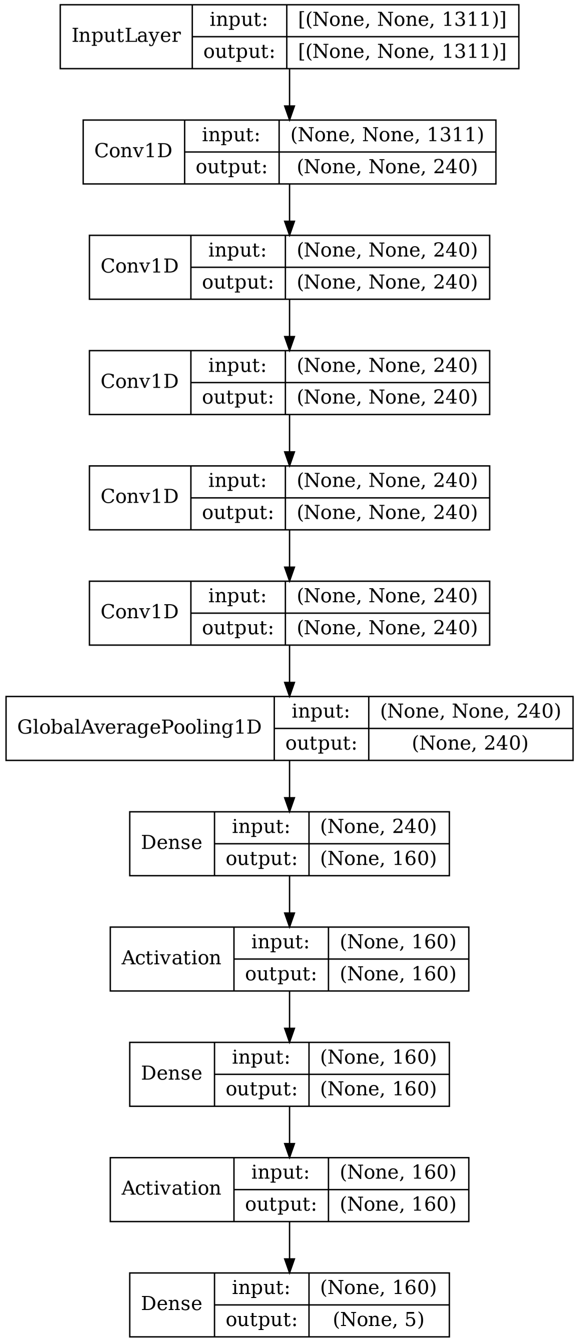


**Supplementary Figure S8. Structure of deep neural network.**
