## Supplementary Note and Tables for "Pathway redistribution across cellular states reveals a shared signaling backbone and context-dependent regulatory modules in RNA-binding protein networks"

### Supplementary Note for Figure 3. Definition of contribution score–based regions, thresholds, and enrichment analyses

#### Contribution score ranking and binning

For each nucleic acid-binding protein (NABP), DeepLIFT contribution scores were computed for all predicted gene-level regulatory regions and ranked from highest positive to lowest negative values. Ranked scores were partitioned into consecutive bins of 50 ranks, yielding 26 bins per NABP (rank 1: 1–50; rank 2: 51–100; …; rank 26: 1,251–1,310), unless otherwise specified.

#### Definition of high- and low-contribution regions

High- and low-contribution regions were defined using both rank-based and score-based thresholds to assess the robustness of enrichment results. Rank-based thresholds included upper and lower rank cutoffs (e.g., top or bottom 20 and 50 ranks), whereas score-based thresholds were defined using absolute contribution score cutoffs (e.g., ≥ 0.01 or ≥ 0.001). All threshold combinations used in the analyses are summarized in **Supplementary Tables S2** and **S3.**

#### Genomic regions used for overlap analyses

For DNA-binding proteins (DBPs), ChIP-seq overlap analyses were conducted in two genomic contexts: promoter-proximal regions (±5 kb from the transcription start site [TSS]) and extended promoter–enhancer regions (±100 kb from the TSS). For RNA-binding proteins (RBPs), eCLIP overlap analyses were restricted to transcribed regions spanning from the TSS to the transcription end site (TES), including both exonic and intronic intervals.

### Random controls

Random control regions were generated by sampling genomic intervals matched for size and genomic context while excluding annotated binding sites. These controls were used to estimate expected overlap by chance. All enrichment and overlap analyses were performed in parallel using these matched random controls.

### Background configuration for DeepLIFT analyses

Unless otherwise specified, DeepLIFT contribution scores were computed using a reduced background reference (input × 0.5). Analyses using an alternative amplified background (input × 2) produced qualitatively consistent results and are presented in **Supplementary Figures S2** and **S3.**

### Cell type configurations used for model training and comparative analyses

To accommodate differences in data availability and analytical objectives, multiple cell type configurations were used. For DBP analyses, models were trained using HFFs, HMECs, NPCs, and HepG2, enabling validation with ChIP-seq datasets available for HepG2. For RBP analyses, models were trained using HFFs, HMECs, NPCs, and K562, enabling validation with eCLIP datasets available for K562. For analyses requiring direct comparison of regulatory contribution patterns between HepG2 and K562 for the same NABPs (e.g., **Fig. 3d**), a unified four–cell-type configuration (HFFs, NPCs, HepG2, and K562) was used. This enabled direct comparison of DeepLIFT contribution score distributions across these cell types while maintaining a shared transcriptomic background. In all cases, cell type selection was guided by analytical requirements and the availability of independent validation datasets, and no validation data were used as inputs during model training.

### Gene co-expression–based predictions align with ChIP-seq–defined DNA-binding targets

High-ranking predicted binding regions frequently overlapped with ChIP-seq peaks, particularly within promoter–enhancer intervals. Deep learning–derived DNA-binding site predictions based on gene co-expression data showed substantial concordance with experimentally defined binding sites, especially when broader genomic regions encompassing promoters and putative enhancers are considered. When overlap was assessed within ±100 kb of transcription start sites (TSSs)—a window supported by prior enhancer–gene mapping studies[1,2]—the mean and median overlap ratios reached 0.70 and 0.77, respectively (Ratio A2 / B2; **Supplementary Tables S2** and **S3**). In contrast, overlap was lower when restricted to promoter-proximal regions alone (±5 kb), with corresponding mean and median values of 0.35 and 0.34. Several DNA-binding proteins, including AKAP8, POGK, SPEN, THAP7, ZNF142, and ZNF691, exhibited notably high promoter-region overlap ratios (>0.5), suggesting potential direct transcriptional regulatory roles. For RNA-binding proteins, overlap was evaluated within transcribed regions spanning gene bodies, yielding mean and median overlap ratios of 0.70 and 0.76, respectively (Ratio A2 / B2; **Supplementary Table S4**). Although high-scoring predicted regions frequently coincided with experimentally identified binding sites, not all ChIP-seq or eCLIP peaks correspond to functionally relevant regulatory interactions. Conversely, some model-inferred regulatory targets may reflect indirect or context-dependent regulation not captured by binding assays. Together, these results indicate strong but incomplete concordance between model-inferred regulatory signals and experimental binding data, highlighting the complementary nature of these approaches.

### Functional interpretation of ΔNES-based pathway redistribution across PKM, HNRNPK, and NELFE

To contextualize the ΔNES-based redistribution patterns described in the main text, we examined established biological functions of PKM, HNRNPK, and NELFE in relation to the pathway categories identified by contribution score–based GSEA.

#### PKM

PKM2, the predominant isoform in proliferative and stem-like states, is a central regulator of metabolic reprogramming and exhibits non-canonical nuclear functions that coordinate transcriptional programs during tumorigenesis[3,4]. In addition to its canonical role in glycolysis, PKM2 regulates gene expression through interactions with transcriptional and chromatin-associated factors. Isoform switching between PKM2 and PKM1 accompanies metabolic transitions during differentiation, including neuronal maturation, where PKM1-enriched states are associated with oxidative metabolism and terminal differentiation[5]. PKM has also been identified as a ribosome-associated factor capable of modulating mRNA translation[6], and accumulating evidence suggests that metabolic enzymes, including PKM, can function as non-canonical RNA-binding protein [7].

Beyond its roles in metabolism and transcriptional regulation, PKM2 contributes to immunometabolic processes by modulating cytokine production and immune cell activation[8-10], linking metabolic state to immune signaling outputs. This is consistent with the enrichment of immune-associated pathways observed in ΔNES-based analyses.

PKM2 is also integrated into multiple signaling pathways. For example, receptor tyrosine kinase signaling, including FGFR1, phosphorylates PKM2, and promotes a shift toward a less active dimeric form, contributing to the Warburg effect[11]. PKM2 has also been linked to NOTCH signaling, where coordinated regulation of PKM2 and NOTCH1 influences tumor progression[12], as well as to GPCR-associated pathways such as the GPR3–β-arrestin2–PKM2 axis[13]. In addition, PKM2 contributes to TGF-β signaling by stabilizing receptor complexes and promoting fibrotic responses [14].

PKM2 further participates in diverse context-dependent processes, including lipid metabolism, angiogenesis, and metastatic progression[15-19]. These multifunctional roles position PKM2 at the interface of metabolism, signaling, and gene regulation.

Collectively, these properties are consistent with the ΔNES-inferred redistribution of neuronal, immune, metabolic, and signaling-associated pathways across neural progenitor and leukemia contexts. Rather than driving strictly lineage-specific pathway activation, PKM2 appears to contribute to context-dependent repositioning of shared regulatory networks along the ΔNES axis[20-22].

**HNRNPK**

HNRNPK is a multifunctional RNA-binding protein that integrates transcriptional and post-transcriptional regulatory processes. Haploinsufficiency of HNRNPK predisposes to hematologic malignancies and disrupts proliferation and differentiation programs in myeloid lineages[23]. In stem and progenitor contexts, HNRNPK promotes transcriptional activation of proliferation-associated genes, including MYC and growth factor signaling components, while also regulating RNA stability and processing[24]. These coordinated transcriptional and post-transcriptional functions position HNRNPK as a regulator of the balance between proliferation and differentiation. Accordingly, the ΔNES-based redistribution of immune- and neural-associated pathways across cellular contexts is consistent with a model in which HNRNPK modulates differentiation–proliferation dynamics rather than directing a single lineage-specific regulatory program.

**NELFE**

NELFE is a subunit of the negative elongation factor (NELF) complex that regulates promoter-proximal RNA polymerase II pausing, a mechanism critical for rapid transcriptional responsiveness and developmental gene regulation[25,26]. Oncogenic activation of NELFE enhances MYC-associated transcriptional programs and promote tumor progression[27]. In hematopoietic differentiation, modulation of NELF abundance alters genome-wide transcriptional pausing and influences granulocytic lineage commitment[28]. NELF also regulates progenitor expansion and stem cell maintenance through modulation of p53-dependent transcriptional programs[29]. These findings indicate that NELFE functions as a context-dependent regulator of transcriptional architecture, controlling lineage transitions through regulation of elongation dynamics. The ΔNES redistribution patterns observed for NELFE parallel those of HNRNPK, supporting convergence of distinct RNA-binding proteins on shared regulatory frameworks that coordinate differentiation and proliferative states across cellular contexts.

**Shared signaling architecture**

Across all three proteins, Signal Transduction emerged as the dominant top-level Reactome category. Subpathway analysis revealed enrichment of broadly utilized signaling modules, including FGFR/RTK signaling, IRS-mediated cascades, WNT/BMP pathways, and receptor-associated second-messenger systems. Although neuronal receptor–related pathways were also detected, they represented a minority of enriched subpathways (**Fig. 6c**). This distribution argues against a simple reflection of neuron-specific transcriptional signatures and instead supports the presence of a conserved upstream signaling backbone that is differentially engaged across cellular contexts.

Importantly, the ΔNES-based framework does not imply direct activation or repression of lineage-specific programs by individual proteins. Rather, it captures relative shifts in the positioning of pathway-associated genes along contribution-ranked lists, reflecting redistribution of inferred regulatory influence across cellular states. The convergence of these redistribution patterns across metabolically oriented (PKM), transcriptionally integrative (HNRNPK), and elongation-regulatory (NELFE) proteins supports a model in whith contribution score-based pathway ranking reveals a conserved regulatory scaffold that underlies context-dependent functional organization.

**Supplementary Table S1. Co-expression–based selection enhances DeepLIFT contribution scores of NABP binding sites.**

Binding sites identified through gene co-expression analysis exhibited higher absolute DeepLIFT contribution scores than those derived solely from ChIP-seq data without co-expression-based refinement. This trend was consistently observed for DNA-binding proteins (DBPs) and was more pronounced for RNA-binding proteins (RBPs) across the analyzed cell types. For each protein class, the table reports the mean and median values of positive and negative contribution scores. These results indicate that co-expression–based feature selection preferentially enriches for regulatory interactions with stronger inferred contributions to gene expression prediction. The upper and lower sections of the table correspond to analyses performed using reduced (input × 0.5) and amplified (input × 2) DeepLIFT background references, respectively.

| DBP / RBP | Cell type | Co-expression–based predictions<br>Upper row: mean values<br>Lower row: median values | Non-co-expression–based predictions<br>Upper row: mean values<br>Lower row: median values |
|---|---|---|---|
| DBP | HepG2 | 5.3.E-03 | 3.7.E-03 |
| | | 2.8.E-03 | 1.9.E-03 |
| RBP | K562 | 8.0.E-03 | 3.6.E-03 |
| | | 5.6.E-03 | 1.8.E-03 |
| DBP | NPC | 3.9.E-03 | 3.8.E-03 |
| | | 1.9.E-03 | 1.9.E-03 |
| RBP | NPC | 8.2.E-03 | 3.6.E-03 |
| | | 5.9.E-03 | 1.7.E-03 |
| DBP | HMEC | 4.0.E-03 | 3.8.E-03 |
| | | 1.8.E-03 | 1.8.E-03 |
| RBP | HMEC | 8.9.E-03 | 3.5.E-03 |
| | | 6.4.E-03 | 1.7.E-03 |
| DBP | HFF | 3.9.E-03 | 3.8.E-03 |
| | | 1.7.E-03 | 1.8.E-03 |
| RBP | HFF | 8.8.E-03 | 3.5.E-03 |
| | | 6.5.E-03 | 1.7.E-03 |

| DBP / RBP | Cell type | Co-expression–based:<br>mean (upper)<br>median (lower) | Non-co-expression–based:<br>mean (upper)<br>median (lower) |
|---|---|---|---|
| DBP | HepG2 | 5.28.E-03 | 3.71.E-03 |
| | | 2.78.E-03 | 1.90.E-03 |
| RBP | K562 | 7.95.E-03 | 3.56.E-03 |
| | | 5.60.E-03 | 1.78.E-03 |
| DBP | NPC | 3.88.E-03 | 3.82.E-03 |
| | | 1.87.E-03 | 1.87.E-03 |
| RBP | NPC | 8.20.E-03 | 3.56.E-03 |
| | | 5.90.E-03 | 1.73.E-03 |
| DBP | HMEC | 4.01.E-03 | 3.81.E-03 |
| | | 1.80.E-03 | 1.80.E-03 |
| RBP | HMEC | 8.92.E-03 | 3.55.E-03 |
| | | 6.37.E-03 | 1.68.E-03 |
| DBP | HFF | 3.89.E-03 | 3.82.E-03 |
| | | 1.70.E-03 | 1.79.E-03 |
| RBP | HFF | 8.76.E-03 | 3.55.E-03 |
| | | 6.49.E-03 | 1.71.E-03 |

**Supplementary Table S2. Enrichment of high-contribution promoter regions overlapping experimentally validated DNA-binding protein binding sites.** (See Supplementary Excel file)

This table summarizes the proportion of promoter-proximal regions (±5 kb from transcription start sites [TSSs]) that were identified as high- or low-contribution regions based on DeepLIFT scores and that overlapped with ChIP-seq-validated binding sites for the corresponding DNA-binding proteins (DBPs). Overlap ratios were calculated using multiple threshold definitions to assess the robustness of enrichment results.

High- and low-contribution regions were defined using both rank-based and score-based thresholds applied to the 1,310 nucleic acid–binding proteins (NABPs) analyzed:

**Rank-based thresholds**:

(A1 and B1) top or bottom 20 ranked proteins (rank ≤ 20 or ≥ 1,290)

(A2 and B2) top or bottom 50 ranked proteins (rank ≤ 50 or ≥ 1,260)

**Score-based thresholds**:

(C1 and D1) contribution score ≥ 0.01 or ≤ –0.01

(C2 and D2) contribution score ≥ 0.001 or ≤ –0.001

**Supplementary Table S3. Enrichment of high-contribution promoter–enhancer regions overlapping ChIP-seq–validated DNA-binding protein binding sites.**

This table summarizes the proportion of predicted high- and low-contribution regions—defined within ±100 kb of transcription start sites (TSSs) and encompassing both promoter-proximal and distal enhancer regions—that overlap with ChIP-seq-validated binding sites for the corresponding DNA-binding proteins (DBPs). Overlap ratios were calculated using multiple threshold definitions to assess the robustness of enrichment results.

Across all threshold definitions, substantial concordance was observed between model-derived high-contribution regions and ChIP-seq peaks, supporting the reliability of DeepLIFT contribution scoring for identifying functionally relevant regulatory regions.

High- and low-contribution regions were defined using both rank-based and score-based thresholds applied to the 1,310 nucleic acid–binding proteins (NABPs) analyzed:

**Rank-based thresholds**:

(A1 and B1) top or bottom 20 ranked proteins (i.e., rank ≤ 20 or ≥ 1,290)

(A2 and B2) top or bottom 50 ranked proteins (i.e., rank ≤ 50 or ≥ 1,260)

**Score-based thresholds**:

(C1 and D1) contribution score ≥ 0.01 or ≤ –0.01

(C2 and D2) contribution score ≥ 0.001 or ≤ –0.001

### Supplementary Table S4. Enrichment of high-contribution transcribed regions overlapping eCLIP–validated RNA-binding protein binding sites.

This table summarizes the proportion of predicted high- and low-contribution regions—defined within gene transcribed regions (from transcription start site [TSS] to transcription end site [TES])—that overlap with eCLIP–validated binding sites for the corresponding RNA-binding proteins (RBPs). Overlap ratios were calculated using multiple threshold definitions to assess the robustness of enrichment results. Across all threshold definitions, substantial concordance was observed between model-derived high-contribution regions and eCLIP peaks, supporting the reliability of DeepLIFT contribution scoring for identifying functionally relevant regulatory regions.

High- and low-contribution regions were defined using both rank-based and score-based thresholds applied to the 1,310 nucleic acid–binding proteins (NABPs) analyzed:

**Rank-based thresholds**:

(A1 and B1) top or bottom 20 ranked proteins (i.e., rank ≤ 20 or ≥ 1,290)

(A2 and B2) top or bottom 50 ranked proteins (i.e., rank ≤ 50 or ≥ 1,260)

**Score-based thresholds**:

(C1 and D1) contribution score ≥ 0.01 or ≤ –0.01

(C2 and D2) contribution score ≥ 0.001 or ≤ –0.001

### References


1 Avsec, Z. *et al.* Effective gene expression prediction from sequence by integrating long-range interactions. *Nature methods* **18**, 1196–1203 (2021). https://doi.org/10.1038/s41592-021-01252-x

2 Gasperini, M. *et al.* A Genome-wide Framework for Mapping Gene Regulation via Cellular Genetic Screens. *Cell* **176**, 1516 (2019). https://doi.org/10.1016/j.cell.2019.02.027

3 Yang, W. *et al.* ERK1/2-dependent phosphorylation and nuclear translocation of PKM2 promotes the Warburg effect. *Nat Cell Biol* **14**, 1295–1304 (2012). https://doi.org/10.1038/ncb2629

4 Israelsen, W. J. & Vander Heiden, M. G. Pyruvate kinase: Function, regulation and role in cancer. *Semin Cell Dev Biol* **43**, 43–51 (2015). https://doi.org/10.1016/j.semcdb.2015.08.004

5 He, P. *et al.* PKM2 is a key factor to regulate neurogenesis and cognition by controlling lactate homeostasis. *Stem Cell Reports* **20**, 102381 (2025). https://doi.org/10.1016/j.stemcr.2024.11.011

6 Kejiou, N. S. *et al.* Pyruvate Kinase M (PKM) binds ribosomes in a poly-ADP ribosylation dependent manner to induce translational stalling. *Nucleic acids research* **51**, 6461–6478 (2023). https://doi.org/10.1093/nar/gkad440

7 Castello, A. *et al.* Insights into RNA biology from an atlas of mammalian mRNA-binding proteins. *Cell* **149**, 1393–1406 (2012). https://doi.org/10.1016/j.cell.2012.04.031

8 Toller-Kawahisa, J. E. *et al.* Metabolic reprogramming of macrophages by PKM2 promotes IL-10 production via adenosine. *Cell Rep* **44**, 115172 (2025). https://doi.org/10.1016/j.celrep.2024.115172

9 Yang, P. *et al.* Pyruvate kinase M2 accelerates pro-inflammatory cytokine secretion and cell proliferation induced by lipopolysaccharide in colorectal cancer. *Cell Signal* **27**, 1525–1532 (2015). https://doi.org/10.1016/j.cellsig.2015.02.032

10 Liu, Z., Le, Y., Chen, H., Zhu, J. & Lu, D. Role of PKM2-Mediated Immunometabolic Reprogramming on Development of Cytokine Storm. *Front Immunol* **12**, 748573 (2021). https://doi.org/10.3389/fimmu.2021.748573

11 Kachel, P. *et al.* Phosphorylation of pyruvate kinase M2 and lactate dehydrogenase A by fibroblast growth factor receptor 1 in benign and malignant thyroid tissue. *BMC Cancer* **15**, 140 (2015). https://doi.org/10.1186/s12885-015-1135-y

12 Wang, J. *et al.* Down-regulation of NOTCH1 and PKM2 can inhibit the growth and metastasis of colorectal cancer cells. *Am J Transl Res* **14**, 5455–5465 (2022).

13 Dong, T. *et al.* Activation of GPR3-beta-arrestin2-PKM2 pathway in Kupffer cells stimulates glycolysis and inhibits obesity and liver pathogenesis. *Nat Commun* **15**, 807 (2024). https://doi.org/10.1038/s41467-024-45167-5

14 Gao, J., Zhao, Y., Li, T., Gan, X. & Yu, H. The Role of PKM2 in the Regulation of Mitochondrial Function: Focus on Mitochondrial Metabolism, Oxidative Stress, Dynamic, and Apoptosis. PKM2 in Mitochondrial Function. *Oxid Med Cell Longev* **2022**, 7702681 (2022). https://doi.org/10.1155/2022/7702681

15 Liu, Z., Le, Y. & Lu, D. PKM2: A novel helmsman of lipid metabolism. *Cell Signal* **134**, 111967 (2025). https://doi.org/10.1016/j.cellsig.2025.111967

16 Lee, K. C. Y. *et al.* PKM2 is a key regulator of cardiac lipid metabolism in mice. *Mitochondrion* **85**, 102070 (2025). https://doi.org/10.1016/j.mito.2025.102070

17 Li, L., Zhang, Y., Qiao, J., Yang, J. J. & Liu, Z. R. Pyruvate kinase M2 in blood circulation facilitates tumor growth by promoting angiogenesis. *J Biol Chem* **289**, 25812–25821 (2014). https://doi.org/10.1074/jbc.M114.576934

18 Gomez-Escudero, J. *et al.* PKM2 regulates endothelial cell junction dynamics and angiogenesis via ATP production. *Sci Rep* **9**, 15022 (2019). https://doi.org/10.1038/s41598-019-50866-x

19 He, D. *et al.* Methionine oxidation activates pyruvate kinase M2 to promote pancreatic cancer metastasis. *Molecular cell* **82**, 3045–3060 e3011 (2022). https://doi.org/10.1016/j.molcel.2022.06.005

20 Zhang, X., Lei, Y., Zhou, H., Liu, H. & Xu, P. The Role of PKM2 in Multiple Signaling Pathways Related to Neurological Diseases. *Mol Neurobiol* **61**, 5002–5026 (2024). https://doi.org/10.1007/s12035-023-03901-y

21 Venkatesh, H. S. *et al.* Electrical and synaptic integration of glioma into neural circuits. *Nature* **573**, 539–545 (2019). https://doi.org/10.1038/s41586-019-1563-y

22 Yu, L., Chen, X., Sun, X., Wang, L. & Chen, S. The Glycolytic Switch in Tumors: How Many Players Are Involved? *J Cancer* **8**, 3430–3440 (2017). https://doi.org/10.7150/jca.21125

23 Gallardo, M. *et al.* hnRNP K Is a Haploinsufficient Tumor Suppressor that Regulates Proliferation and Differentiation Programs in Hematologic Malignancies. *Cancer Cell* **28**, 486–499 (2015). https://doi.org/10.1016/j.ccell.2015.09.001

24 Li, J. *et al.* HNRNPK maintains epidermal progenitor function through transcription of proliferation genes and degrading differentiation promoting mRNAs. *Nat Commun* **10**, 4198 (2019). https://doi.org/10.1038/s41467-019-12238-x

25 Adelman, K. & Lis, J. T. Promoter-proximal pausing of RNA polymerase II: emerging roles in metazoans. *Nature reviews. Genetics* **13**, 720–731 (2012). https://doi.org/10.1038/nrg3293

26 Abuhashem, A., Garg, V. & Hadjantonakis, A. K. RNA polymerase II pausing in development: orchestrating transcription. *Open Biol* **12**, 210220 (2022). https://doi.org/10.1098/rsob.210220

27 Dang, H. *et al.* Oncogenic Activation of the RNA Binding Protein NELFE and MYC Signaling in Hepatocellular Carcinoma. *Cancer Cell* **32**, 101–114 e108 (2017). https://doi.org/10.1016/j.ccell.2017.06.002

28 Liu, X. *et al.* Dynamic Change of Transcription Pausing through Modulating NELF Protein Stability Regulates Granulocytic Differentiation. *Blood Adv* **1**, 1358–1367 (2017). https://doi.org/10.1182/bloodadvances.2017008383

29 Robinson, D. C. L. *et al.* Negative elongation factor regulates muscle progenitor expansion for efficient myofiber repair and stem cell pool repopulation. *Dev Cell* **56**, 1014–1029 e1017 (2021). https://doi.org/10.1016/j.devcel.2021.02.025
