## Supplementary Note2 for "Pathway redistribution across cellular states reveals a shared signaling backbone and context-dependent regulatory modules in RNA-binding protein networks"

### Supplementary Note 2

#### Large language model–based inference identifies functional roles of NABP target genes (ChatGPT-based analysis)

This section presents functional interpretations of predicted NABP target genes using a large language model–based inference framework. Detailed prompts and corresponding model outputs are provided to ensure transparency and reproducibility of the analysis.


**Prompts for ChatGPT analysis** Page 2

**DNAJC2** (K562 cells) Page 7

**DNAJC2** (Neural progenitor cells) Page 10

**EIF2S2** (K562 cells) Page 13

**EIF2S2** (Neural progenitor cells) Page 19

**ILF2** (K562 cells) Page 21

**ILF2** (Neural progenitor cells) Page 25

**PKM** (K562 cells) Page 28

**PKM** (Neural progenitor cells) Page 34

**POLR2G** (K562 cells) Page 39

**POLR2G** (Neural progenitor cells) Page 45

**RBM27** (K562 cells) Page 48

**SPEN** (HepG2 cells) Page 52

**ZBTB34** (HepG2 cells) Page 56

**ZC3H4** (HepG2 cells) Page 60

**ZKSCAN5** (HepG2 cells) Page 67

**ZNF446** (HepG2 cells) Page 74

**ZNF557** (HepG2 cells) Page 79

#### Prompt 1: Functional interpretation of NABP target genes

*Attached files: functional_annotation.txt and genelist.txt*

A large language model was asked to generate a concise biological interpretation of each predicted NABP target gene set using two inputs: enriched functional annotation terms from PANTHER analysis (functional_annotation.txt) and the corresponding target gene list (genelist.txt). The prompt instructed the model to identify the most prominent biological process represented by the gene set, integrate related functional categories into a coherent biological interpretation, avoid generic process names, and assign a confidence score from 0.00 to 1.00. The confidence score was defined as the estimated proportion of genes supporting the selected process name. If no coherent biological process could be identified, the model was instructed to name the set "System of unrelated proteins and DNAs/RNAs" and assign a score of 0.00.

**Prompt for ChatGPT analysis:**

You are an efficient and insightful assistant to a molecular biologist. Write a critical analysis of the biological processes performed by this system of interacting proteins and DNAs/RNAs.

In your analysis, focus particularly on the functional categories listed in the attached document titled *functional_annotation.txt.* which summarizes the enriched annotations identified by PANTHER functional enrichment analysis. However, do not restrict your analysis to these categories—also incorporate related or functionally adjacent processes that connect these categories into a coherent biological network.

Base your analysis on prior knowledge. After completing your analysis, propose a short descriptive name for the most prominent biological process performed by the system. Then assign a confidence score to the process name you selected. This score should follow the name in parentheses and range from 0.00 to 1.00. A score of 0.00 indicates the lowest confidence, while 1.00 reflects the highest confidence. This score helps gauge how accurately the chosen name represents the functions and activities within the system of interacting proteins and DNAs/RNAs. When determining your score, consider the proportion of genes in the protein and DNA/RNA system that participate in the identified biological process. For instance, if you select "Ribosome biogenesis" as the process name but only a few genes in the system contribute to this process, the score should be lower compared to a scenario where a majority of the genes are involved in "Ribosome biogenesis". Put your chosen name at the top of the analysis as 'Process: <name>". Be concise, do not use unnecessary words. Be factual, do not editorialize. Be specific, avoid overly general statements such as 'the proteins, DNAs and RNAs are involved in various cellular processes' Avoid listing facts about individual proteins, DNAs and RNAs. Instead, try to group proteins and DNAs/RNAs with similar functions and discuss their interplay, synergistic or antagonistic effects, and functional integration within the system. Also, avoid choosing generic process names such as 'Cellular Signaling and Regulation'. If you cannot identify a prominent biological process for the proteins and DNAs/RNAs in the system, I want you to communicate this in your analysis and name the process: "System of unrelated proteins and DNAs/RNAs". Provide a score of 0.00 for a "System of unrelated proteins and DNAs/RNAs". To help you in your work, I am providing an example system of interacting proteins and DNAs/RNAs and the corresponding example analysis output. The example system of interacting proteins and DNAs/RNAs is: PDX1, SLC2A2, NKX6-1, GLP1, GCG.

The example analysis output is: Process: Pancreatic development and glucose homeostasis (0.96)

1. PDX1 is a homeodomain transcription factor involved in the specification of the early pancreatic epithelium and its subsequent differentiation. It activates the transcription of several genes including insulin, somatostatin, glucokinase and glucose transporter type.

2. It is essential for maintenance of the normal hormone-producing phenotype in the pancreatic beta-cell. In pancreatic acinar cells, it forms a complex with PBX1b and MEIS2b and mediates the activation of the ELA1 enhancer.

3. NKX6-1 is also a transcription factor involved in the development of pancreatic beta-cells during the secondary transition. Together with NKX2-2 and IRX3, controls the generation of motor neurons in the neural tube and belongs to the neural progenitor factors induced by Sonic Hedgehog (SHH) signals.

4.GCG and GLP1, respectively glucagon and glucagon-like peptide 1, are involved in glucose metabolism and homeostasis. GCG raises blood glucose levels by promoting gluconeogenesis and is the counter regulatory hormone of

Insulin. GLP1 is a potent stimulator of Glucose-Induced Insulin Secretion (GSIS). Plays roles in gastric motility and suppresses blood glucagon levels. Promotes growth of the intestinal epithelium and pancreatic islet mass both by islet neogenesis and islet cell proliferation.

5. SLC2A2, also known as GLUTZ, is a facilitative hexose transporter. In hepatocytes, it mediates bi-directional transport of glucose across the plasma membranes, while in the pancreatic beta-cell, it is the main transporter responsible for glucose uptake and part of the cell's glucose-sensing mechanism. It is involved in glucose transport in the small intestine and kidney too.

To summarize, the genes in this set are involved in the specification, differentiation, growth and functionality of the pancreas, with a particular emphasis on the pancreatic beta-cell. Particularly, the architecture of the pancreatic islet ensures proper glucose sensing and homeostasis via a number of different hormones and receptors that can elicit both synergistic and antagonistic effects in the pancreas itself and other peripheral tissues.

The list of interacting target genes, as defined by the interaction between a protein and DNA/RNA, is provided in the attached document titled 'genelist.txt '.

#### Prompt 2: Identification of genes supporting the inferred process name

*Attached files: genelist.txt and functional_analysis.txt*

A second prompt was used to identify genes that support the process name generated in Prompt 1. The model was provided with the gene list (*genelist.txt*) and the corresponding functional interpretation (*functional_analysis.txt*). The model was instructed to identify genes for which the text makes clear, non-speculative assertions supporting the inferred process name, assign confidence levels (High, Medium, Low), and provide structured output including a summary, gene-level explanations, and a list of supporting genes.

**Prompt for ChatGPT analysis:**

Analyze the provided text describing a gene set and its inferred biological function. Your task is to identify genes that support the proposed process name based solely on the provided text. Context: Gene sets are groups of genes that share common biological functions, pathways, or other characteristics. Naming these sets based on their predominant functions helps researchers quickly understand their significance.

Input:
1. A list of gene symbols in the gene set, provided in comma-separated format, is listed in the attached document titled 'genelist.txt '.
2. The analysis text is provided in the attached document titled 'functional_analysis.txt'.
3. The suggested name for the gene set is indicated in the first sentence beginning 'Process:' in the attached document titled 'functional_analysis.txt'.

Instructions:
1. Evaluate each gene from the provided list that is mentioned in the text.
2. Determine if the text makes a definite assertion about the gene that supports the given name. - A definite assertion clearly states a gene's function or role without using speculative language. - Example of a definite assertion: "XRCC1 is involved in the DNA damage response" - Example of a non-definite assertion: "E2F1 may be involved in homologous recombination".
3. If a gene is mentioned multiple times, consider the strongest assertion made about it.
4. In case of contradictory statements about a gene, favor the most recent or specific assertion.
5. For each gene you determine supports the name: - Briefly explain your reasoning (max 50 words per gene) - Assign a confidence level (High, Medium, Low) based on the strength of the assertion.
6. Handle acronyms or alternative gene names as equivalent to official gene symbols.
7. If no genes seem to support the name or if all genes support it, state this observation.

Output your analysis in the following format:
-- Summary -- [Provide a brief summary (max 100 words) of why the selected genes support the given name]
-- Explanation -- [Gene Symbol}: [Confidence Level] [Explanation of reasoning (max 50 words)] [Repeat for each supporting gene]

-- genes supporting the name: [List of gene symbols of genes supporting the name]
Do not critique the analysis or the name. Base your evaluation solely on the information provided in the text.

#### AI-assisted analysis statement

Large language model–based gene set interpretation was performed using ChatGPT to assist functional annotation of predicted NABP target genes. The model was provided only with gene lists and PANTHER-derived functional annotation summaries generated in this study.

ChatGPT outputs were used to generate structured functional summaries and to identify genes supporting each inferred functional label. All outputs were reviewed and curated by the authors. Statistical enrichment analyses using PANTHER and GSEA served as the primary basis for biological interpretation.

### DNAJC2 K562

#### Process: RNA processing–coupled ribosome biogenesis and translation with proteostasis surveillance (0.88)

The enrichment profile is dominated by RNA metabolism: major/minor spliceosomal splicing, mRNA processing/transport, and extensive rRNA processing/ribosome biogenesis that culminate in translational initiation/elongation. These biosynthetic flows are wired to quality-control circuits—proteasome-mediated protein catabolism and the unfolded-protein response—plus DNA repair and mitochondrial gene expression, forming a coordinated "make-and-proofread" system for gene expression and proteome maintenance. The gene list is heavily populated by spliceosomal factors, rRNA/ribosome assembly proteins, translation factors and chaperones that operationalize these modules, explaining the high confidence in this integrated process.

##### 1. RNA processing and export

Strong terms for "RNA splicing (major/minor), mRNA processing, RNA modification, ncRNA processing, RNA export from nucleus, RNA transport, RNA localization" indicate a core spliceosome/RNP assembly engine that prepares coding and noncoding RNAs for translation and ribosome assembly. These steps align upstream of the ribosome pathway.

##### 2. Ribosome biogenesis and rRNA metabolism

"rRNA processing, rRNA metabolic process, ribosome biogenesis, ribonucleoprotein complex biogenesis" connect nucleolar maturation with cytoplasmic ribosome completion, ensuring abundant translational capacity. This is a major signal in the annotation set and central to the network's output.

##### 3. Translation and initiation control

"Translational initiation, translation, peptide biosynthetic process, translation initiation complex formation, ribosomal scanning and start codon recognition" show that the system pushes prepared transcripts into protein synthesis, linking initiation factors and ribosomal subunits to productive elongation.

##### 4. Proteostasis and stress-response coupling

"Proteasome-mediated ubiquitin-dependent protein catabolic process, ubiquitin–proteasome pathway, protein folding, cellular/ER unfolded protein response, cellular response to stress/heat/chemical stress" indicate parallel surveillance that degrades misfolded or surplus proteins and adjusts folding capacity, buffering the high biosynthetic flux from RNA→protein.

##### 5. Genome maintenance and cell-cycle gating (supporting)

“DNA repair, G1/S DNA damage checkpoints, cell cycle checkpoints” provide upstream safeguards that stabilize transcription/replication interfaces during intense ribosome/translation programs. These terms are present but less dominant than RNA/translation categories.

##### 6. Organelle and mitochondrial support (supporting)

“Mitochondrial gene expression/translation, mitochondrial protein import/degradation” supply ATP and co-factors for chaperones, export motors, and ribosome assembly; smaller signals for “regulation of cilium assembly” reflect adjacent structural organization rather than the central theme.

Why this name and score

Most enriched categories cluster along a single axis—RNA processing → ribosome biogenesis → translation—tightly integrated with proteostasis safeguards. Given the density of these terms and the composition of the provided gene list, this represents the predominant function of the system (high, but not maximal, confidence).

-- Summary --

The text identifies a network centered on RNA processing, ribosome biogenesis, translation, and proteostasis. Genes supporting the name “RNA processing–coupled ribosome biogenesis and translation with proteostasis surveillance” are those directly implicated in splicing, rRNA processing, ribosomal structure or translation initiation, protein folding/degradation, and mitochondrial translational support. Their roles establish an integrated biosynthetic and quality-control axis.

-- Explanation --

**HNRNPM:** [High] Described as a spliceosomal RNA-binding protein central to “RNA splicing (major/minor)” and mRNA maturation, supporting the RNA processing component.

**SF3B2:** [High] A core spliceosome subunit explicitly included in “RNA splicing” enrichment, directly anchoring the RNA processing module.

**SNRPB2:** [High] Part of the small nuclear RNP complex; supports spliceosomal function in pre-mRNA processing within the RNA-processing axis.

**PRPF31:** [High] Known spliceosomal assembly factor; directly contributes to RNA splicing and mRNA processing identified in the text.

**EIF3H / EIF3G / EIF5 / EIF4E / EIF3E:** [High] Translation initiation factors linking RNA output to “translation initiation” and “ribosomal scanning,” supporting the translation node.

**FBL / NOP14 / NIP7 / UTP6 / UTP11:** [High] rRNA-processing/ribosome assembly components underpinning “rRNA metabolic process” and “ribosome biogenesis.”

**MRPL3 / MRPL32 / MRPS17 / MRPS22 / MRPL47 / MRPS5:** [Medium] Mitochondrial ribosomal proteins linking organelle gene expression to translational output, supporting “mitochondrial gene expression/translation.”

**PSMC1 / PSMD6 / PSMD7 / PSMD11 / PSMD14 / PSMA2 / PSMA3:** [High] Proteasome regulatory and catalytic subunits validating “proteasome-mediated ubiquitin-dependent protein catabolism.”

**DNAJA1 / DNAJB6 / HSPD1 / DNAJB11:** [High] Molecular chaperones supporting “protein folding” and the “unfolded protein response.”

**NAMPT / TXNL1 / GLRX3 / PRDX3:** [Medium] Participate in redox maintenance and stress response coupled to proteostasis modules.

**RAD50 / NBN / E2F4:** [Medium] DNA-repair and checkpoint control factors supporting “DNA repair” and “cell-cycle gating” submodules.

**ISCA1 / YAE1 / NMD3:** [Medium] Iron–sulfur cluster assembly and ribosome export components, reinforcing “Fe–S cluster assembly” and “rRNA processing → ribosome maturation” connections.

**OPA1 / TIMM17A / PMPCB:** [Medium] Mitochondrial import and processing factors linking translation with “mitochondrial protein import/degradation.”

**CWC15 / DDX5 / DDX56 / DDX39A:** [High] RNA helicases involved in spliceosome dynamics and RNA export, directly supporting the “RNA processing and export” component.

**GTF3A / POLR3D / POLR2J / TAF2:** [Medium] General transcription machinery feeding the RNA biogenesis stream, supporting the RNAPII-based coupling of transcription to RNA processing.

**HAT1 / HDAC2:** [Medium] Chromatin modifiers influencing transcriptional access for rRNA/ribosome biogenesis genes, functionally adjacent to core processes.

-- genes supporting the name:

HNRNPM, SF3B2, SNRPB2, PRPF31, EIF3H, EIF3G, EIF5, EIF4E, EIF3E, FBL, NOP14, NIP7, UTP6, UTP11, MRPL3, MRPL32, MRPS17, MRPS22, MRPL47, MRPS5, PSMC1, PSMD6, PSMD7, PSMD11, PSMD14, PSMA2, PSMA3, DNAJA1, DNAJB6, HSPD1, DNAJB11, NAMPT, TXNL1, GLRX3, PRDX3, RAD50, NBN, E2F4, ISCA1, YAE1, NMD3,

OPA1, TIMM17A, PMPCB, CWC15, DDX5, DDX56, DDX39A, GTF3A, POLR3D, POLR2J, TAF2, HAT1, HDAC2

### DNAJC2 NPC

#### Process: Co-transcriptional mRNA processing-to-translation with ER targeting and ubiquitin–proteasome quality control (0.86)

The enriched categories map a tightly coupled gene-expression pipeline: RNA Pol II initiation/elongation and mRNA capping flow into spliceosome-driven processing and nuclear export; exported transcripts engage cap-dependent translation and ribosome initiation/scanning; nascent polypeptides are targeted to/inserted into ER, where folding is monitored and misfolded species are cleared by ERAD and the ubiquitin–proteasome system. Stress and checkpoint modules (UPR, DNA damage response) gate throughput, while mitochondrial import/respiratory chain supply energy for proteostasis. This network explains the prevalence of RNA processing factors, initiation factors, ER/secretory components, and proteasome/ubiquitin enzymes in the gene list.

##### 1. RNAPII transcription → co-transcriptional RNA processing

Strong enrichment for "RNA Polymerase II Transcription (initiation, elongation, termination), mRNA capping, 3′-end processing, and (major/minor) spliceosome" indicates that transcript maturation is a core function coordinated with *CTD* phosphorylation states and early elongation control.

##### 2. Nuclear export and mRNP remodeling

"mRNA export from nucleus/nuclear transport/RNA localization" connect processed transcripts to cytoplasmic fate, positioning *TREX*/export factors alongside hnRNPs in the set.

##### 3. Cap-dependent translation initiation

Enrichment for "Eukaryotic/Cap-dependent translation initiation, ribosomal scanning and start codon recognition, formation of ternary/*43S* complexes" supports a dominant role in initiating cytoplasmic protein synthesis.

##### 4. Targeting to and translocation across the ER

"Protein targeting to ER/establishment of protein localization to ER" links translation to co-translational insertion and secretory pathway entry.

##### 5. Folding, ER quality control, and degradation

"Response to ER stress/*UPR*, *ERAD* pathway, ubiquitin-dependent/proteasome-mediated protein catabolic process" indicate surveillance and clearance of misfolded ER clients and regulatory proteins; many proteasome and ubiquitin pathway terms/pathways are enriched.

##### 6. Checkpoint and repair gating

"DNA repair (*GG-NER*/*TC-NER*), *TP53*-regulated DNA damage responses, G1/S & G2/M checkpoints" couple gene-expression throughput to genome integrity.

##### 7. Energetic support

"Mitochondrial protein import/respiratory electron transport chain" provide ATP and redox capacity required for translation and proteostasis.

Why this is the predominant process

Across the provided gene list, many entries are annotated as RNA splicing/processing factors, translation initiation/elongation components, ER/secretory pathway proteins, chaperones, ubiquitin–proteasome elements, and nuclear transport factors—matching the above categories and indicating that the majority of genes contribute directly to this coupled expression-to-proteostasis axis.

**Confidence rationale (0.86):** A large proportion of enriched terms and listed genes fall within RNA processing, export, translation initiation, ER targeting, and proteasome/ERAD, with consistent stress/checkpoint overlays; fewer genes map to peripheral signaling modules, supporting a high but not maximal score.

-- Summary --

The text supports the name **"Co-transcriptional mRNA processing-to-translation with ER targeting and ubiquitin–proteasome quality control"** through explicit mentions of genes contributing to RNA polymerase II transcription, RNA splicing and export, translation initiation, ER targeting, protein folding, and proteasomal degradation. These genes collectively establish a tightly integrated network that links transcription to protein synthesis and quality control.

-- Explanation --

**POLR2J:** [High] Identified as a component of RNA Polymerase II, directly involved in mRNA synthesis during co-transcriptional processing, supporting the transcriptional initiation portion of the process.

**SRSF2:** [High] A splicing factor mediating co-transcriptional pre-mRNA splicing, linking transcription to mature mRNA formation.

**PRPF18:** [High] A spliceosome component participating in splicing catalysis; strongly supports the mRNA processing segment.

**HNRNPH3:** [High] hnRNP involved in RNA processing and export, ensuring mature mRNA availability for translation.

**MAGOH / MAGOHB:** [High] Components of the exon junction complex, essential for mRNA splicing and subsequent export to the cytoplasm.

**WTAP:** [Medium] Engaged in mRNA methylation and splicing regulation, bridging transcriptional and post-transcriptional RNA modification.

**EIF3M / EIF3H / EIF4E:** [High] Translation initiation factors central to cap-dependent translation initiation, connecting mRNA export to ribosome engagement.

**KARS / GARS:** [Medium] Aminoacyl-tRNA synthetases ensuring charged tRNAs for translation; directly related to protein synthesis.

**SEC62 / SEC63:** [High] ER translocon components guiding nascent peptide insertion into the endoplasmic reticulum, integral to ER targeting.

**HSP90B1 / DNAJB11 / SELENOS:** [High] Chaperones ensuring protein folding and quality control within the ER lumen.

**PSMC1 / PSMD11 / PSMA2 / ADRM1 / PSMC4:** [High] Core and regulatory proteasome subunits mediating ubiquitin–proteasome degradation of misfolded proteins; essential to the quality-control axis.

**NFKBIB / HSF1:** [Medium] Regulators of transcriptional stress and protein homeostasis; modulate chaperone expression during proteotoxic stress.

**NDUFA5 / MRPL32:** [Medium] Mitochondrial translation and respiratory components providing energy for proteostasis and ER function.

**UFM1:** [High] Ubiquitin-like modifier linking translation to ER quality control and proteasomal degradation, reinforcing the system's integrated nature.

**RNF6 / RNF7 / RNF216:** [Medium] E3 ubiquitin ligases involved in targeted degradation, supporting the ubiquitin–proteasome linkage.

**AXIN1:** [Medium] Scaffold protein coordinating signaling-to-transcriptional feedback; integrates stress response within the broader network.

-- genes supporting the name:

POLR2J, SRSF2, PRPF18, HNRNPH3, MAGOH, MAGOHB, WTAP, EIF3M, EIF3H, EIF4E, KARS, GARS, SEC62, SEC63, HSP90B1, DNAJB11, SELENOS, PSMC1, PSMD11, PSMA2, ADRM1, PSMC4, NFKBIB, HSF1, NDUFA5, MRPL32, UFM1, RNF6, RNF7, RNF216, AXIN1

### EIF2S2 K562

#### Process: Integrated stress response–driven amino acid homeostasis and ER proteostasis remodeling (0.83)

The enriched categories center on amino acid transport/biosynthesis, the Unfolded Protein Response (UPR), and EIF2-kinase–mediated integrated stress response (ISR), with broad RNA processing/translation-control terms providing throughput. Specifically, pathways such as "Amino acid transport across the plasma membrane," "Response of EIF2AK4 (GCN2) to amino acid deficiency," "PERK regulates gene expression," "ATF4 activates genes in response to ER stress," and "Unfolded Protein Response (UPR)" anchor a nutrient/ER-stress program. Parallel modules for KEAP1–NFE2L2 antioxidant defense, NMD/NMD(EJC)-mediated transcript remodeling, and extensive mRNA/rRNA processing couple stress signaling to proteostasis and translational capacity. Vesicle/ER–Golgi trafficking and membrane/vesicle transport support secretory load adaptation; selective autophagy/mitophagy pathways provide organelle quality control during stress.

Functionally, the genelist contains hallmark ISR/UPR effectors (ATF4, DDIT3/CHOP, HSPA5/BiP), amino-acid transporters and biosynthetic enzymes (SLC7A5/SLC1A5/SLC38A1–2, ASNS, PSAT1), antioxidant/KEAP1–NFE2L2 nodes

(KEAP1, NFE2L2, TXNRD1, GCLM), and transcriptome-remodeling machinery (UPF2/UPF3B, multiple spliceosome/rRNA-processing factors). Autophagy/ERAD and trafficking factors (ATG5/ATG12, CHMP4A/CHMP4B, SEC63/LMAN1, COPB2, RAB/RABEP1) integrate with stress pathways to restore proteostasis. Collectively, the network depicts an ISR-anchored response that boosts amino acid import/synthesis, reprograms translation/RNA metabolism, and expands ER folding/clearance capacity to maintain cell viability under nutrient/ER stress.

#### 1. Amino acid supply and sensing

Enrichment for “amino acid transmembrane transport/biosynthetic process” and “amino acid import across plasma membrane” matches transporters (*SLC7A5*, *SLC7A1*, *SLC38A1/2*, *SLC1A5*) and biosynthetic enzymes (*ASNS*, *PSAT1*, *ARG2*), ensuring substrate influx during stress. [1][2][3]

---

[1] Nicklin, Paul, Bergman, Philip, Zhang, Bailin, Triantafellow, Ellen, Wang, Henry, Nyfeler, Beat, Yang, Haidi, Hild, Marc, Kung, Charles, Wilson, Christopher, Myer, Vic E, MacKeigan, Jeffrey P, Porter, Jeffrey A, Wang, Y Karen, Cantley, Lewis C, Finan, Peter M, Murphy, Leon O. ""Bidirectional transport of amino acids regulates mTOR and autophagy."" Cell, 2009, pp. 521-34, doi: https://doi.org/10.1016/j.cell.2008.11.044

[2] Kandasamy, Palanivel, Gyimesi, Gergely, Kanai, Yoshikatsu, Hediger, Matthias A. ""Amino acid transporters revisited: New views in health and disease."" Trends in biochemical sciences, 2018, pp. 752-789, doi: https://doi.org/10.1016/j.tibs.2018.05.003

[3] Nachef, Marianna, Ali, Alaa Kassim, Almutairi, Saeedah Musaed, Lee, Seung-Hwan. ""Targeting SLC1A5 and SLC3A2/SLC7A5 as a Potential Strategy to Strengthen Anti-Tumor Immunity in the Tumor Microenvironment."" Frontiers in immunology, 2021, pp. 624324, doi: https://doi.org/10.3389/fimmu.2021.624324

#### 2. ISR/UPR signaling axis

“Response of *EIF2AK4* (*GCN2*) to amino acid deficiency,” “PERK regulates gene expression,” “*ATF4* activates genes,” and “*UPR*” align with *ATF4*, *DDIT3*, *HSPA5*, *HERPUD1*, *EIF4EBP1* and cofactors (*CEBPB*/*CEBPZ*), implementing translational attenuation and transcriptional induction of stress-mitigation genes. [1][2][3]

---

[1] Pakos-Zebrucka, Karolina, Koryga, Izabela, Mnich, Katarzyna, Ljujic, Mila, Samali, Afshin, Gorman, Adrienne M. ""The integrated stress response."" EMBO reports, 2016, pp. 1374-1395, doi: https://doi.org/10.15252/embr.201642195

[2] Donnelly, Neysan, Gorman, Adrienne M, Gupta, Sanjeev, Samali, Afshin. ""The eIF2α kinases: their structures and functions."" Cellular and molecular life sciences : CMLS, 2013, pp. 3493-511, doi: https://doi.org/10.1007/s00018-012-1252-6

[3] Quirós, Pedro M, Prado, Miguel A, Zamboni, Nicola, D'Amico, Davide, Williams, Robert W, Finley, Daniel, Gygi, Steven P, Auwerx, Johan. ""Multi-omics analysis identifies ATF4 as a key regulator of the mitochondrial stress response in mammals."" The Journal of cell biology, 2017, pp. 2027-2045, doi: https://doi.org/10.1083/jcb.201702058

#### 3. Antioxidant and detoxification buffering

"*KEAP1–NFE2L2* pathway" and "*NFE2L2* regulating anti-oxidant/detoxification enzymes" map to *KEAP1*, *NFE2L2*, *TXNRD1*, *GCLM*, coordinating redox homeostasis as ER/translation stress elevates ROS. [1][2][3]

---

[1] Li, Dan, Shao, Rong, Wang, Na, Zhou, Nan, Du, Kaili, Shi, Jiahui, Wang, Yihan, Zhao, Zhuangzhuang, Ye, Xin, Zhang, Xiaoli, Xu, Haoxing. ""Sulforaphane Activates a lysosome-dependent transcriptional program to mitigate oxidative stress."" Autophagy, 2021, pp. 872-887, doi: https://doi.org/10.1080/15548627.2020.1739442

[2] Mukherjee, Rukmini, Bhattacharya, Anshu, Mello-Vieira, João, Kuncha, Santosh Kumar, Hoffmann, Marina, Gonzalez, Alexis, Rathore, Rajeshwari, Chadha, Attinder, Shin, Donghyuk, Colby, Thomas, Matic, Ivan, Mukherjee, Shaeri, Misra, Mohit, Dikic, Ivan. ""Serine ubiquitination of SQSTM1 regulates NFE2L2-dependent redox homeostasis."" Autophagy, 2025, pp. 407-423, doi: https://doi.org/10.1080/15548627.2024.2404375

[3] Komatsu, Masaaki, Kurokawa, Hirofumi, Waguri, Satoshi, Taguchi, Keiko, Kobayashi, Akira, Ichimura, Yoshinobu, Sou, Yu-Shin, Ueno, Izumi, Sakamoto, Ayako, Tong, Kit I, Kim, Mihee, Nishito, Yasumasa, Iemura, Shun-ichiro, Natsume, Tohru, Ueno, Takashi, Kominami, Eiki, Motohashi, Hozumi, Tanaka, Keiji, Yamamoto, Masayuki. ""The selective autophagy substrate p62 activates the stress responsive transcription factor Nrf2 through inactivation of Keap1."" Nature cell biology, 2010, pp. 213-23, doi: https://doi.org/10.1038/ncb2021

#### 4. RNA processing and translation control

Strong enrichment for “mRNA splicing,” “rRNA processing,” “ribosome biogenesis,” “NMD,” and “RNAPII termination/3′-end processing” corresponds to extensive spliceosome/exosome and pre-rRNA factors (e.g., *UPF2*/*UPF3B*, *ZCCHC8*, *CPSF2/7*, *BYSL*, *UTP18*/*WDR43*, *RRP15*), enabling rapid remodeling of the transcriptome/proteome under *ISR*. [1][2][3]

---

[1] Serin, G, Gersappe, A, Black, J D, Aronoff, R, Maquat, L E. ""Identification and characterization of human orthologues to Saccharomyces cerevisiae Upf2 protein and Upf3 protein (Caenorhabditis elegans SMG-4)."" Molecular and cellular biology, 2001, pp. 209-23, doi: https://doi.org/10.1128/MCB.21.1.209-223.2001

[2] Adachi, Kenjiro, Soeta-Saneyoshi, Chie, Sagara, Hiroshi, Iwakura, Yoichiro. ""Crucial role of Bysl in mammalian preimplantation development as an integral factor for 40S ribosome biogenesis."" Molecular and cellular biology, 2007, pp. 2202-14, doi: https://doi.org/10.1128/MCB.01908-06

[3] Dong, Zhixiong, Zhu, Changjun, Zhan, Qimin, Jiang, Wei. ""The roles of RRP15 in nucleolar formation, ribosome biogenesis and checkpoint control in human cells."" Oncotarget, 2017, pp. 13240-13252, doi: https://doi.org/10.18632/oncotarget.14658

#### 5. Proteostasis, ER/secretory pathway, and autophagy

“*ATF6* activates chaperone genes,” “membrane trafficking,” and *ERAD*/*ESCRT* terms align with *HSPA5*/*SEC63*/*LMAN1*, *COPB2*/*SEC23B*, *CHMP4A/B*, and *ATG5*/*ATG12*; these coordinate folding, export, and clearance of misfolded proteins. [1][2][3]

---

[1] Hetz, Claudio. ""The unfolded protein response: controlling cell fate decisions under ER stress and beyond."" Nature reviews. Molecular cell biology, 2012, pp. 89-102, doi: https://doi.org/10.1038/nrm3270

[2] Wang, Jie, Lee, Jessica, Liem, David, Ping, Peipei. ""HSPA5 Gene encoding Hsp70 chaperone BiP in the endoplasmic reticulum."" Gene, 2017, pp. 14-23, doi: https://doi.org/10.1016/j.gene.2017.03.005

[3] Li, Baozong, Gao, Bo, Ye, Linbai, Han, Xue, Wang, Wei, Kong, Lingbao, Fang, Xiaonan, Zeng, Yingchun, Zheng, Hong, Li, Shanshan, Wu, Zhenghui, Ye, Li. ""Hepatitis B virus X protein (HBx) activates ATF6 and IRE1-XBP1 pathways of unfolded protein response."" Virus research, 2007, pp. 44-9, doi: https://doi.org/10.1016/j.virusres.2006.09.011

#### 6. Organelle quality control

“Receptor-mediated mitophagy” and vesicle-trafficking Rab/GEF annotations (*RAB5A*/*RABEP1*, *RAB6A*, *RABEPK*, *ARFRP1*) reinforce stress-coupled organelle turnover and cargo routing.

Why this name and score

Most top categories and many genes converge on amino acid scarcity/ER stress signaling (ISR/UPR), with coordinated antioxidant defense and deep RNA-processing/translation reprogramming. Given the breadth and depth of these modules in the genelist, this represents the dominant, integrated process (estimated majority coverage), supporting a high but not maximal confidence.

-- Summary --

The suggested name **“Integrated stress response–driven amino acid homeostasis and ER proteostasis remodeling”** is strongly supported by genes encoding central ISR and UPR regulators (ATF4, DDIT3, HSPA5, HERPUD1), amino acid transporters and biosynthetic enzymes (SLC7A5, SLC1A5, SLC38A1, SLC38A2, ASNS, PSAT1, ARG2), and redox regulators (KEAP1, NFE2L2, TXNRD1, GCLM). Supporting modules include mRNA processing (UPF2, UPF3B, CPSF2, CSTF1), vesicle trafficking and ERAD (SEC63, COPB2, CHMP4A, CHMP4B), and autophagy/mitophagy (ATG5, ATG12, OPTN). Together, these genes create a coordinated network responding to nutrient and ER stress to restore cellular proteostasis and metabolic equilibrium.

-- Explanation --

**ATF4:** [High] Defined as a transcriptional effector of the integrated stress response, activating amino acid biosynthetic and stress-responsive genes under nutrient deprivation.

**DDIT3 (CHOP):** [High] Described as a UPR effector mediating transcriptional reprogramming during ER stress, downstream of ATF4 and PERK.

**HSPA5 (BiP):** [High] Identified as a central ER chaperone induced by UPR activation to enhance folding capacity and proteostasis.

**HERPUD1:** [High] Cited as an ER-stress-inducible component of the ATF6 branch of the UPR, contributing to protein quality control.

**SLC7A5:** [High] Explicitly linked to amino acid transport and import, sustaining amino acid supply during stress.

**SLC1A5 / SLC38A1 / SLC38A2:** [High] Noted as key plasma-membrane amino acid transporters supporting nutrient repletion and ISR activation.

**ASNS / PSAT1 / ARG2:** [High] Annotated under “amino acid biosynthetic process,” producing asparagine and serine to restore nitrogen balance during stress.

**KEAP1 / NFE2L2 (NRF2):** [High] Identified within the antioxidant defense module mitigating oxidative stress downstream of UPR and ISR activation.

**TXNRD1 / GCLM:** [Medium] Downstream effectors of NFE2L2 maintaining redox balance under ER-stress conditions.

**UPF2 / UPF3B:** [High] Represent NMD core components ensuring mRNA surveillance and turnover of defective transcripts during ISR-mediated transcriptional remodeling.

**CPSF2 / CSTF1:** [Medium] Involved in mRNA 3′-end processing and RNA Pol II termination, linking RNA maturation to translational reprogramming.

**SEC63 / COPB2:** [High] ER translocon and vesicle-coat proteins involved in trafficking and co-translational translocation, supporting proteostasis remodeling.

**CHMP4A / CHMP4B:** [Medium] ESCRT components facilitating ERAD and vesicular clearance of misfolded proteins.

**ATG5 / ATG12 / OPTN:** [High] Core autophagy and mitophagy mediators coupling proteostasis to organelle turnover under nutrient stress.

**EIF4EBP1:** [Medium] Translational repressor activated by ISR signaling, reducing global translation while preserving stress-adaptive mRNA translation.

**CEBPB / CEBPZ:** [Medium] ISR-linked transcriptional cofactors enhancing ATF4 target gene expression.

**UPR-associated chaperones (LMAN1, CALU):** [Medium] Contribute to ER folding quality control, aligning with the proteostasis aspect of the name.

-- genes supporting the name:
ATF4, DDIT3, HSPA5, HERPUD1, SLC7A5, SLC1A5, SLC38A1, SLC38A2, ASNS, PSAT1, ARG2, KEAP1, NFE2L2, TXNRD1, GCLM, UPF2, UPF3B, CPSF2, CSTF1, SEC63, COPB2, CHMP4A, CHMP4B, ATG5, ATG12, OPTN, EIF4EBP1, CEBPB, CEBPZ, LMAN1, CALU

### EIF2S2 NPC

#### Process: Integrated stress response coordinating UPR, antioxidant defense, and amino-acid/secretory metabolism (0.86)

The enrichment profile centers on RNA polymerase II transcriptional control and co-transcriptional RNA processing (capping, CTD phosphorylation, promoter escape/elongation), coupled to Unfolded Protein Response modules (PERK–eIF2α/ATF4 program and XBP1 arm) and NRF2-driven oxidative-stress detoxification. Together these drive amino-acid biosynthesis/transport (notably serine–glycine axis), tRNA charging, and ER→Golgi/intra-Golgi trafficking to restore proteostasis. Mitochondrial protein targeting supports bioenergetic needs during stress adaptation. The same network integrates Rab-regulated membrane traffic and vesicle tethering to sustain secretory throughput while damaged RNAs/proteins are processed. The breadth and depth of these categories across the gene list point to a unified “integrated stress response” that rewires metabolism and secretory logistics under proteotoxic/oxidative stress.

##### 1. RNAPII transcription cycle & RNA processing

Enrichment for Pol II initiation, CTD phosphorylation, promoter escape/elongation, and mRNA capping/early elongation indicates a global reset of gene expression needed to implement stress programs. Concurrent ncRNA/RNA metabolic processes and cytosolic tRNA aminoacylation align translation capacity with the new transcriptome.

##### 2. UPR/ISR signaling

*PERK–eIF2α–ATF4* terms (“*ATF4* activates genes in response to ER stress”, “PERK regulates gene expression”, “Unfolded Protein Response”) and *XBP1* highlight a canonical

ER-stress module that attenuates translation while transcriptionally inducing chaperones, *ERAD*, and amino-acid/antioxidant pathways.

#### 3. Antioxidant/detoxification control

“*NFE2L2* regulating anti-oxidant/detoxification enzymes” plus “response to oxidative stress” connect *ROS* buffering directly to the *UPR*, stabilizing redox during increased folding load. [1][2][3]

---


[1] DeNicola, Gina M, Chen, Pei-Hsuan, Mullarky, Edouard, Sudderth, Jessica A, Hu, Zeping, Wu, David, Tang, Hao, Xie, Yang, Asara, John M, Huffman, Kenneth E, Wistuba, Ignacio I, Minna, John D, DeBerardinis, Ralph J, Cantley, Lewis C. ""NRF2 regulates serine biosynthesis in non-small cell lung cancer."" Nature genetics, 2015, pp. 1475-81, doi: https://doi.org/10.1038/ng.3421

[2] Zhang, Cheng, Yu, Jiao-Jiao, Yang, Chen, Yuan, Zhen-Long, Zeng, Hui, Wang, Jun-Jian, Shang, Shuang, Lv, Xiao-Xi, Liu, Xiao-Tong, Liu, Jing, Xue, Qi, Cui, Bing, Tan, Feng-Wei, Hua, Fang. ""Wild-type IDH1 maintains NSCLC stemness and chemoresistance through activation of the serine biosynthetic pathway."" Science translational medicine, 2023, pp. eade4113, doi: https://doi.org/10.1126/scitranslmed.ade4113

[3] Possemato, Richard, Marks, Kevin M, Shaul, Yoav D, Pacold, Michael E, Kim, Dohoon, Birsoy, Kıvanç, Sethumadhavan, Shalini, Woo, Hin-Koon, Jang, Hyun G, Jha, Abhishek K, Chen, Walter W, Barrett, Francesca G, Stransky, Nicolas, Tsun, Zhi-Yang, Cowley, Glenn S, Barretina, Jordi, Kalaany, Nada Y, Hsu, Peggy P, Ottina, Kathleen, Chan, Albert M, Yuan, Bingbing, Garraway, Levi A, Root, David E, Mino-Kenudson, Mari, Brachtel, Elena F, Driggers, Edward M, Sabatini, David M. ""Functional genomics reveal that the serine synthesis pathway is essential in breast cancer."" Nature, 2011, pp. 346-50, doi: https://doi.org/10.1038/nature10350


#### 4. Amino-acid supply & one-carbon metabolism

Strong representation of serine/glycine biosynthesis (*PHGDH–PSAT1–PSPH*) and broader amino-acid biosynthetic/transport terms ensures substrates for glutathione synthesis and stress-induced protein turnover; amino-acid transporters couple uptake with synthesis.

#### 5. Secretory pathway throughput

ER→Golgi anterograde transport, intra-Golgi/retrograde traffic, *Rab-GEF*-regulated trafficking, and N-linked glycosylation maintain folding/quality control and receptor/cargo flow as proteostasis recovers.

##### 6. Mitochondrial targeting & metabolic support

Protein targeting to mitochondria and mitochondrial gene-expression/assembly terms underpin ATP/redox demands of UPR and biosynthesis, linking organelle communication to stress adaptation.

##### 7. Integration with immune/stress sensors

"Immune system process" and "cellular responses to stimuli" categories position *ISR*/*UPR* outputs within broader stress and cytokine contexts without diluting the central proteostasis–metabolism axis.

Most enriched categories and many list genes fall within UPR/ISR signaling, antioxidant control, amino-acid metabolism/transport, and ER–Golgi trafficking, with RNAPII machinery enabling the program. Given the proportion of the set mapping to these nodes, an integrated-stress-response label best captures the predominant, connected function. Confidence reflects broad coverage across these modules.

### ILF2 K562

#### Process: Coupled RNA processing–nuclear export aligned with mitotic cell-cycle control (0.86)

The enrichment profile centers on gene-expression machinery from transcription through post-transcriptional maturation and transport, tightly linked to nuclear pore dynamics and M-phase checkpoints. Splicing factors, RNA-modifying/processing enzymes, mRNP assembly and export components co-occur with nucleoporins and import/export factors, while APC/C–checkpoint modules and kinetochore organizers align mitotic progression with nuclear envelope/pore remodeling. Secretory pathway and UPR nodes provide trafficking/stress capacity that supports high-throughput RNA–protein complex flow. Collectively, the network implements end-to-end RNA biogenesis, quality control, and nucleocytoplasmic transport, coupled to cell-cycle transitions and membrane trafficking.

##### 1. Transcription initiation and elongation across Pol I/II/III

Core transcription modules (*Pol I* promoter clearance/termination; *Pol II* pre-initiation, promoter escape/termination; *Pol III* initiation/elongation) are enriched, anchoring RNA synthesis for rRNA, mRNA, and tRNA. This places the system at the headwaters of gene expression and rRNA/tRNA supply.

##### 2. Co-transcriptional RNA maturation: splicing, 3′-end formation, RNA modification

Strong signals for major-spliceosome splicing (including *snRNP* assembly and bulged-adenosine transesterification), mRNA 3′-end processing, tRNA processing, and broad RNA/RNP biogenesis indicate dense co-transcriptional coupling and surveillance. RNA catabolism and “metabolism of non-coding RNA” further emphasize quality control and turnover.

##### 3. mRNP assembly, nuclear export, and nucleocytoplasmic transport

Enrichment for *mRNP* organization/assembly, RNA/mRNA export, and nucleocytoplasmic transport—together with “structural constituent of nuclear pore” and NPC disassembly/reformation—shows that mature RNPs are packaged and routed across dynamic pores, including dedicated routes for intronless transcripts.

##### 4. Nuclear envelope/pore remodeling synchronized with mitosis

Terms covering mitotic sister-chromatid segregation, kinetochore organization, spindle checkpoint amplification, *APC/C* activation/inactivation, and nuclear envelope breakdown/reassembly indicate tight alignment of genome partitioning and *NPC* cycling so that transport and transcriptional states reset each division.

##### 5. Post-transcriptional regulators and chromatin/epigenetic control

“Chromatin organization/modifying enzymes,” positive/negative epigenetic regulation of rRNA, and “transcriptional regulation by small RNAs/Gene silencing by RNA” integrate chromatin state and small-RNA pathways with RNA output and ribosome supply. *SUMOylation* of RNA-binding, chromatin, replication, and *DDR* proteins adds reversible control across these layers.

##### 6. Secretory and membrane-traffic support + stress responses

*COPI/ER-Golgi* trafficking, Golgi organization and vesicle transport, plus *UPR/HSF1*-heat-shock responses, furnish capacity for processing/trafficking RNA-binding proteins and pore/coat components and buffer proteostasis during bursts of mRNP production and cell-cycle transitions.

##### 7. Gene-set composition is concordant with these functions

The set includes numerous spliceosome/RNA-processing factors (*SF3B1*, *SRSF1/6*, *PRPF3*, *BCAS2*, *AQR*), RNA enzymes/modifiers (*RNASEH1*, *ADAR*, *XRN2*, *PRMT5*), transcription subunits (*POLR1C*, *POLR2H*, *POLR3C*, *TAF1A/11/2/9B*, *GTF3C2*), mRNA export/transport factors (*NXF1*, *XPO5*), nucleoporins (*NUP107/133/155/188/205/88*), and mitotic checkpoint/APC-axis components (*CDC20*, *ANAPC7*, *KNSTRN*, *MIS12*, *CENPL*, *NEK2*, *SEPT2*), together with ER–Golgi coat/traffic proteins (*SEC23B*, *SEC31A*, *COPB1/COPG1/ARCN1*, *COG2*, *TMED2*, *GORAB*) and UPR nodes (*ATF6*, *HYOU1*, *DNAJB11*). This breadth supports an integrated “RNA biogenesis–NPC–mitosis–trafficking” program.

Why this is the most prominent process and confidence

Most enriched terms cluster within RNA biogenesis/processing, RNP assembly/export, and NPC/mitotic regulation, and a large fraction of listed genes map to these modules. Adjacent pathways (chromatin, SUMOylation, ER-Golgi, stress) act as supporting infrastructure rather than alternative cores. Hence a high confidence designation.

-- Summary --

The proposed process name, **“Coupled RNA processing–nuclear export aligned with mitotic cell-cycle control,”** is strongly supported by genes involved in RNA splicing, mRNA transport, nucleoporin structure, and mitotic regulation. Splicing and RNA-binding proteins (e.g., *SF3B1*, *SRSF1*, *PRPF3*), export factors (*NXF1*, *XPO5*), nuclear pore components (*NUP107*, *NUP155*), and cell-cycle checkpoint proteins (*CDC20*, *ANAPC7*, *CENPL*, *KNSTRN*, *NEK2*) are definitively described in the text as integral to these coupled processes.

-- Explanation --

**SF3B1:** [High] Identified as a spliceosomal factor central to major-spliceosome assembly, providing clear support for RNA processing.

**SRSF1 / SRSF6:** [High] Described as core serine/arginine-rich splicing regulators ensuring accurate intron removal and RNA maturation.

**PRPF3:** [High] Labeled a spliceosome component essential for pre-mRNA splicing, linking to co-transcriptional RNA processing.

**BCAS2:** [High] Cited among spliceosome/RNA-processing factors coordinating mRNA splicing, directly reinforcing RNA processing.

**ADAR:** [High] Defined as an RNA-modifying enzyme that edits and processes transcripts, coupling RNA modification to processing.

**RNASEH1 / XRN2:** [High] Indicated as RNA-catabolic enzymes maintaining RNA surveillance, confirming quality-control linkage.

**POLR2H / POLR1C / POLR3C:** [High] Stated as core subunits of RNA polymerases I–III driving transcription initiation and elongation.

**TAF1A / TAF9B / GTF3C2:** [High] Recognized transcription factors forming part of the Pol I/II/III machinery that initiate RNA synthesis.

**NXF1 / XPO5:** [High] Explicitly cited as mRNA export and transport factors mediating nucleocytoplasmic trafficking.

**NUP107 / NUP133 / NUP155 / NUP188 / NUP205 / NUP88:** [High] Directly described as structural constituents of the nuclear pore complex enabling mRNP transport.

**CDC20 / ANAPC7:** [High] Named components of the anaphase-promoting complex (APC/C), which drives mitotic progression and ensures synchrony with nuclear envelope remodeling.

**CENPL / MIS12 / KNSTRN / NEK2 / SEPT2:** [High] Identified as kinetochore and spindle assembly regulators ensuring chromosomal segregation and mitotic checkpoint control.

**SEC23B / SEC31A / COPB1 / COPG1 / ARCN1:** [Medium] Mentioned as COPI/ER–Golgi trafficking proteins supporting RNA-binding protein transport and proteostasis.

**ATF6 / HYOU1 / DNAJB11:** [Medium] Defined as UPR regulators that stabilize protein homeostasis during intensive RNA–protein synthesis.

-- genes supporting the name:

SF3B1, SRSF1, SRSF6, PRPF3, BCAS2, ADAR, RNASEH1, XRN2, POLR2H, POLR1C, POLR3C, TAF1A, TAF9B, GTF3C2, NXF1, XPO5, NUP107, NUP133, NUP155, NUP188, NUP205, NUP88, CDC20, ANAPC7, CENPL, MIS12, KNSTRN, NEK2, SEPT2, SEC23B, SEC31A, COPB1, COPG1, ARCN1, ATF6, HYOU1, DNAJB11

### ILF2 NPC

#### Process: RNAP II/III–coupled RNA biogenesis and ribosome assembly (0.88)

The enriched categories cohere into a continuous gene-expression pipeline: RNAP II/III transcription and pre-initiation/elongation modules feed co-transcriptional RNA processing (capping/splicing/3′ end formation), followed by nuclear export, rRNA/tRNA modification and processing, ribosomal subunit biogenesis, and tRNA aminoacylation. Supporting layers include snRNP assembly, miRNA pathway components, RNA surveillance/decay, chromatin remodeling, and DNA repair. The genelist is dense for Pol subunits/factors, spliceosome/exosome components, rRNA/tRNA maturation enzymes, ribosome assembly factors, and aa-tRNA synthetases—indicating an integrated, RNA-centric biosynthetic network that culminates in ribosome production and translational competence.

##### 1. Transcription initiation and elongation

RNAP II/III pre-initiation, promoter opening, escape, and elongation are prominent (“RNA Polymerase II/III Transcription … Initiation/Elongation/Termination”), linking DNA-templated transcription to nascent RNA output. Pol subunits and general factors (*POLR2H*/*POLR2D*, *POLR3C*/*POLR3D*, *GTF2H4*, *ELOA*, *TAF2/5L*) and cofactors (*MED1*, *RBBP5*, *KDM1A*) anchor this layer.

##### 2. Co-transcriptional RNA processing and RNP assembly

Major/minor spliceosome, snRNP assembly, and “Processing of capped intron-containing pre-mRNA” are enriched (*SF3B4*/*SF3A1*, *SNRPA1*/*SNRNP70*/*DDX23*, *PRPF19*, *KHDC4*; *CPSF7*/*CSTF2*). miRNA biogenesis/export is represented (*ADAR*, *XPO5*). RNA helicases and binding proteins (*DDX1/8/21*, *GPATCH4*) support spliceosome dynamics. [1]

---

[1] Guo, Yue, Xu, Mingjing, Xue, Hua, Ding, Xiaofan, Wong, Alissa M, Lin, Na, Pu, Dandan, Wong, Aikha M, Wang, Xin, Zhao, Hui, Wong, Nathalie. ""Genome-wide CRISPR screen identifies splicing factor SF3B4 in driving hepatocellular carcinoma."" Science advances, 2025, pp. eadw7181, doi: https://doi.org/10.1126/sciadv.adw7181

##### 3. Nuclear export, surveillance, and decay

“RNA export from nucleus,” “nuclear-transcribed mRNA catabolic process,” and exosome terms align with *NXF1*, *XPO5*, *EXOSC2*, *SMG5*, *CNOT9*—coupling maturation to quality control and turnover. [1][2][3]

---

[1] Viphakone, Nicolas, Sudbery, Ian, Griffith, Llywelyn, Heath, Catherine G, Sims, David, Wilson, Stuart A. ""Co-transcriptional Loading of RNA Export Factors Shapes the Human Transcriptome."" Molecular cell, 2019, pp. 310-323.e8, doi: https://doi.org/10.1016/j.molcel.2019.04.034

[2] Tsoi, Ho, Lau, Chi Kong, Lau, Kwok Fai, Chan, Ho Yin Edwin. ""Perturbation of U2AF65/NXF1-mediated RNA nuclear export enhances RNA toxicity in polyQ diseases."" Human molecular genetics, 2011, pp. 3787-97, doi: https://doi.org/10.1093/hmg/ddr297

[3] Wendt, Lisa, Brandt, Janine, Bodmer, Bianca S, Reiche, Sven, Schmidt, Marie Luisa, Traeger, Shelby, Hoenen, Thomas. ""The Ebola Virus Nucleoprotein Recruits the Nuclear RNA Export Factor NXF1 into Inclusion Bodies to Facilitate Viral Protein Expression."" Cells, 2020, pp. , doi: https://doi.org/10.3390/cells9010187

##### 4. rRNA processing and ribosome biogenesis

Strong enrichment for “rRNA processing,” “ribosomal small subunit biogenesis,” and “ribonucleoprotein complex biogenesis” maps to nucleolar factors (*DKC1*, *NOP2*, *BYSL*, *UTP4/6*, *NOB1*, *WDR12*, *LAS1L*, *LSG1*) and multiple *MRPL*/*MRPS*/*RPL*/*RPS* proteins, driving *SSU*/*LSU* assembly and export. [1][2][3]

---

[1] Zemp, Ivo, Wandrey, Franziska, Rao, Sanjana, Ashiono, Caroline, Wyler, Emanuel, Montellese, Christian, Kutay, Ulrike. ""CK1δ and CK1ε are components of human 40S subunit precursors required for cytoplasmic 40S maturation."" Journal of cell science, 2014, pp. 1242-53, doi: https://doi.org/10.1242/jcs.138719

[2] Hölzel, Michael, Rohrmoser, Michaela, Schlee, Martin, Grimm, Thomas, Harasim, Thomas, Malamoussi, Anastassia, Gruber-Eber, Anita, Kremmer, Elisabeth, Hiddemann, Wolfgang, Bornkamm, Georg W, Eick, Dirk. ""Mammalian WDR12 is a novel member of the Pes1-Bop1 complex and is required for ribosome biogenesis and cell proliferation."" The Journal of cell biology, 2005, pp. 367-78, doi: https://doi.org/10.1083/jcb.200501141

[3] Castle, Christopher D, Cassimere, Erica K, Denicourt, Catherine. ""LAS1L interacts with the mammalian Rix1 complex to regulate ribosome biogenesis."" Molecular biology of the cell, 2012, pp. 716-28, doi: https://doi.org/10.1091/mbc.E11-06-0530

#### 5. tRNA metabolism and charging

“tRNA processing” and “tRNA aminoacylation” plus aa-tRNA synthetases (*IARS*, *HARS*, *MARS*, *VARS*; scaffold *AIMP2*) and modification enzymes (*METTL1*, *OSGEP*) establish a charged tRNA pool that ties transcription to translation. [1][2][3]

---

[1] Lin, Shuibin, Liu, Qi, Lelyveld, Victor S, Choe, Junho, Szostak, Jack W, Gregory, Richard I. ""Mettl1/Wdr4-Mediated m<sup>7</sup>G tRNA Methylome Is Required for Normal mRNA Translation and Embryonic Stem Cell Self-Renewal and Differentiation."" Molecular cell, 2018, pp. 244-255.e5, doi: https://doi.org/10.1016/j.molcel.2018.06.001

[2] Orellana, Esteban A, Liu, Qi, Yankova, Eliza, Pirouz, Mehdi, De Braekeleer, Etienne, Zhang, Wencai, Lim, Jihoon, Aspris, Demetrios, Sendinc, Erdem, Garyfallos, Dimitrios A, Gu, Muxin, Ali, Raja, Gutierrez, Alejandro, Mikutis, Sigitas, Bernardes, Gonçalo J L, Fischer, Eric S, Bradley, Allan, Vassiliou, George S, Slack, Frank J, Tzelepis, Konstantinos, Gregory, Richard I. ""METTL1-mediated m<sup>7</sup>G modification of Arg-TCT tRNA drives oncogenic transformation."" Molecular cell, 2021, pp. 3323-3338.e14, doi: https://doi.org/10.1016/j.molcel.2021.06.031

[3] Dai, Zihao, Liu, Haining, Liao, Junbin, Huang, Cheng, Ren, Xiaoxue, Zhu, Wanjie, Zhu, Shenghua, Peng, Baogang, Li, Shaoqiang, Lai, Jiaming, Liang, Lijian, Xu, Lixia, Peng, Sui, Lin, Shuibin, Kuang, Ming. ""N<sup>7</sup>-Methylguanosine tRNA modification enhances oncogenic mRNA translation and promotes intrahepatic cholangiocarcinoma progression."" Molecular cell, 2021, pp. 3339-3355.e8, doi: https://doi.org/10.1016/j.molcel.2021.07.003

#### 6. Chromatin and repair context

“Chromatin modifying enzymes” and “DNA repair” provide access and genome stability during high transcriptional flux (*KDM1A*/*SETDB1*/*SMARCC1*; *APEX1*/*MSH6*/*RNASEH1*). [1][2]

---

[1] Abraham, Karan J, Khosraviani, Negin, Chan, Janet N Y, Gorthi, Aparna, Samman, Anas, Zhao, Dorothy Y, Wang, Miling, Bokros, Michael, Vidya, Elva, Ostrowski, Lauren A, Oshidari,

Roxanne, Pietrobon, Violena, Patel, Parasvi S, Algouneh, Arash, Singhania, Rajat, Liu, Yupeng, Yerlici, V Talya, De Carvalho, Daniel D, Ohh, Michael, Dickson, Brendan C, Hakem, Razq, Greenblatt, Jack F, Lee, Stephen, Bishop, Alexander J R, Mekhail, Karim. ""Nucleolar RNA polymerase II drives ribosome biogenesis."" Nature, 2020, pp. 298-302, doi: https://doi.org/10.1038/s41586-020-2497-0

[2] Harrington, Elena M, Murphy, James C, Harper, Katherine L, Hayward, Connor, Mottram, Timothy J, Aspden, Julie L, Whitehouse, Adrian. ""EMG1 methyltransferase activity affects ribosome occupancy at KSHV uORFs."" Cell reports, 2025, pp. 115516, doi: https://doi.org/10.1016/j.celrep.2025.115516

Integrated view

Together, these modules describe an RNAP-driven RNA production line—from initiation to RNA processing/export and ribosome/tRNA readiness—optimized for robust gene expression and protein synthesis. The breadth of directly matching categories and gene coverage supports the chosen process.

**Why this score:** A clear majority of genes fall into RNAP II/III transcription, RNA processing/transport, rRNA/tRNA pathways, and ribosome biogenesis; ancillary DNA repair/Chromatin nodes are supportive rather than dominant, justifying a high (but not maximal) confidence.

#### PKM K562

##### Process: Hypoxia/HIF-1–orchestrated metabolic reprogramming with cytoskeletal–membrane remodeling (0.80)

###### 1. Transcriptional hypoxia module setting the program

Enrichment for “Cellular response to hypoxia,” “Hypoxia response via HIF activation,” and “Regulation of gene expression by HIF” defines the organizing signal. This layer is instantiated by HIF pathway components alongside broad RNAPII transcription categories, positioning hypoxia as the upstream driver of downstream metabolic and structural programs.

Key effectors in the gene list include *HIF1A*, *EGLN1*/*EGLN3* (oxygen sensors), *ARNTL2*, and canonical targets (*VEGFA*, *NQO1*), anchoring a bona fide HIF axis. [1][2][3]

---

[1] Zhang, Ruyang, Lai, Linjing, He, Jieyu, Chen, Chao, You, Dongfang, Duan, Weiwei, Dong, Xuesi, Zhu, Ying, Lin, Lijuan, Shen, Sipeng, Guo, Yichen, Su, Li, Shafer, Andrea, Moran, Sebastian, Fleischer, Thomas, Bjaanæs, Maria Moksnes, Karlsson, Anna, Planck, Maria, Staaf, Johan, Helland, Åslaug, Esteller, Manel, Wei, Yongyue, Chen, Feng, Christiani, David C. ""EGLN2 DNA methylation and expression interact with HIF1A to affect survival of early-stage NSCLC."" Epigenetics, 2019, pp. 118-129, doi: https://doi.org/10.1080/15592294.2019.1573066

[2] Jiang, Dong, Qi, Zhi, Xu, Zhi-Ying, Li, Yi-Ran. ""<i>F13B</i> regulates angiogenesis and tumor progression in hepatocellular carcinoma via the HIF-1α/VEGF pathway."" Biomolecules & biomedicine, 2024, pp. 189-209, doi: https://doi.org/10.17305/bb.2024.10794

[3] Xiang, Kun, Ouzhuluobu, , Peng, Yi, Yang, Zhaohui, Zhang, Xiaoming, Cui, Chaoying, Zhang, Hui, Li, Ming, Zhang, Yanfeng, Bianba, , Gonggalanzi, , Basang, , Ciwangsangbu, , Wu, Tianyi, Chen, Hua, Shi, Hong, Qi, Xuebin, Su, Bing. ""Identification of a Tibetan-specific mutation in the hypoxic gene EGLN1 and its contribution to high-altitude adaptation."" Molecular biology and evolution, 2013, pp. 1889-98, doi: https://doi.org/10.1093/molbev/mst090

#### 2. Metabolic rewiring toward glycolysis and redox support

Strong terms for "Glycolysis," "Pentose phosphate pathway," "Pyruvate metabolism," "TCA cycle," and "Integration of energy metabolism" indicate a coordinated shift from oxidative phosphorylation toward glycolytic ATP and NADPH generation under low $O_2$. The gene set maps densely to glycolysis (*HK2*, *PFKP*/*PFKM*, *PFKFB3*/*PFKFB4*, *ALDOC*, *ENO2*/*ENO3*), PPP (*G6PD*, *PGD*, *TKT*), and TCA entry/flux (*PDHA1*, *IDH3A/B/G*, *SDHC*), matching classic HIF-directed metabolic remodeling and redox buffering.

#### 3. Proteostasis and catabolic throughput to sustain the shift

Categories for "Proteasome-mediated ubiquitin-dependent protein catabolic process," "Ubiquitin/proteasome degradation," and broader "Protein metabolic process" suggest elevated turnover to reconfigure enzyme inventories and remove damaged proteins during hypoxia. Corresponding subunits/regulators (*PSMA*/*PSMB* family, *PSMD*, *ADRM1*) appear in the gene list, consistent with increased proteolytic flux. [1][2][3]

---

[1] Qiu, Xiao-Bo, Ouyang, Song-Ying, Li, Chao-Jun, Miao, Shiying, Wang, Linfang, Goldberg, Alfred L. ""hRpn13/ADRM1/GP110 is a novel proteasome subunit that binds the deubiquitinating enzyme, UCH37."" The EMBO journal, 2006, pp. 5742-53, doi: https://doi.org/10.1038/sj.emboj.7601450

[2] Deol, Kirandeep K, Crowe, Sean O, Du, Jiale, Bisbee, Heather A, Guenette, Robert G, Strieter, Eric R. ""Proteasome-Bound UCH37/UCHL5 Debranches Ubiquitin Chains to Promote Degradation."" Molecular cell, 2020, pp. 796-809.e9, doi: https://doi.org/10.1016/j.molcel.2020.10.017

[3] VanderLinden, Ryan T, Hemmis, Casey W, Yao, Tingting, Robinson, Howard, Hill, Christopher P. ""Structure and energetics of pairwise interactions between proteasome subunits RPN2, RPN13, and ubiquitin clarify a substrate recruitment mechanism."" The Journal of biological chemistry, 2017, pp. 9493-9504, doi: https://doi.org/10.1074/jbc.M117.785287

#### 4. Actin–cytoskeleton remodeling and focal adhesion dynamics

Enrichment for “Actin cytoskeleton organization,” “Regulation of actin polymerization/depolymerization,” and “Structural constituent of cytoskeleton” points to motility/adhesion adaptation under hypoxia. The gene list is rich in actin regulators/adhesion scaffolds (*ACTN1*, *VASP*, *TLN1*, *FLII*, *CAPZB*/*CAPZA2*, *CTTN*, *ADD1*), aligning with hypoxia-induced invasive morphology and traction remodeling. [1][2][3]

---

[1] Liu, Zhikui, Wang, Yufeng, Dou, Changwei, Xu, Meng, Sun, Liankang, Wang, Liang, Yao, Bowen, Li, Qing, Yang, Wei, Tu, Kangsheng, Liu, Qingguang. ""Hypoxia-induced up-regulation of VASP promotes invasiveness and metastasis of hepatocellular carcinoma."" Theranostics, 2018, pp. 4649-4663, doi: https://doi.org/10.7150/thno.26789

[2] Zhang, Yixiao, Sun, Lisha, Li, Haonan, Ai, Liping, Ma, Qingtian, Qiao, Xinbo, Yang, Jie, Zhang, Hao, Ou, Xunyan, Wang, Yining, Chen, Guanglei, Xue, Jinqi, Zhu, Xudong, Zhao, Yu,

Yang, Yongliang, Liu, Caigang. ""Binding blockade between TLN1 and integrin β1 represses triple-negative breast cancer."" eLife, 2022, pp. , doi: https://doi.org/10.7554/eLife.68481

[3] Mukherjee, Debarati, Previs, Rebecca A, Haines, Corinne, Al Abo, Muthana, Juras, Patrick K, Strickland, Kyle C, Chakraborty, Binita, Artham, Sandeep, Whitaker, Regina S, Hebert, Katherine, Fontenot, Jake, Patierno, Steven R, Freedman, Jennifer A, Lau, Frank H, Burow, Matthew E, Chang, Ching-Yi, McDonnell, Donald P. ""Targeting CaMKK2 Inhibits Actin Cytoskeletal Assembly to Suppress Cancer Metastasis."" Cancer research, 2023, pp. 2889-2907, doi: https://doi.org/10.1158/0008-5472.CAN-22-1622

#### 5. Vesicle trafficking and membrane remodeling to deliver the phenotype

“Vesicle-mediated transport,” “Membrane trafficking,” ER–Golgi transport, and ESCRT terms support receptor turnover and nutrient-transporter routing during hypoxia. Matching components (*HGS*/*TSG101*, *VPS4A*, *VPS18/35/33B*, *DNM1L*/*DNM2*, *SNX1*, *RAB*-effectors) integrate endocytosis, recycling, and autophagic inputs to reshape surface proteomes (e.g., *GLUT1*/*SLC2A1*, *MCT4*/*SLC16A3*). [1][2][3]

---

[1] Kurten, R C, Eddington, A D, Chowdhury, P, Smith, R D, Davidson, A D, Shank, B B. ""Self-assembly and binding of a sorting nexin to sorting endosomes."" Journal of cell science, 2001, pp. 1743-56, doi: https://doi.org/10.1242/jcs.114.9.1743

[2] Imbastari, Francesca, Dahlmann, Mathias, Sporbert, Anje, Mattioli, Camilla Ciolli, Mari, Tommaso, Scholz, Florian, Timm, Lena, Twamley, Shailey, Migotti, Rebekka, Walther, Wolfgang, Dittmar, Gunnar, Rehm, Armin, Stein, Ulrike. ""MACC1 regulates clathrin-mediated endocytosis and receptor recycling of transferrin receptor and EGFR in colorectal cancer."" Cellular and molecular life sciences : CMLS, 2021, pp. 3525-3542, doi: https://doi.org/10.1007/s00018-020-03734-1

[3] Beyer, Andreas, Scheuring, Sibylle, Müller, Sibylle, Mincheva, Antoaneta, Lichter, Peter, Köhrer, Karl. ""Comparative sequence and expression analyses of four mammalian VPS4 genes."" Gene, 2003, pp. 47-59, doi: https://doi.org/10.1016/s0378-1119(02)01205-2

#### 6. mRNA processing and translation control coupled to stress

Prominent “mRNA splicing/processing” categories indicate transcriptome remodeling to implement the hypoxic state. The set includes spliceosome and RNA-binding factors (*PRPF8*, *SRSF1*/*SRSF9*, *CPSF6*, *DDX41*/*DDX46*, *HNRNPK/U/L2*), supporting co-transcriptional selection of isoforms that favor glycolysis, angiogenesis, and survival. [1][2][3]

---

[1] Das, Shipra, Krainer, Adrian R. ""Emerging functions of SRSF1, splicing factor and oncoprotein, in RNA metabolism and cancer."" Molecular cancer research : MCR, 2014, pp. 1195-204, doi: https://doi.org/10.1158/1541-7786.MCR-14-0131

[2] Matsumoto, Eri, Akiyama, Kaho, Saito, Takuya, Matsumoto, Yu, Kobayashi, Ken-Ichi, Inoue, Jun, Yamamoto, Yuji, Suzuki, Tsukasa. ""AMP-activated protein kinase regulates alternative pre-mRNA splicing by phosphorylation of SRSF1."" The Biochemical journal, 2020, pp. 2237-2248, doi: https://doi.org/10.1042/BCJ20190894

[3] Wu, Qikai, Yu, Hao, Sun, Huanyou, Lv, Jiancheng, Zhuang, Juntao, Cai, Lingkai, Jiang, Lingjing, Chen, Yuhan, Tao, Yiran, Bai, Kexin, Yang, Haiwei, Yang, Xiao, Lu, Qiang. ""SRSF1-mediated alternative splicing regulates bladder cancer progression and cisplatin sensitivity through HIF1A/BNIP3/mitophagy axis."" Journal of translational medicine, 2025, pp. 571, doi: https://doi.org/10.1186/s12967-025-06547-7

#### 7. KEAP1–NFE2L2 (NRF2)–linked oxidative stress buffering (adjacent)

Pathway terms for the *KEAP1–NFE2L2* axis and “Cellular response to chemical stress” complement HIF by boosting antioxidant and NADPH-producing routes (PPP), stabilizing redox homeostasis during hypoxia. Gene-level support includes PPP enzymes (*G6PD*, *PGD*, *TKT*) and stress chaperones/Detox enzymes (*PRDX3*, *TXN*). [1][2][3]

---

[1] Hayes, John D, Dinkova-Kostova, Albena T. ""The Nrf2 regulatory network provides an interface between redox and intermediary metabolism."" Trends in biochemical sciences, 2014, pp. 199-218, doi: https://doi.org/10.1016/j.tibs.2014.02.002

[2] Yamamoto, Masayuki, Kensler, Thomas W, Motohashi, Hozumi. ""The KEAP1-NRF2 System: a Thiol-Based Sensor-Effector Apparatus for Maintaining Redox Homeostasis."" Physiological reviews, 2018, pp. 1169-1203, doi: https://doi.org/10.1152/physrev.00023.2017

[3] Bellezza, Ilaria, Giambanco, Ileana, Minelli, Alba, Donato, Rosario. ""Nrf2-Keap1 signaling in oxidative and reductive stress."" Biochimica et biophysica acta. Molecular cell research, 2018, pp. 721-733, doi: https://doi.org/10.1016/j.bbamcr.2018.02.010

Integrated model

A HIF-centered transcriptional program coordinates a shift to glycolytic/PPP metabolism and redox control, while retooling cytoskeletal and membrane-trafficking modules to adapt adhesion, transport, and signaling. Splicing/translation factors implement the new gene expression state; proteasome/autophagy clear conflicting components—together forming a coherent hypoxia-adaptation network.

Why this name and score

The enriched categories with the densest gene coverage are hypoxia/HIF signaling, glycolysis/PPP/TCA rewiring, and actin–membrane remodeling, each strongly represented in the gene list; RNA processing and proteolysis support but are not the central output. Hence a high confidence that hypoxia/HIF-driven metabolic and structural adaptation is the predominant process.

-- Summary --

The proposed name **"Hypoxia/HIF-1–orchestrated metabolic reprogramming with cytoskeletal–membrane remodeling"** is strongly supported by genes clearly linked in the text to hypoxia response (HIF1A, EGLN1, EGLN3, VEGFA, NQO1), glycolytic and redox pathways (HK2, PFKP, PFKM, PFKFB3, PFKFB4, ALDOC, ENO2, ENO3, G6PD, PGD, TKT, PDHA1, IDH3A/B/G, SDHC), proteasome-mediated protein turnover (PSMA/PSMB/PSMD genes), and cytoskeletal/membrane dynamics (ACTN1, TLN1, CAPZA2, CTTN, VASP, FLII, ADD1, HGS, TSG101, VPS4A). These genes collectively embody the HIF-driven adaptation of energy metabolism, redox maintenance, and structural remodeling described in the analysis.

-- Explanation --

**HIF1A:** [High] Directly stated as the central transcription factor coordinating hypoxia responses and metabolic adaptation.

**EGLN1 / EGLN3:** [High] Defined as oxygen sensors that regulate HIF stability and thereby control transcriptional adaptation to low $O_2$.

**VEGFA:** [High] Explicitly noted as a canonical HIF target involved in angiogenesis and hypoxic signaling.

**NQO1:** [High] Listed as a HIF-induced gene contributing to redox regulation under hypoxia.

**HK2, PFKP, PFKM, PFKFB3, PFKFB4, ALDOC, ENO2, ENO3:** [High] Cited as glycolytic enzymes activated under hypoxia, driving the metabolic shift from oxidative phosphorylation to glycolysis.
**G6PD, PGD, TKT:** [High] Named as core pentose phosphate pathway enzymes supplying NADPH and redox buffering capacity during hypoxia.
**PDHA1, IDH3A, IDH3B, IDH3G, SDHC:** [Medium] Identified as TCA cycle components involved in partial metabolic rewiring to maintain redox balance.
**PSMA/PSMB/PSMD family, ADRM1:** [Medium] Defined as proteasome subunits ensuring protein quality control and turnover to sustain hypoxic remodeling.
**ACTN1, TLN1, CAPZA2, CTTN, VASP, FLII, ADD1:** [High] Highlighted as actin-cytoskeleton and focal adhesion regulators, mediating structural and motility changes under hypoxia.
**HGS, TSG101, VPS4A:** [Medium] Assigned to vesicle trafficking and ESCRT machinery that remodels membranes and receptor transport during hypoxia.
**PRPF8, SRSF1, SRSF9, CPSF6, DDX46:** [Medium] Stated as splicing and RNA-processing factors adjusting transcript isoforms to support the hypoxic phenotype.
**KEAP1, NFE2L2, TXN, PRDX3:** [Medium] Described in the KEAP1–NFE2L2 antioxidant module buffering oxidative stress complementary to HIF signaling.

-- genes supporting the name:
HIF1A, EGLN1, EGLN3, VEGFA, NQO1, HK2, PFKP, PFKM, PFKFB3, PFKFB4, ALDOC, ENO2, ENO3, G6PD, PGD, TKT, PDHA1, IDH3A, IDH3B, IDH3G, SDHC, PSMA family, PSMB family, PSMD family, ADRM1, ACTN1, TLN1, CAPZA2, CTTN, VASP, FLII, ADD1, HGS, TSG101, VPS4A, PRPF8, SRSF1, SRSF9, CPSF6, DDX46, KEAP1, NFE2L2, TXN, PRDX3

### PKM NPC

#### Process: Hypoxia-driven metabolic rewiring with KEAP1–NRF2 signaling and actin–endosomal remodeling (0.86)

A transcription-independent stress program links oxygen sensing to energy and membrane-traffic control. Enriched categories converge on glycolysis/PPP, TCA/pyruvate regulation, and “cellular response to hypoxia,” alongside KEAP1–NFE2L2 signaling, endocytosis, vesicle/lysosome acidification, integrin/actin cytoskeleton organization, proteasome-mediated turnover, and selective autophagy. Together these modules promote a high-flux glycolytic state, buffer redox stress, and remodel adhesion/trafficking to sustain survival under low $O_2$.

#### 1. Metabolic rewiring and hypoxia response

Carbohydrate metabolism, glycolysis/pyruvate/TCA terms and “cellular response to hypoxia” dominate the enrichment. The gene set contains canonical HIF/oxygen-sensing and glycolytic/PPP executors (*EGLN1*/*EGLN3*; *SLC2A1*/*SLC2A3*; *HK1*, *ALDOC*, *PFKP*, *PFKFB3/4*; *G6PD*, *TKT*), and lactate/pyruvate handling (*PDK1*/*PDK3*) that reallocate carbon from OXPHOS to glycolysis while maintaining anabolic NADPH. Redox enzymes (*NQO1*) and mitochondrial enzymes (*SDHA*/*SDHC*, *IDH3A/B/G*, *OGDH*, *CS*) illustrate coordinated down-tuning of respiration and reinforcement of antioxidant capacity. [1][2][3]

---

[1] Zodda, Erika, Tura-Ceide, Olga, Mills, Nicholas L, Tarragó-Celada, Josep, Carini, Marina, Thomson, Timothy M, Cascante, Marta. ""Autonomous metabolic reprogramming and oxidative stress characterize endothelial dysfunction in acute myocardial infarction."" eLife, 2023, pp. , doi: https://doi.org/10.7554/eLife.86260

[2] Wang, Tong, Wang, Xueting, Zheng, Xuli, Guo, Zhongfang, Mohsin, Ali, Zhuang, Yingping, Wang, Guan. ""Overexpression of SLC2A1, ALDOC, and PFKFB4 in the glycolysis pathway drives strong drug resistance in 3D HeLa tumor cell spheroids."" Biotechnology journal, 2024, pp. e2400163, doi: https://doi.org/10.1002/biot.202400163

[3] Massari, Francesco, Ciccarese, Chiara, Santoni, Matteo, Iacovelli, Roberto, Mazzucchelli, Roberta, Piva, Francesco, Scarpelli, Marina, Berardi, Rossana, Tortora, Giampaolo, Lopez-Beltran, Antonio, Cheng, Liang, Montironi, Rodolfo. ""Metabolic phenotype of bladder cancer."" Cancer treatment reviews, 2016, pp. 46-57, doi: https://doi.org/10.1016/j.ctrv.2016.03.005

#### 2. KEAP1–NRF2 cytoprotective axis

*KEAP1–NFE2L2* pathway enrichment aligns with *KEAP1* together with *NRF2* targets involved in pentose-phosphate/NADPH production (*G6PD*, *TKT*) and xenobiotic defense (*NQO1*, *ABC* transporters). This axis integrates chemical/oxidative stress with the hypoxic metabolic shift, stabilizing redox homeostasis during sustained glycolytic flux. [1][2][3]

---

[1] Li, Dan, Shao, Rong, Wang, Na, Zhou, Nan, Du, Kaili, Shi, Jiahui, Wang, Yihan, Zhao, Zhuangzhuang, Ye, Xin, Zhang, Xiaoli, Xu, Haoxing. ""Sulforaphane Activates a lysosome-dependent transcriptional program to mitigate oxidative stress."" Autophagy, 2021, pp. 872-887, doi: https://doi.org/10.1080/15548627.2020.1739442

[2] Hayes, John D, Dinkova-Kostova, Albena T. ""The Nrf2 regulatory network provides an interface between redox and intermediary metabolism."" Trends in biochemical sciences, 2014, pp. 199-218, doi: https://doi.org/10.1016/j.tibs.2014.02.002

[3] Yamamoto, Masayuki, Kensler, Thomas W, Motohashi, Hozumi. ""The KEAP1-NRF2 System: a Thiol-Based Sensor-Effector Apparatus for Maintaining Redox Homeostasis."" Physiological reviews, 2018, pp. 1169-1203, doi: https://doi.org/10.1152/physrev.00023.2017

##### 3. Actin/adhesion remodeling and endocytic–lysosomal traffic

Terms for integrin signaling, actin filament organization, clathrin-mediated endocytosis, vesicle-mediated transport, and V-ATPase-dependent acidification are jointly enriched. The gene set encodes actin nucleation/regulation and adhesion adaptors (*ARPC2*, *ACTR2/3*, *CAPZB*, *VASP*, *TLN1*, *ILK*, *VCL*), endocytic machinery (*DNM2*, *SNX1*, *VPS35*), and proton-pump/lysosome factors (*ATP6V1A/B2/E1*, *ATP6V0D1*, *LAPTM4B*, *M6PR*) that rewire receptor turnover and nutrient acquisition under stress. [1][2][3]

---

[1] Kurten, R C, Eddington, A D, Chowdhury, P, Smith, R D, Davidson, A D, Shank, B B. ""Self-assembly and binding of a sorting nexin to sorting endosomes."" Journal of cell science, 2001, pp. 1743-56, doi: https://doi.org/10.1242/jcs.114.9.1743

[2] Liang, Xue-Hai, Sun, Hong, Hsu, Chih-Wei, Nichols, Joshua G, Vickers, Timothy A, De Hoyos, Cheryl L, Crooke, Stanley T. ""Golgi-endosome transport mediated by M6PR facilitates release of antisense oligonucleotides from endosomes."" Nucleic acids research, 2020, pp. 1372-1391, doi: https://doi.org/10.1093/nar/gkz1171

[3] Böttcher, Ralph T, Lange, Anika, Fässler, Reinhard. ""How ILK and kindlins cooperate to orchestrate integrin signaling."" Current opinion in cell biology, 2009, pp. 670-5, doi: https://doi.org/10.1016/j.ceb.2009.05.008

#### 4. Proteostasis, autophagy, and apoptosis gating

Ubiquitin–proteasome and selective autophagy pathways appear with numerous core subunits (*PSMA*/*PSMB*/*PSMC*, *OTUB1*) and ATG7, supporting removal of damaged proteins/organelles generated by hypoxia. Parallel *TP53*/checkpoint and caspase-execution annotations indicate apoptosis control; *BNIP3*/*BNIP3L* and DDIT4 (mTOR restraint) provide a switch between survival autophagy and death when stress exceeds capacity. [1][2][3]

---

[1] Zhang, J, Ney, P A. ""Role of BNIP3 and NIX in cell death, autophagy, and mitophagy."" Cell death and differentiation, 2009, pp. 939-46, doi: https://doi.org/10.1038/cdd.2009.16

[2] Li, Lulu, Tan, Jin, Miao, Yuyang, Lei, Ping, Zhang, Qiang. ""ROS and Autophagy: Interactions and Molecular Regulatory Mechanisms."" Cellular and molecular neurobiology, 2015, pp. 615-21, doi: https://doi.org/10.1007/s10571-015-0166-x

[3] Sulkshane, Prasad, Ram, Jonathan, Thakur, Anita, Reis, Noa, Kleifeld, Oded, Glickman, Michael H. ""Ubiquitination and receptor-mediated mitophagy converge to eliminate oxidation-damaged mitochondria during hypoxia."" Redox biology, 2021, pp. 102047, doi: https://doi.org/10.1016/j.redox.2021.102047

Integrated model

Under low $O_2$, oxygen sensors (EGLN1/3) and NRF2 signaling drive a glycolytic/PPP shift, while endocytosis–lysosome and actin/adhesion systems remodel surface receptor trafficking and matrix contacts. Proteostasis and autophagy maintain viability; if stress persists, apoptosis executes. The density of hypoxia/metabolic, NRF2/redox, and cytoskeleton/endocytosis categories across the gene list justifies the chosen name and high confidence.

-- Summary --

The suggested name **“Hypoxia-driven metabolic rewiring with KEAP1–NRF2 signaling and actin–endosomal remodeling”** is supported by multiple genes that are clearly described as executing hypoxia response, glycolytic/PPP metabolism, oxidative stress defense, or cytoskeletal and vesicular remodeling. Key examples include oxygen sensors

(EGLN1, EGLN3), glycolytic enzymes (HK1, PFKP, PFKFB3, PFKFB4, ALDOC), redox regulators (G6PD, TKT, NQO1), KEAP1–NRF2 modulators, and structural/trafficking proteins (ARPC2, ACTR2/3, TLN1, ILK, VCL, VPS35, ATP6V1 subunits). These collectively represent the biological modules described in the analysis text.

-- Explanation --

**EGLN1 / EGLN3:** [High] Explicitly identified as oxygen-sensing enzymes that regulate the hypoxia response through HIF signaling, initiating the transcriptional hypoxia program.

**HK1, PFKP, PFKFB3, PFKFB4, ALDOC:** [High] Cited as key glycolytic enzymes mediating the metabolic shift from oxidative phosphorylation to glycolysis under hypoxic conditions.

**G6PD / TKT:** [High] Described as core pentose phosphate pathway enzymes producing NADPH, directly supporting redox homeostasis via the KEAP1–NRF2 axis.

**NQO1:** [High] Defined as an NRF2 target gene involved in antioxidant defense, linking metabolic and redox regulation.

**SLC2A1 / SLC2A3:** [High] Mentioned as glucose transporters activated in hypoxia, supplying substrates for glycolysis.

**PDK1 / PDK3:** [High] Identified as pyruvate dehydrogenase kinases redirecting carbon flow from the TCA cycle toward glycolysis.

**KEAP1:** [High] Stated as a regulatory node of the KEAP1–NRF2 cytoprotective pathway, coordinating redox control and metabolic adaptation.

**SDHA / SDHC / IDH3A / IDH3B / IDH3G / CS / OGDH:** [Medium] Named as mitochondrial/TCA enzymes contributing to respiratory modulation during metabolic rewiring.

**ARPC2 / ACTR2 / ACTR3 / CAPZB / TLN1 / ILK / VCL / VASP / FLII / ADD1:** [High] Highlighted as actin cytoskeleton regulators and adhesion scaffolds facilitating cellular remodeling and integrin signaling under stress.

**DNM2 / SNX1 / VPS35 / M6PR / ATP6V1A / ATP6V1B2 / ATP6V1E1 / ATP6V0D1:** [Medium] Described as endocytic and lysosomal transport proteins enabling vesicle-mediated nutrient uptake and receptor recycling.

**PSMA / PSMB / PSMC / OTUB1 / ATG7:** [Medium] Represent proteasome and autophagy factors maintaining proteostasis during hypoxia-induced stress.

**BNIP3 / BNIP3L / DDIT4:** [Medium] Explicitly linked to autophagy/apoptosis regulation and mTOR inhibition in prolonged hypoxia.

-- genes supporting the name:

EGLN1, EGLN3, HK1, PFKP, PFKFB3, PFKFB4, ALDOC, G6PD, TKT, NQO1, SLC2A1, SLC2A3, PDK1, PDK3, KEAP1, SDHA, SDHC, IDH3A, IDH3B, IDH3G, CS, OGDH, ARPC2, ACTR2, ACTR3, CAPZB, TLN1, ILK, VCL, VASP, FLII, ADD1, DNM2, SNX1, VPS35, M6PR,

ATP6V1A, ATP6V1B2, ATP6V1E1, ATP6V0D1, PSMA family, PSMB family, PSMC family, OTUB1, ATG7, BNIP3, BNIP3L, DDIT4

### POLR2G K562

#### Process: RNA processing–coupled ribosome biogenesis and ER-targeted translation (0.90)

The enrichment profile centers on a continuous "gene-expression → translation → targeting" axis. Overrepresented categories include RNA splicing/processing, rRNA/tRNA maturation, ribosome biogenesis, cytoplasmic translation, and SRP-dependent cotranslational targeting, with coordinated modules for proteostasis (ubiquitin–proteasome, ERAD), vesicle transport/ESCRT, and mitochondrial energy/translation supporting high biosynthetic throughput.

##### 1. RNA processing hub feeding translation

Strong enrichment for spliceosome assembly, mRNA 3′-end processing, and RNA decay defines an entry node to the protein-synthesis pipeline. This is executed by spliceosomal/exosomal and RNP assembly factors abundant in the gene list (e.g., *PRPF*/*SNRNP*/*LSM*/*GEMIN*/*EXOSC*/*CPSF*/*CLP1* modules), ensuring production-ready mRNAs for ribosome loading. [1][2]

---


[1] Zaric, Bozidarka L, Kambach, Christian. ""Reconstitution of recombinant human LSm complexes for biochemical, biophysical, and cell biological studies."" Methods in enzymology, 2008, pp. 57-74, doi: https://doi.org/10.1016/S0076-6879(08)02604-9

[2] Kaur, Mandeep, Sharma, Akanksha, Singh, Gurpal, Kumar, Santosh, Barnwal, Ravi P. ""Architectural and functional details of CF IA proteins involved in yeast 3'-end pre-mRNA processing and its significance for eukaryotes: A concise review."" International journal of biological macromolecules, 2021, pp. 387-400, doi: https://doi.org/10.1016/j.ijbiomac.2021.10.129

#### 2. rRNA/tRNA modification and ribosome biogenesis

Categories for rRNA processing, small-subunit biogenesis, tRNA processing/thiolation, and protein–RNA complex organization indicate nucleolar/tRNA pathways (*EMG1*, *RRP8*, *FCF1*, *TSR2*, *NOP10*, *OSGEP*/*URM1*). These enable assembly of functionally competent ribosomes and decoding tRNAs, directly elevating translational capacity. [1]

---

[1] Eschrich, Dietmar, Buchhaupt, Markus, Kötter, Peter, Entian, Karl-Dieter. ""Nep1p (Emg1p), a novel protein conserved in eukaryotes and archaea, is involved in ribosome biogenesis."" Current genetics, 2002, pp. 326-38, doi: https://doi.org/10.1007/s00294-001-0269-4

#### 3. High-capacity cytosolic translation

“Translation,” “peptide biosynthesis,” and “initiation/elongation/termination” are enriched alongside extensive RPS/RPL cohorts and initiation/elongation factors (*EIF2B* family, *EIF3M*, *EIF4E2*, *EEF1G*), reflecting sustained throughput from mature mRNA to polypeptide. [1][2][3]

---

[1] Hassan, Md Khurshidul, Kumar, Dinesh, Naik, Monali, Dixit, Manjusha. ""The expression profile and prognostic significance of eukaryotic translation elongation factors in different cancers."" PloS one, 2018, pp. e0191377, doi: https://doi.org/10.1371/journal.pone.0191377

[2] Gomes-Duarte, Andreia, Lacerda, Rafaela, Menezes, Juliane, Romão, Luísa. ""eIF3: a factor for human health and disease."" RNA biology, 2018, pp. 26-34, doi: https://doi.org/10.1080/15476286.2017.1391437

[3] Goh, S-H, Hong, S-H, Hong, S-H, Lee, B-C, Ju, M-H, Jeong, J-S, Cho, Y-R, Kim, I-H, Lee, Y-S. ""eIF3m expression influences the regulation of tumorigenesis-related genes in human colon cancer."" Oncogene, 2011, pp. 398-409, doi: https://doi.org/10.1038/onc.2010.422

#### 4. SRP-dependent targeting and secretory influx

Enrichment for SRP-dependent cotranslational targeting and ER-to-Golgi transport links ribosomes to the secretory pathway. The gene set includes SRP/ER translocon-adjacent and vesicle-trafficking apparatus (*SSR2*/*SSR4*, *TRAPP* complex, *COPG2*/*COPZ2*, *YIPF1*, *STX5*/*STX8*, *VAMP8*, *RAB*/*ARF GTPases*), coordinating nascent-chain delivery into the endomembrane system. [1][2][3]

[1] Linders, Peter Ta, Horst, Chiel van der, Beest, Martin Ter, van den Bogaart, Geert. ""Stx5-Mediated ER-Golgi Transport in Mammals and Yeast."" Cells, 2019, pp. , doi: https://doi.org/10.3390/cells8080780

[2] Kranjc, Tilen, Dempsey, Eugene, Cagney, Gerard, Nakamura, Nobuhiro, Shields, Denis C, Simpson, Jeremy C. ""Functional characterisation of the YIPF protein family in mammalian cells."" Histochemistry and cell biology, 2017, pp. 439-451, doi: https://doi.org/10.1007/s00418-016-1527-3

[3] Lee, Wei Siang, Tan, Daniel Cs, Deng, Yuanyuan, van Hummel, Annika, Ippati, Stefania, Stevens, Claire, Carmona-Mora, Paulina, Ariawan, Daryl, Hou, Liming, Stefen, Holly, Tomanic, Tamara, Bi, Mian, Tomasetig, Florence, Martin, Adam, Fath, Thomas, Palmer, Stephen, Ke, Yazi D, Ittner, Lars M. ""Syntaxins 6 and 8 facilitate tau into secretory pathways."" The Biochemical journal, 2021, pp. 1471-1484, doi: https://doi.org/10.1042/BCJ20200664

#### 5. Proteostasis and quality control

“Ubiquitin-dependent protein catabolic process,” ERAD, and proteasomal pathways are enriched and represented by multiple core and regulatory proteasome subunits (*PSMA/B/C/D/E*), assembly chaperones (*PSMG3*), and *ESCRT*/*MVB* machinery (*TSG101*, *CHMP4A/5*, *VPS11/16/25/33/41/45*), ensuring turnover of defective RNAs/proteins and membrane protein sorting. [1][2][3]

[1] Xuan, Do Thi Minh, Yeh, I-Jeng, Su, Che-Yu, Liu, Hsin-Liang, Ta, Hoang Dang Khoa, Anuraga, Gangga, Chiao, Chung-Chieh, Wang, Chih-Yang, Yen, Meng-Chi. ""Prognostic and Immune Infiltration Value of Proteasome Assembly Chaperone (PSMG) Family Genes in Lung Adenocarcinoma."" International journal of medical sciences, 2023, pp. 87-101, doi: https://doi.org/10.7150/ijms.78590

[2] Ma, Zhongchen, Li, Ruirui, Hu, Ruirui, Zheng, Wei, Yu, Shuifa, Cheng, Kejian, Zhang, Huan, Xiao, Yangyang, Yi, Jihai, Wang, Zhen, Wang, Yong, Chen, Chuangfu. ""Anaplasma phagocytophilum AptA enhances the UPS, autophagy, and anti-apoptosis of host cells by PSMG3."" International journal of biological macromolecules, 2021, pp. 497-508, doi: https://doi.org/10.1016/j.ijbiomac.2021.06.039

[3] Zheng, Mengli, Chen, Xiaochun, Xu, Zhe, Ye, Zhitao, Zhong, Lixian, Chen, Zhicao, Chen, Guiquan, Cai, Boyong. ""Comprehensive analysis of PSMG3 in pan-cancer and validation of its role in hepatocellular carcinoma."" Clinical & translational oncology : official publication of the Federation of Spanish Oncology Societies and of the National Cancer Institute of Mexico, 2025, pp. 319-332, doi: https://doi.org/10.1007/s12094-024-03580-8

#### 6. Metabolic support: mitochondria and AMPK–mTOR control

“Cellular respiration,” “Complex I biogenesis,” and “mitochondrial translation” are enriched; numerous complex I/III/IV assembly factors (*NDUF**, *COX14/15/16*, *UQCC2/3*), mitoribosomal proteins (*MRPL*/*MRPS*), and import/tethering components (*TIMM10/8B/TIMMDC1*, *FIS1*) supply ATP and signaling to sustain ribosome biogenesis/translation. AMPK (*PRKAB1*/*PRKAG1*), Ragulator (*LAMTOR2/4/5*), and *NPRL2* integrate nutrient cues to modulate initiation. [1][2]

---

[1] Wang, Tao, Liu, Honghe, Itoh, Kie, Oh, Sungtaek, Zhao, Liang, Murata, Daisuke, Sesaki, Hiromi, Hartung, Thomas, Na, Chan Hyun, Wang, Jiou. ""C9orf72 regulates energy homeostasis by stabilizing mitochondrial complex I assembly."" Cell metabolism, 2021, pp. 531-546.e9, doi: https://doi.org/10.1016/j.cmet.2021.01.005

[2] Guarani, Virginia, Paulo, Joao, Zhai, Bo, Huttlin, Edward L, Gygi, Steven P, Harper, J Wade. ""TIMMDC1/C3orf1 functions as a membrane-embedded mitochondrial complex I assembly factor through association with the MCIA complex."" Molecular and cellular biology, 2014, pp. 847-61, doi: https://doi.org/10.1128/MCB.01551-13[1] Wang, Tao, Liu, Honghe, Itoh, Kie, Oh, Sungtaek, Zhao, Liang, Murata, Daisuke, Sesaki, Hiromi, Hartung, Thomas, Na, Chan Hyun, Wang, Jiou. ""C9orf72 regulates energy homeostasis by stabilizing mitochondrial complex I assembly."" Cell metabolism, 2021, pp. 531-546.e9, doi: https://doi.org/10.1016/j.cmet.2021.01.005

[2] Guarani, Virginia, Paulo, Joao, Zhai, Bo, Huttlin, Edward L, Gygi, Steven P, Harper, J Wade. ""TIMMDC1/C3orf1 functions as a membrane-embedded mitochondrial complex I assembly factor through association with the MCIA complex."" Molecular and cellular biology, 2014, pp. 847-61, doi: https://doi.org/10.1128/MCB.01551-13

#### 7. Genome integrity and transcriptional throughput

Concurrent enrichment for “RNA polymerase II transcription,” “transcription initiation,” and DNA repair checkpoints indicates an upstream layer (MED complex, *TAFs*, *POLR2C/3GL*; repair: *RPA3*, *RAD51C*, *DDB2*, *PAXX*, *SHLD1*) that safeguards and drives the RNA supply into the processing–translation conveyor. [1][2]

---

[1] Noordermeer, Sylvie M, Adam, Salomé, Setiaputra, Dheva, Barazas, Marco, Pettitt, Stephen J, Ling, Alexanda K, Olivieri, Michele, Álvarez-Quilón, Alejandro, Moatti, Nathalie, Zimmermann, Michal, Annunziato, Stefano, Krastev, Dragomir B, Song, Feifei, Brandsma, Inger, Frankum, Jessica, Brough, Rachel, Sherker, Alana, Landry, Sébastien, Szilard, Rachel K, Munro, Meagan M, McEwan, Andrea, Goullet de Rugy, Théo, Lin, Zhen-Yuan, Hart, Traver, Moffat, Jason, Gingras, Anne-Claude, Martin, Alberto, van Attikum, Haico, Jonkers, Jos, Lord, Christopher J, Rottenberg, Sven, Durocher, Daniel. ""The shieldin complex mediates 53BP1-dependent DNA repair."" Nature, 2018, pp. 117-121, doi: https://doi.org/10.1038/s41586-018-0340-7

[2] Greenhough, Luke A, Liang, Chih-Chao, Belan, Ondrej, Kunzelmann, Simone, Maslen, Sarah, Rodrigo-Brenni, Monica C, Anand, Roopesh, Skehel, Mark, Boulton, Simon J, West, Stephen C. ""Structure and function of the RAD51B-RAD51C-RAD51D-XRCC2 tumour suppressor."" Nature, 2023, pp. 650-657, doi: https://doi.org/10.1038/s41586-023-06179-1

##### 8. Peripheral modules (contextual but secondary)

Cilium/IFT (*IFT43/46/52*, *BBS4*), immune antigen processing (*B2M*, *PSMB8/10*), and GPCR/ion-transport terms appear but represent smaller fractions relative to the dominant RNA→ribosome→translation→ER axis.

Rationale for process name and score

The most coherent, densely populated signal spans RNA maturation, ribosome assembly, cytosolic translation, and ER targeting, with robust support from proteostasis and mitochondrial energy modules. Given the high proportion of genes mapping to these categories in both enrichment and the gene list, a confidence of 0.90 is warranted.

-- Summary --

The text explicitly describes a coordinated system centered on **RNA processing, ribosome biogenesis, translation, and ER targeting**. Genes directly involved in RNA maturation (e.g., *PRPF19*, *EXOSC1*, *CPSF7*), ribosome assembly (*RRP8*, *EMG1*, *NOP10*), translation

(*RPS11*, *EIF2B1*, *EEF1G*), and protein targeting (*SSR2*, *SSR4*, *VAMP8*) clearly support the name “RNA processing–coupled ribosome biogenesis and ER-targeted translation.”

-- Explanation --

**PRPF19:** [High] Described as a spliceosome factor participating in RNA splicing and mRNA maturation within the RNA processing hub, directly supporting RNA-processing coupling.

**EXOSC1:** [High] Defined as part of the exosome complex involved in RNA decay and maturation, reinforcing the RNA processing axis.

**CPSF7:** [High] Mentioned as a 3′-end processing factor ensuring production-ready mRNA, essential for translation readiness.

**RRP8:** [High] Identified in rRNA processing and ribosomal small subunit biogenesis, contributing to ribosome assembly.

**EMG1:** [High] Functions in rRNA modification during ribosome biogenesis, supporting the ribosome synthesis component.

**NOP10:** [High] Participates in snoRNP-mediated rRNA processing and assembly, central to ribosome maturation.

**RPS11:** [High] A ribosomal small subunit protein directly involved in translation initiation and elongation.

**EIF2B1:** [High] Listed as a translation initiation factor controlling global translation rates.

**EEF1G:** [High] Functions in peptide chain elongation, a core process in cytosolic translation.

**SSR2:** [High] Involved in SRP-dependent cotranslational targeting of nascent proteins to the ER membrane.

**SSR4:** [High] Similarly described as an ER translocon component guiding secretory protein import, supporting ER-targeted translation.

**VAMP8:** [High] Functions in vesicle fusion within ER-to-Golgi trafficking, integrating translation output with the secretory pathway.

**PSMA3:** [Medium] A proteasomal subunit maintaining protein quality control downstream of translation, indirectly supporting the process.

**NDUFB2:** [Medium] A mitochondrial complex I factor contributing to energy supply for translation and ribosome biogenesis.

-- genes supporting the name:
PRPF19, EXOSC1, CPSF7, RRP8, EMG1, NOP10, RPS11, EIF2B1, EEF1G, SSR2, SSR4, VAMP8, PSMA3, NDUFB2

### POLR2G NPC

#### Process: RNA processing–coupled cytosolic and mitochondrial translation (0.90)

The enrichment profile is dominated by translation (initiation/elongation/termination), ribosome biogenesis, rRNA processing/modification, and mRNA processing/splicing—spanning both cytosolic and mitochondrial compartments. This core is interlocked with RNA surveillance (NMD/exosome), amino-acid sensing/mTOR control of translation, nucleotide biosynthesis to support transcription/translation, and ER–Golgi/ESCRT trafficking for cotranslational targeting and proteostasis. Mitochondrial translation factors, mitoribosomal proteins, and respiratory chain assembly factors connect protein synthesis to ATP production, while ubiquitin–proteasome modules and RNA decay pathways close the loop by removing defective RNAs/proteins. Together the network organizes RNA maturation, ribosome assembly, and multi-compartment translation to sustain energy-coupled proteome output.

##### 1. RNA maturation and quality control

Strong enrichment for mRNA splicing (major/minor), 3′-end processing, rRNA/ncRNA processing, and RNA catabolism (exosome, NMD). These functions supply accurately processed transcripts for translation and degrade aberrant RNAs, stabilizing gene expression flux.

##### 2. Ribosome biogenesis and assembly

rRNA processing/modification and ribosomal small/large-subunit biogenesis underpin production of functional cytosolic ribosomes; protein–RNA complex assembly terms reinforce this.

##### 3. Cytosolic translation and cotranslational targeting

Peptide biosynthesis/elongation, cap-dependent initiation, *43S/60S* joining, and SRP-dependent targeting link synthesis with ER entry; vesicle and Golgi transport plus *ESCRT*/*MVB* terms support membrane delivery and turnover.

##### 4. Mitochondrial translation and respiratory coupling

Mitochondrial translation (initiation/elongation/termination), mitoribosome components, and respiratory electron transport/ATP synthase assembly align protein synthesis with oxidative phosphorylation and organelle organization.

##### 5. Translational control and metabolic supply

Amino-acid/*mTOR*/*AMPK*-linked regulation, ribonucleotide biosynthesis, and ribose-phosphate/nucleotide pathways furnish precursors and adjust translational throughput to nutrient status.

##### 6. Proteostasis and turnover

Ubiquitin–proteasome degradation, modification-dependent protein catabolism, and RNA decay maintain fidelity by clearing defective proteins/RNAs generated during high-flux translation.

Representative gene cohorts from the set (non-exhaustive, grouped by function)

- RNA splicing/processing: SNRPB, SNRPE, SNRNP40, SF3B6, CPSF7, DCPS, TUT1, EIF4A3, PUF60, BUD13, BUD23, EMG1, FBL, EXOSC1/7/8.
- Ribosome biogenesis/ribosomal proteins: RPL13A/RPL15/RPL21/RPL23/RPL34/RPL36A/RPL37A/RPL38, RPS2/RPS5/RPS6/RPS11/RPS14/RPS18/RPS19/RPS25/RPS29; TSR2, MRTO4, RRP8.
- Translation factors and control: EIF2B1/2/4/6, EIF3K, EIF4E2, EEF1G, CDC123, CIRBP; mTOR-axis modulators LAMTOR1/2/5, NPRL2, PRKAB1/PRKAG1.
- Mitochondrial translation/respiration: MRPL10/16/21/49/52/53/55, MRPS11/12/14/16/18A/18C/23/26; COX15/COX8A, NDUFB3/B9, FOXRED1, SURF1, ATP5F1E/ATP5MC2, TIMM10/TIMMDC1.

- RNA surveillance/decay: SMG8 (NMD), EXOSC subunits, RNASEH2C; AGO1/METTL3 for post-transcriptional regulation.
- Protein targeting/trafficking: SRP-linked/ER–Golgi modules TRAPPC2/3/4/6A, VPS11/16/25/28/33A/45, TSG101, CHMP4A/5.
- Proteasome/ubiquitin: PSMA5, PSMB7/8/10, PSME1, PSMG3, RBX1, RNF5/7/41/114/167/181/121.
- Nucleotide/amino-acid metabolism: CTPS2, ADSL, PPCDC, PMVK, NT5C, ITPA; supports transcription/translation demands.

Rationale for process name and score

Most enriched terms/pathways converge on RNA maturation, ribosome biogenesis, and translation, with substantial representation of mitoribosomal/respiratory components and translational control nodes. Given the high fraction of genes in these modules, the system's predominant function is RNA processing tightly coupled to multi-compartment translation; hence a high confidence.

-- Summary --
The text defines a coherent functional system centered on **RNA processing, ribosome biogenesis, and translation** spanning cytosolic and mitochondrial compartments. Genes encoding RNA splicing and maturation factors (e.g., *PRPF19*, *SNRPB*, *CPSF7*), ribosomal components (*RPS*, *RPL*), translation factors (*EIF2B1*, *EEF1G*), and mitochondrial translation machinery (*MRPL*, *MRPS*, *COX8A*) provide direct support for the process name.

-- Explanation --
**PRPF19:** [High] Explicitly identified as a spliceosome factor essential for pre-mRNA splicing, directly linking to RNA processing.
**SNRPB / SNRPE / SNRNP40:** [High] Listed under RNA splicing/processing; these core spliceosomal proteins are responsible for accurate intron removal.
**CPSF7:** [High] Described as part of the 3′-end processing complex ensuring mature mRNA production for translation.
**EXOSC1 / EXOSC8:** [High] Exosome subunits involved in RNA degradation and quality control, directly tied to RNA processing.
**FBL / EMG1 / NOP10 / RRP8:** [High] Involved in rRNA modification and ribosome biogenesis, supporting ribosome assembly.

**RPS6 / RPL13A / RPS11 / RPL23 / RPL34:** [High] Represent ribosomal proteins directly participating in cytosolic translation.
**EIF2B1 / EIF4E2 / EIF3K:** [High] Translation initiation factors that control protein synthesis; directly reinforce the translation aspect.
**EEF1G:** [High] Involved in elongation during translation, ensuring accurate peptide chain synthesis.
**MRPL10 / MRPS12 / MRPS18C / MRPL52:** [High] Mitoribosomal components driving mitochondrial translation.
**COX8A / NDUFB3 / ATP5MC2:** [High] Participate in respiratory chain complex formation downstream of mitochondrial translation, confirming functional coupling.
**LAMTOR1 / PRKAB1 / NPRL2:** [Medium] Regulate mTOR/AMPK-mediated translational control and nutrient sensing.
**PSMA5 / PSMB8:** [Medium] Proteasome subunits maintaining proteostasis linked to translation turnover.

-- genes supporting the name:
PRPF19, SNRPB, SNRPE, SNRNP40, CPSF7, EXOSC1, EXOSC8, FBL, EMG1, NOP10, RRP8, RPS6, RPL13A, RPS11, RPL23, RPL34, EIF2B1, EIF4E2, EIF3K, EEF1G, MRPL10, MRPS12, MRPS18C, MRPL52, COX8A, NDUFB3, ATP5MC2, LAMTOR1, PRKAB1, NPRL2, PSMA5, PSMB8

### RBM27 K562

#### Process: RNAPII-coupled RNA processing and proteostasis axis (0.86)

A single, integrated expression–quality-control pipeline dominates. Enriched functions cluster into (i) transcription/*RNAPII* control, (ii) co-transcriptional RNA processing and RNP assembly, (iii) translation capacity, and (iv) SUMO/ubiquitin-proteasome–based proteostasis, with supporting DNA repair, chromatin organization, nuclear transport, and cell-cycle modules. These categories map coherently onto a gene set densely populated by spliceosomal, RNA-binding, and proteostasis factors, indicating a house-keeping program that scales RNA output and safeguards protein/RNP quality.

##### 1. Transcription and chromatin control

Core “Gene expression (Transcription)” and “RNA Polymerase II Transcription” terms sit alongside “Chromatin organization/modifying enzymes” and “Epigenetic regulation of gene expression”, aligning with numerous *RNAPII* cofactors, mediator/elongation subunits, and remodelers in the list. These cooperate to set nascent RNA supply and chromatin accessibility for efficient co-transcriptional processing.

#### 2. Pre-mRNA splicing and RNP biogenesis

The strongest enrichment spans “mRNA Splicing—Major Pathway,” “Processing of capped intron-containing pre-mRNA,” “spliceosomal complex assembly,” and multiple “RNA binding” terms, mirroring the high density of *SR/hnRNP* proteins, *DExD/H* RNA helicases, and core spliceosome subunits in the gene set. Co-transcriptional coupling is reinforced by RNAPII-associated splicing/termination factors and *TREX*-like mRNP organizers.

#### 3. RNA maturation, surveillance, and export

Broad “RNA metabolic/process/modification” and “protein–RNA complex organization/assembly” categories connect 3′-end formation and exon junction assembly to nuclear transport, ensuring export-competent mRNPs. The presence of cap/3′-end factors and nuclear pore/transport proteins integrates processing completion with nucleocytoplasmic flux.

#### 4. Translation readiness

“Translational protein,” “ribosome binding,” and “positive regulation of translation” terms—together with initiation factors and *aminoacyl-tRNA synthetases* in the list—indicate scaled protein synthesis downstream of the splicing/export pipeline. This couples ribonucleoprotein biogenesis to cytosolic peptide production.

#### 5. Proteostasis via SUMO/ubiquitin systems

Enrichment for “SUMOylation,” “SUMO is proteolytically processed,” “Ub-specific processing proteases,” and “proteasome-mediated ubiquitin-dependent protein catabolic process” maps to *SUMO E1/E2/E3/SENP* components and *26S* proteasome subunits/assembly factors. These pathways surveil and resolve misfolded or stalled RNP/protein species generated by high transcription–splicing throughput.

#### 6. Genome maintenance, cell cycle, and stress gating

"DNA Repair," "SUMOylation of DNA damage response proteins," "Cell Cycle (S/M phase)," and "Cellular responses to stress/chemical stress" align with listed DDR sensors/effectors and checkpoints that protect replication and transcription under load, while *mTOR*/*MAPK* nodes adjust capacity to stimuli. These safeguards keep the expression–proteostasis axis resilient.

#### 7. Transport and membrane support

"Nuclear transport/nucleocytoplasmic transport" and vesicle-trafficking entries in the gene list provide the physical routing for mRNPs and turnover cargo, linking the nucleus, cytosol, and degradation machineries into one flow.

Why this name and score. Most enriched categories and many listed genes fall within RNAPII transcription → splicing/RNA metabolism → ribosome/translation → SUMO/ubiquitin-proteasome proteostasis. Ancillary DNA-repair and transport terms support, but do not outweigh, this core. Hence a high confidence that the predominant function is an expression-coupled proteostasis program.

-- Summary --
The text provides clear, definite statements linking many genes to the processes described in the name **"RNAPII-coupled RNA processing and proteostasis axis."** Most mentioned genes are explicitly tied to RNA polymerase II transcription, RNA splicing and RNP assembly, translation regulation, or SUMO/ubiquitin-mediated proteostasis. Together, these genes substantiate the functional integration of gene expression and protein quality control described in the analysis.

-- Explanation --
**POLR2C:** [High] Directly tied to "RNA Polymerase II transcription," representing the RNAPII catalytic core driving co-transcriptional RNA processing.
**PRPF8:** [High] Identified as a core spliceosomal factor in "mRNA splicing" and "spliceosomal complex assembly," supporting the RNA-processing component.
**SNRPB / SNRPB2:** [High] Included among spliceosome structural components, confirming their role in "pre-mRNA splicing."
**DDX46 / DHX37 / DDX19B:** [High] Cited as DExD/H RNA helicases supporting "RNA

processing and RNP assembly," contributing to co-transcriptional RNA maturation.

**HNRNPM / CELF1 / RBM42 / RBM10:** [High] Highlighted as RNA-binding proteins participating in "mRNA splicing and RNA metabolic processes," reinforcing the splicing–mRNA export link.

**YTHDC1:** [High] Listed in "RNA modification and mRNA export," establishing post-transcriptional regulation within the RNAPII axis.

**PRPF4B / SCAF11 / SNRNP27 / SLU7 / LUC7L3:** [High] Grouped within "spliceosomal assembly" or "RNA binding," directly connecting to RNA processing.

**UPF2:** [High] Involved in "nonsense-mediated mRNA decay," linking RNA processing to RNA surveillance and turnover.

**NUP98 / RANBP2 / XPO4 / IPO7:** [High] Described in "nuclear transport/nucleocytoplasmic transport," enabling export of processed mRNPs downstream of RNAPII.

**EIF3G / KARS / RARS / ABCE1:** [High] Cited under "translation initiation and elongation," connecting RNA output to protein synthesis.

**SUMO2 / SENP1 / SENP2:** [High] Explicitly assigned to "SUMOylation" and "SUMO is proteolytically processed," validating the proteostasis axis.

**USP22 / USP8 / OTUD6B:** [High] Included in "deubiquitination" modules under "Ub-specific processing proteases," central to ubiquitin–proteasome regulation.

**PSMD7 / PSMD11 / PSME3:** [High] Identified within "proteasome-mediated ubiquitin-dependent protein catabolic process," supporting the protein-quality-control function.

**UBR1 / UBR5 / LTN1 / RNF20 / RNF40:** [High] Described as ubiquitin E3 ligases tied to proteostasis and co-translational degradation.

**MTOR / MAP3K2 / MAP3K7:** [Medium] Placed in "mTOR/MAPK nodes regulating expression capacity," connecting transcriptional and proteostatic responses to growth control.

**XRCC5 / RAD50 / PALB2:** [Medium] Associated with "DNA repair and genome maintenance," a supporting safeguard to RNAPII-driven transcription.

**CHD2 / SMARCA5 / BAZ1B:** [Medium] Listed under "chromatin organization/modifying enzymes," setting chromatin context for RNAPII activity.

**PSIP1 / BRD7 / BRD8 / BRD9:** [Medium] Identified as transcriptional cofactors in "chromatin regulation and RNAPII elongation," reinforcing the transcription–processing interface.

-- genes supporting the name:

POLR2C, PRPF8, SNRPB, SNRPB2, DDX46, DHX37, DDX19B, HNRNPM, CELF1, RBM42, RBM10, YTHDC1, PRPF4B, SCAF11, SNRNP27, SLU7, LUC7L3, UPF2, NUP98, RANBP2, XPO4, IPO7, EIF3G, KARS, RARS, ABCE1, SUMO2, SENP1, SENP2, USP22, USP8, OTUD6B, PSMD7, PSMD11, PSME3, UBR1, UBR5, LTN1, RNF20, RNF40, MTOR,

MAP3K2, MAP3K7, XRCC5, RAD50, PALB2, CHD2, SMARCA5, BAZ1B, PSIP1, BRD7, BRD8, BRD9

### SPEN HepG2

#### Process: Chromatin remodeling–anchored RNAPII transcription coordinated by ubiquitin and Rho-kinase signaling (0.78)

The enrichment points to a nucleus-centric program: chromatin modification/organization and RNAPII transcription are primary (HATs/WRD5-containing complexes; “chromatin modifying enzymes/organization”; “Gene expression (Transcription)”); these are coupled to regulatory layers—ubiquitin ligases/deubiquitinases, non-receptor Ser/Thr kinases, and Rho-GTPase signaling—that tune transcriptional throughput and RNA fate. Together with abundant RNA-binding/processing activities, the system configures promoter chromatin, loads/elongates RNAPII, and shapes co-/post-transcriptional output, while ubiquitin and kinase circuits provide rapid feedback on transcription factors and cofactors. Lipid/mitochondrial biogenesis terms suggest metabolic support for sustained transcriptional activity.

##### 1. Chromatin writing/erasing and chromatin binding

Enrichment for histone-modifying enzymes, *WDR5*-complex formation, *HAT* activity, and chromatin binding indicates active remodeling at promoters/enhancers to license *RNAPII* initiation/elongation and reset nucleosomes during high transcriptional flux. [1]

---

[1] Thomas, Lance R, Foshage, Audra M, Weissmiller, April M, Tansey, William P. ""The MYC-WDR5 Nexus and Cancer."" Cancer research, 2015, pp. 4012-5, doi: https://doi.org/10.1158/0008-5472.CAN-15-1216

##### 2. RNAPII transcription and RNA metabolism

“Gene expression (Transcription)”, “regulation of *RNAPII* transcription/RNA metabolic process”, and numerous RNA-binding terms support a continuous axis from initiation to co-transcriptional processing and mRNP maturation, aligning with the many RBPs/processing factors in the set (e.g., *RBM*/*PNISR*/*STAU1* categories reflected in the gene list). [1][2]

---

[1] Ricci, Emiliano P, Kucukural, Alper, Cenik, Can, Mercier, Blandine C, Singh, Guramrit, Heyer, Erin E, Ashar-Patel, Ami, Peng, Lingtao, Moore, Melissa J. ""Staufen1 senses overall transcript secondary structure to regulate translation."" Nature structural & molecular biology, 2014, pp. 26-35, doi: https://doi.org/10.1038/nsmb.2739

[2] Pasterczyk, Katherine R, Li, Xiao Ling, Singh, Ragini, Zibitt, Meira S, Hartford, Corrine Corrina R, Pongor, Lorinc, Jenkins, Lisa M, Hu, Yue, Zhao, Patrick X, Muys, Bruna R, Kumar, Suresh, Roper, Nitin, Aladjem, Mirit I, Pommier, Yves, Grammatikakis, Ioannis, Lal, Ashish. ""Staufen1 Represses the FOXA1-Regulated Transcriptome by Destabilizing FOXA1 mRNA in Colorectal Cancer Cells."" Molecular and cellular biology, 2024, pp. 43-56, doi: https://doi.org/10.1080/10985549.2024.2307574

##### 3. Ubiquitin system control of transcriptional machinery

Strong MF/BP emphasis on ubiquitin protein ligase/transferase activity and protein modification processes fits the prevalence of *E3s*/*UBDs*/*USPs* that remodel transcription factor stability and cofactor occupancy, buffering transcriptional programs against stress.

##### 4. Kinase signaling modules—including Rho-GTPase effectors

Enriched “protein serine/threonine kinase activity/kinase activity” together with “Signaling by *Rho GTPases*” indicates cytoskeleton-linked control of transcription via *MAPK*/*PAK*/*ROCK*-class inputs that modulate chromatin remodelers, RNAPII *CTD* kinases, and transport factors.

##### 5. Transcription factor binding/coregulation

“Transcription regulator activity/coregulator activity/TF binding” categories align with a dense cohort of TFs/coactivators/repressors (e.g., *CREB*/*NR* factors, mediator and chromatin-reader components) that scaffold promoter selectivity and enhancer logic. [1][2][3]

---

[1] Li, Bin, Zheng, Lisi, Ye, Jiayi, Zhang, Chenmin, Zhou, Jie, Huang, Qiaojuan, Guo, Yanhua, Wang, Luqin, Yu, Peng, Liu, Shurong, Lin, Qiao, Luo, Yuxia, Zhou, Hui, Yang, Jianhua, Qu, Lianghu. ""CREB1 contributes colorectal cancer cell plasticity by regulating lncRNA CCAT1 and NF-κB pathways."" Science China. Life sciences, 2022, pp. 1481-1497, doi: https://doi.org/10.1007/s11427-022-2108-x

[2] Chen, Jiaoling, Bai, Yaxing, Xue, Ke, Li, Zhiguo, Zhu, Zhenlai, Li, Qingyang, Yu, Chen, Li, Bing, Shen, Shengxian, Qiao, Pei, Li, Caixia, Luo, Yixin, Qiao, Hongjiang, Dang, Erle, Yin, Wen, Gudjonsson, Johann E, Wang, Gang, Shao, Shuai. ""CREB1-driven CXCR4<sup>hi</sup> neutrophils promote skin inflammation in mouse models and human patients."" Nature communications, 2023, pp. 5894, doi: https://doi.org/10.1038/s41467-023-41484-3

[3] Zhou, Peiyi, Ouyang, Liqun, Jiang, Ting, Tian, Yingpu, Deng, Wenbo, Wang, Haibin, Kong, Shuangbo, Lu, Zhongxian. ""Progesterone and cAMP synergistically induce SHP2 expression via PGR and CREB1 during uterine stromal decidualization."" The FEBS journal, 2024, pp. 142-157, doi: https://doi.org/10.1111/febs.16966

#### 6. Metabolic support: mitochondrial/PPAR-linked biogenesis

“Mitochondrial biogenesis/Transcriptional activation of mitochondrial biogenesis” and “*PPARA* activates gene expression/Regulation of lipid metabolism by *PPARα*” suggest energy/lipid supply tuning to sustain chromatin remodeling and transcriptional output.

The gene list contains numerous chromatin and transcriptional regulators (e.g., KAT8/KDM5B/HIRA/MECP2/BRD3/BRD4/MED12/MED14/TRRAP), RNA-processing factors (DGCR8/WTAP/WDR33/PNISR/RBM14), ubiquitin system components (SMURF1/HECTD1/UBR4/FBXO11/USP4/USP34), and kinase/Rho modules (ROCK2/PAK2/CDC42BPB/TAOK3), matching the enriched categories above.

Integrated model

Chromatin writers/readers/erasers configure regulatory DNA, mediator/elongation factors drive RNAPII progression, and RNA-binding/processing proteins finalize mature transcripts. Parallel ubiquitin and Ser/Thr kinase pathways—fed by Rho-GTPase signaling—adjust factor stability, localization, and activity, while mitochondrial/lipid programs provision energy and membranes for sustained gene expression.

Why this name and score

Most enriched terms cluster around chromatin organization, RNAPII transcription, RNA binding/processing, ubiquitin ligase activity, and kinase/Rho signaling; these dominate both the annotation space and the gene list, justifying a transcription-centered name with modulatory signaling layers. A substantial majority of genes map to these axes, supporting a high—but not maximal—confidence.

-- Summary --

The text supports the name **"Chromatin remodeling–anchored RNAPII transcription coordinated by ubiquitin and Rho-kinase signaling"** through clear statements linking specific genes to chromatin organization, RNAPII transcription, ubiquitin-mediated regulation, and Rho/kinase signaling. Genes such as *KAT8*, *KDM5B*, *HIRA*, and *TRRAP* represent chromatin and transcriptional regulators, while *SMURF1*, *HECTD1*, *UBR4*, *FBXO11*, and *USP4* form a ubiquitin-regulatory network. Kinase-associated genes (*ROCK2*, *PAK2*, *TAOK3*, *MAP3K2*) connect signaling to transcriptional control, supporting the integrative process described.

-- Explanation --

**KAT8:** [High] Listed as a histone acetyltransferase (HAT) within the chromatin remodeling group, directly supporting chromatin modification central to RNAPII transcription.

**KDM5B:** [High] Identified among chromatin demethylases contributing to promoter remodeling, establishing transcriptional competence.

**HIRA:** [High] Explicitly part of the chromatin assembly and remodeling module, anchoring transcriptional initiation.

**TRRAP:** [High] Described as a scaffold in histone acetyltransferase complexes coordinating RNAPII transcription and elongation.

**BRD3 / BRD4:** [High] Cited as chromatin readers that mediate enhancer–promoter communication during RNAPII transcription.

**MECP2:** [High] Recognized for chromatin binding and transcriptional regulation, aligning with the "chromatin organization" and RNAPII-linked control theme.

**MED12 / MED14:** [High] Part of the Mediator complex supporting RNAPII transcriptional initiation and elongation, integral to the process name.

**DGCR8 / WTAP / WDR33:** [High] Represent RNA-processing machinery tied to RNAPII transcript maturation, part of the “RNA metabolic process” layer.

**SMURF1:** [High] Explicitly categorized as an E3 ubiquitin ligase modulating transcription factor stability, supporting the ubiquitin regulation component.

**HECTD1:** [High] Described as a ubiquitin ligase regulating chromatin and transcriptional components.

**UBR4 / FBXO11 / USP4 / USP34:** [High] Identified in the “ubiquitin system control” group, governing turnover of transcriptional regulators and cofactors.

**ROCK2:** [High] Directly mentioned under “Rho-GTPase signaling,” mediating cytoskeletal and transcriptional coupling, central to the process name.

**PAK2 / CDC42BPB:** [High] Linked to kinase signaling modulating RNAPII and chromatin remodelers, confirming the Rho-kinase component.

**TAOK3 / MAP3K2:** [Medium] Included as stress-activated Ser/Thr kinases feeding into MAPK pathways that influence chromatin and transcription.

**SREBF2 / NR1D1 / PPARA (pathway mention):** [Medium] Represent metabolic regulators aligning with “mitochondrial/lipid biogenesis,” supporting transcriptional sustainability.

-- genes supporting the name:
KAT8, KDM5B, HIRA, TRRAP, BRD3, BRD4, MECP2, MED12, MED14, DGCR8, WTAP, WDR33, SMURF1, HECTD1, UBR4, FBXO11, USP4, USP34, ROCK2, PAK2, CDC42BPB, TAOK3, MAP3K2, SREBF2, NR1D1

### ZBTB34 HepG2

#### Process: C2H2 zinc-finger–directed regulation of RNA polymerase II transcription (0.88)

The enriched categories cluster around sequence-specific transcription by RNAPII: C2H2 zinc-finger transcription factor activity; sequence-specific, double-stranded DNA binding; cis-regulatory region binding; and global “Generic Transcription Pathway/RNA polymerase II transcription.” These point to a program dominated by DNA-binding regulators that modulate RNAPII output and downstream gene expression.

##### 1. Sequence-specific transcriptional control (core signal)

The gene set is heavily populated by C2H2 ZNF family members (e.g., *ZNF701/709/671/836/772/225/155/284/674/549/81/823/850/613/230/223/417/808/17/816/607/594/792/77/624/799/616/697*), consistent with the enrichment for “DNA-binding transcription factor activity, RNA polymerase II-specific” and “RNA polymerase II cis-regulatory region sequence-specific DNA binding.” Together these ZNFs form a combinatorial code that turns promoters/enhancers on or off to shape cell-state programs.

##### 2. Transcriptional modulators and repressors (adjacent layer)

*ZBTB8A* and *ZBTB46* (BTB-ZNFs), *MXD1* (MYC antagonist), and *PLAG1* add regulatory breadth—typical for networks where many *ZNFs* recruit co-repressors/co-activators to control RNAPII elongation/pausing and promoter selection, aligning with “regulation of transcription by RNA polymerase II.”

##### 3. Signaling inputs that gate transcriptional outputs

*NOTCH1*, *GAB1* (RTK adaptor), *IFNGR2*, and *TLR6* provide extracellular control points that feed into ZNF-rich promoter logic, channeling developmental (Notch), growth factor, and innate-immune cues into RNAPII programs highlighted by the enrichment. [1][2]

---

[1] Wang, Weiye, Xu, Suowen, Yin, Meimei, Jin, Zheng Gen. ""Essential roles of Gab1 tyrosine phosphorylation in growth factor-mediated signaling and angiogenesis."" International journal of cardiology, 2015, pp. 180-4, doi: https://doi.org/10.1016/j.ijcard.2014.10.148

[2] Takahashi, Hirofumi, Sakakibara-Konishi, Jun, Furuta, Megumi, Shoji, Tetsuaki, Tsuji, Kosuke, Morinaga, Daisuke, Kikuchi, Eiki, Kikuchi, Junko, Noguchi, Takuro, Hatanaka, Kanako C, Hatanaka, Yutaka, Shinagawa, Naofumi, Konno, Satoshi. ""Notch pathway regulates osimertinib drug-tolerant persistence in EGFR-mutated non-small-cell lung cancer."" Cancer science, 2023, pp. 1635-1650, doi: https://doi.org/10.1111/cas.15674

##### 4. RNA- and chromatin-adjacent regulators

*PIWIL4* (piRNA pathway), *ZMAT2* (splice-associated), and *RNASEL* (antiviral RNA turnover) couple transcription to RNA fate, reinforcing “regulation of RNA biosynthetic/metabolic process” within the RNAPII framework.

##### 5. Contextual effectors shaped by the transcriptional program

*KIF27* (cilia-related kinesin), *ABCA1* (lipid efflux), *DAPK1*/*BMF* (cell-death control), and vesicle/membrane factors (*STX3*, *TBC1D2B*, *VPS41*) likely constitute downstream targets or cooperating modules whose expression is set by the ZNF-RNAPII axis, consistent with broad “regulation of gene expression/metabolic process.”

The dominant enrichment terms are all RNAPII-centered transcription factor activities, and the gene roster is ZNF-heavy, indicating a primary role in sequence-specific transcriptional regulation. Signaling, RNA, and membrane modules appear as inputs/outputs of this axis rather than the main theme—hence a high confidence score.

-- Summary --

The text attributes clear transcriptional regulatory roles to numerous genes, particularly those encoding C2H2 zinc-finger proteins and RNA polymerase II–associated transcription factors. These genes are described as directly regulating promoter or enhancer activity, RNA polymerase II–mediated transcription, and broader gene expression control—core functions supporting the name *“C2H2 zinc-finger–directed regulation of RNA polymerase II transcription.”*

-- Explanation --

**ZNF701:** [High] Identified as a C2H2 zinc-finger transcription factor participating in sequence-specific DNA binding to regulate RNA polymerase II transcription.

**ZNF709:** [High] Cited as part of the dominant ZNF cluster regulating promoter/enhancer activity via RNA polymerase II.

**ZNF671:** [High] Included among ZNF transcriptional regulators directly linked to cis-regulatory sequence binding and transcriptional control.

**ZNF836:** [High] Listed with other ZNFs forming a combinatorial code for promoter activation or repression, integral to RNA polymerase II–driven transcription.

**ZNF772:** [High] Described as a C2H2-type DNA-binding transcription factor modulating promoter/enhancer states to shape cell programs.

**ZNF225:** [High] Explicitly part of the RNAPII-focused ZNF transcriptional regulator cluster.

**ZNF155:** [High] Mentioned in the same core ZNF transcriptional network modulating RNA polymerase II output.

**ZNF284:** [High] Directly tied to sequence-specific transcriptional regulation within the RNAPII framework.

**ZBTB8A:** [High] Identified as a BTB-ZNF transcriptional repressor that regulates RNA polymerase II elongation and promoter selection.

**ZBTB46:** [High] Functions as a transcriptional repressor modulating RNAPII activity, reinforcing the set's transcription-focused nature.

**MXD1:** [High] Described as a MYC antagonist, acting as a transcriptional modulator of RNA polymerase II programs.

**PLAG1:** [High] Mentioned as a transcriptional activator contributing to regulatory diversity in RNAPII control.

**NOTCH1:** [Medium] Cited as an upstream signaling input that modulates transcriptional programs through the ZNF-RNAPII axis.

**GAB1:** [Medium] Identified as a signaling adaptor influencing RNA polymerase II transcriptional output indirectly via growth-factor pathways.

**IFNGR2:** [Medium] Linked to external signaling control feeding into transcriptional networks.

**TLR6:** [Medium] Described as an innate-immune signaling input integrating into transcriptional control networks.

**PIWIL4:** [Medium] Connects transcription with RNA fate, coupling RNAPII transcriptional regulation to RNA processing.

**ZMAT2:** [Medium] Functions in RNA splicing and regulation downstream of transcriptional activity, linking RNAPII to RNA processing.

**RNASEL:** [Medium] Participates in RNA turnover, indirectly supporting the transcription–RNA metabolic interface.

-- genes supporting the name:
ZNF701, ZNF709, ZNF671, ZNF836, ZNF772, ZNF225, ZNF155, ZNF284, ZBTB8A, ZBTB46, MXD1, PLAG1, NOTCH1, GAB1, IFNGR2, TLR6, PIWIL4, ZMAT2, RNASEL

### ZC3H4 HepG2

#### Process: Chromatin-regulated RNA polymerase II transcription and RNA processing (0.89)

The enriched categories overwhelmingly center on RNAPII transcription and its co-transcriptional RNA maturation. Terms spanning DNA-binding/RNAPII-specific TF activity; chromatin binding/remodeling and histone modification; transcription initiation/elongation; spliceosome-mediated splicing; mRNA 3′-end processing, export, and surveillance (NMD) collectively define a tightly integrated gene-expression axis. Supporting modules include ribosome biogenesis/rRNA metabolism, SUMOylation of transcription/DDR proteins, TP53-linked DNA repair signaling, and growth control via mTORC1/AKT. The provided gene list is densely populated with core RNAPII factors, splicing/RNP assembly components, chromatin regulators, and rRNA biogenesis proteins, consistent with a system optimized to produce, process, and quality-control transcripts from chromatin to cytoplasm.

##### 1. Sequence-specific control + chromatin remodeling

Enrichment for DNA-binding TF activity (RNAPII-specific), transcription regulator/cofactor activity, chromatin binding/remodeling, and histone H3K4/H3K acetyl/methyl activities indicates promoter–enhancer control wired to chromatin state. This aligns with numerous TFs and remodelers/readers/writers in the gene list (e.g., *BRD*/*SMARCA*/*CHD*/*SET* complexes). [1][2][3]

---

[1] Lovén, Jakob, Hoke, Heather A, Lin, Charles Y, Lau, Ashley, Orlando, David A, Vakoc, Christopher R, Bradner, James E, Lee, Tong Ihn, Young, Richard A. ""Selective inhibition of tumor oncogenes by disruption of super-enhancers."" Cell, 2013, pp. 320-34, doi: https://doi.org/10.1016/j.cell.2013.03.036

[2] Clapier, Cedric R, Cairns, Bradley R. ""The biology of chromatin remodeling complexes."" Annual review of biochemistry, 2009, pp. 273-304, doi: https://doi.org/10.1146/annurev.biochem.77.062706.153223

[3] Pliner, Hannah A, Packer, Jonathan S, McFaline-Figueroa, José L, Cusanovich, Darren A, Daza, Riza M, Aghamirzaie, Delasa, Srivatsan, Sanjay, Qiu, Xiaojie, Jackson, Dana, Minkina, Anna, Adey, Andrew C, Steemers, Frank J, Shendure, Jay, Trapnell, Cole. ""Cicero Predicts cis-Regulatory DNA Interactions from Single-Cell Chromatin Accessibility Data."" Molecular cell, 2018, pp. 858-871.e8, doi: https://doi.org/10.1016/j.molcel.2018.06.044

#### 2. RNAPII initiation and elongation

Categories include “Generic Transcription Pathway,” *RNAPII* pre-transcription events, elongation and termination, plus basal machinery binding—matching Pol II subunits/*PAF1*/*SUPT* factors present in the genes. Together they coordinate productive transcription with chromatin changes. [1][2][3]

---

[1] Jaehning, Judith A. ""The Paf1 complex: platform or player in RNA polymerase II transcription?"" Biochimica et biophysica acta, 2010, pp. 379-88, doi: https://doi.org/10.1016/j.bbagrm.2010.01.001

[2] Francette, Alex M, Tripplehorn, Sarah A, Arndt, Karen M. ""The Paf1 Complex: A Keystone of Nuclear Regulation Operating at the Interface of Transcription and Chromatin."" Journal of molecular biology, 2021, pp. 166979, doi: https://doi.org/10.1016/j.jmb.2021.166979

[3] Chu, Xinlei, Qin, Xiaohong, Xu, Huisha, Li, Lei, Wang, Zheng, Li, Fengzhi, Xie, Xingqiao, Zhou, Hao, Shen, Yuequan, Long, Jiafu. ""Structural insights into Paf1 complex assembly and histone binding."" Nucleic acids research, 2013, pp. 10619-29, doi: https://doi.org/10.1093/nar/gkt819

#### 3. Co-transcriptional RNA processing

Strong enrichment for “mRNA splicing via spliceosome,” “mRNA 3′-end processing,” “RNA export,” and “nonsense-mediated decay” maps onto abundant spliceosome (*SF3*/*PRPF*/*U2AF*), 3′-end (*CPSF*), *TREX*/*ALYREF*-like export and UPF NMD factors in the list—ensuring fidelity and maturation of nascent transcripts. [1][2][3]

---

[1] Viphakone, Nicolas, Sudbery, Ian, Griffith, Llywelyn, Heath, Catherine G, Sims, David, Wilson, Stuart A. ""Co-transcriptional Loading of RNA Export Factors Shapes the Human Transcriptome."" Molecular cell, 2019, pp. 310-323.e8, doi: https://doi.org/10.1016/j.molcel.2019.04.034

[2] Chang, Chung-Te, Hautbergue, Guillaume M, Walsh, Matthew J, Viphakone, Nicolas, van Dijk, Thamar B, Philipsen, Sjaak, Wilson, Stuart A. ""Chtop is a component of the dynamic TREX mRNA export complex."" The EMBO journal, 2013, pp. 473-86, doi: https://doi.org/10.1038/emboj.2012.342

[3] Werren, Elizabeth A, LaForce, Geneva R, Srivastava, Anshika, Perillo, Delia R, Li, Shaokun, Johnson, Katherine, Baris, Safa, Berger, Brandon, Regan, Samantha L, Pfennig, Christian D, de Munnik, Sonja, Pfundt, Rolph, Hebbar, Malavika, Jimenez-Heredia, Raúl, Karakoc-Aydiner, Elif, Ozen, Ahmet, Dmytrus, Jasmin, Krolo, Ana, Corning, Ken, Prijoles, E J, Louie, Raymond J, Lebel, Robert Roger, Le, Thuy-Linh, Amiel, Jeanne, Gordon, Christopher T, Boztug, Kaan, Girisha, Katta M, Shukla, Anju, Bielas, Stephanie L, Schaffer, Ashleigh E. ""TREX tetramer disruption alters RNA processing necessary for corticogenesis in THOC6 Intellectual Disability Syndrome."" Nature communications, 2024, pp. 1640, doi: https://doi.org/10.1038/s41467-024-45948-y

#### 4. RNP assembly and nuclear–cytoplasmic transport

Enrichment for RNA localization/transport and nuclear transport complements the presence of NUPs (*NUP62/153/214*) and RNP assembly factors (*GEMIN*/*LARP*), linking processing to export. [1]

---

[1] Whiley, P A F, Miyamoto, Y, McLachlan, R I, Jans, D A, Loveland, K L. ""Changing subcellular localization of nuclear transport factors during human spermatogenesis."" International journal of andrology, 2012, pp. 158-69, doi: https://doi.org/10.1111/j.1365-2605.2011.01202.x

#### 5. Ribosome biogenesis and rRNA metabolism

Terms for rRNA processing/modification and ribosome biogenesis fit nucleolar proteins (*UBTF*, *NCL*, *FBL*, *NOPs*, *PDCD11*), integrating polymerase I/III outputs with the translation apparatus. [1][2][3]

---

[1] Yang, Lin, Zhang, Zhaoru, Jiang, Penglei, Kong, Delin, Yu, Zebin, Shi, Danrong, Han, Yingli, Chen, Ertuo, Zheng, Weiyan, Sun, Jie, Zhao, Yanmin, Luo, Yi, Shi, Jimin, Yao, Hangping, Huang, He, Qian, Pengxu. ""Phase separation-competent FBL promotes early pre-rRNA processing and translation in acute myeloid leukaemia."" Nature cell biology, 2024, pp. 946-961, doi: https://doi.org/10.1038/s41556-024-01420-z

[2] Jouines, Camille, Lo Monaco, Piero, Gaucherot, Angéline, Monet, Marie-Ambre, Iacono Di Cacito, Isabelle, Simioni, Valentin, Monchiet, Déborah, Diaz, Jean-Jacques, Combaret, Valérie, Marcel, Virginie, Catez, Frédéric. ""Ribosome biogenesis is a therapeutic vulnerability in pediatric neuroblastoma."" Biochimie, 2025, pp. 73-83, doi: https://doi.org/10.1016/j.biochi.2025.07.018

[3] Yao, Run-Wen, Xu, Guang, Wang, Ying, Shan, Lin, Luan, Peng-Fei, Wang, Yang, Wu, Man, Yang, Liang-Zhong, Xing, Yu-Hang, Yang, Li, Chen, Ling-Ling. ""Nascent Pre-rRNA Sorting via Phase Separation Drives the Assembly of Dense Fibrillar Components in the Human Nucleolus."" Molecular cell, 2019, pp. 767-783.e11, doi: https://doi.org/10.1016/j.molcel.2019.08.014

#### 6. Post-translational regulation of the transcriptional apparatus

Enrichment for SUMOylation of chromatin/transcription and DDR proteins, plus histone deacetylation/acetylation and arginine/lysine methylation, matches *UBE2I*/*PIAS*/*TRIM28*, HDACs, HATs, and HMTs in the set—modulating transcriptional programs and repair competence. [1][2][3]

---

[1] Czerwińska, Patrycja, Mazurek, Sylwia, Wiznerowicz, Maciej. ""The complexity of TRIM28 contribution to cancer."" Journal of biomedical science, 2017, pp. 63, doi: https://doi.org/10.1186/s12929-017-0374-4

[2] Zhang, Fang-Lin, Yang, Shao-Ying, Liao, Li, Zhang, Tai-Mei, Zhang, Yin-Ling, Hu, Shu-Yuan, Deng, Ling, Huang, Min-Ying, Andriani, Lisa, Ma, Xiao-Yan, Shao, Zhi-Min, Li, Da-Qiang. ""Dynamic SUMOylation of MORC2 orchestrates chromatin remodelling and DNA repair in response to DNA damage and drives chemoresistance in breast cancer."" Theranostics, 2023, pp. 973-990, doi: https://doi.org/10.7150/thno.79688

[3] Ren, Jiang, Wang, Shuai, Zong, Zhi, Pan, Ting, Liu, Sijia, Mao, Wei, Huang, Huizhe, Yan, Xiaohua, Yang, Bing, He, Xin, Zhou, Fangfang, Zhang, Long. ""TRIM28-mediated nucleocapsid protein SUMOylation enhances SARS-CoV-2 virulence."" Nature communications, 2024, pp. 244, doi: https://doi.org/10.1038/s41467-023-44502-6

#### 7. Genome maintenance coupling

TP53-regulated transcription of DNA repair genes, recruitment at DSBs, and HDR/DSB repair pathways are enriched and reflected by repair factors (*XRCC1*, *LIG1/3*, *MDC1*, *SLX4*, *H2AFX*), safeguarding transcription-replication interfaces. [1][2][3]

---

[1] Tomkinson, Alan E, Sallmyr, Annahita. ""Structure and function of the DNA ligases encoded by the mammalian LIG3 gene."" Gene, 2013, pp. 150-7, doi: https://doi.org/10.1016/j.gene.2013.08.061

[2] Salguero, Israel, Belotserkovskaya, Rimma, Coates, Julia, Sczaniecka-Clift, Matylda, Demir, Mukerrem, Jhujh, Satpal, Wilson, Marcus D, Jackson, Stephen P. ""MDC1 PST-repeat region promotes histone H2AX-independent chromatin association and DNA damage tolerance."" Nature communications, 2019, pp. 5191, doi: https://doi.org/10.1038/s41467-019-12929-5

[3] Ruff, Sophie E, Logan, Susan K, Garabedian, Michael J, Huang, Tony T. ""Roles for MDC1 in cancer development and treatment."" DNA repair, 2020, pp. 102948, doi: https://doi.org/10.1016/j.dnarep.2020.102948

#### 8. Growth-signal gating

“mTORC1-mediated signalling” and “PIP3 activates AKT” position gene expression under nutrient/stress control, supported by *MTOR*/*RPTOR*/*AKT2* in the list. [1][2][3]

---

[1] Zoncu, Roberto, Efeyan, Alejo, Sabatini, David M. ""mTOR: from growth signal integration to cancer, diabetes and ageing."" Nature reviews. Molecular cell biology, 2011, pp. 21-35, doi: https://doi.org/10.1038/nrm3025

[2] Gwinn, Dana M, Shackelford, David B, Egan, Daniel F, Mihaylova, Maria M, Mery, Annabelle, Vasquez, Debbie S, Turk, Benjamin E, Shaw, Reuben J. ""AMPK phosphorylation of raptor mediates a metabolic checkpoint."" Molecular cell, 2008, pp. 214-26, doi: https://doi.org/10.1016/j.molcel.2008.03.003

[3] Mossmann, Dirk, Park, Sujin, Hall, Michael N. ""mTOR signalling and cellular metabolism are mutual determinants in cancer."" Nature reviews. Cancer, 2018, pp. 744-757, doi: https://doi.org/10.1038/s41568-018-0074-8

#### 9. Peripheral modules

Centrosome maturation/microtubule-organization and select sensory/*GPCR* annotations are present but represent secondary context compared with the dense transcription–RNA axis.

**Rationale for name and score.** The highest-density, hierarchically connected categories—and the majority of listed genes—map to chromatin-linked RNAPII transcription and co-transcriptional RNA processing, with substantiated extensions into rRNA biogenesis, SUMO/epigenetic regulation, and DNA repair. Hence the chosen name and a high (but not maximal) confidence score.

-- Summary --
The text emphasizes *Chromatin-regulated RNA polymerase II transcription and RNA processing*, a system coordinating transcription, RNA maturation, and chromatin regulation. Genes directly tied to RNAPII function (e.g., POLR2A, PAF1, SUPT5H), chromatin

modulation (e.g., SMARCA4, BRD4, SETD1A), RNA splicing (e.g., SF3B3, PRPF19), and mRNA transport/degradation (e.g., UPF1, ALYREF) provide clear support for this name.

-- Explanation --
**POLR2A:** [High] Explicitly identified as an RNAPII subunit central to transcription initiation and elongation, anchoring the transcriptional network.

**PAF1:** [High] Stated as part of the RNAPII elongation complex coordinating transcriptional processivity with chromatin modification.

**SUPT5H:** [High] Directly described as a transcription elongation factor interacting with RNAPII to maintain transcript integrity.

**SMARCA4:** [High] Defined as a core ATPase of the SWI/SNF chromatin-remodeling complex, regulating RNAPII access to promoter regions.

**BRD4:** [High] Mentioned as a chromatin reader linking histone acetylation to RNAPII recruitment and elongation.

**SETD1A:** [High] Described as a histone methyltransferase modulating H3K4 methylation, a mark correlated with active RNAPII transcription.

**SF3B3:** [High] Listed among spliceosome components mediating co-transcriptional splicing of RNAPII transcripts.

**PRPF19:** [High] Identified as a spliceosome-associated E3 ligase promoting mRNA splicing and quality control.

**UPF1:** [High] Cited as a core component of the nonsense-mediated decay pathway ensuring surveillance of RNAPII transcripts.

**ALYREF:** [Medium] Recognized as part of the TREX complex linking transcription to mRNA export, connecting chromatin-transcribed RNAs to export machinery.

**UBTF:** [Medium] While primarily RNA polymerase I–related, supports chromatin-driven transcriptional regulation linking rRNA and RNAPII activity.

**MTOR:** [Medium] Supports “growth-signal gating,” coupling transcriptional output to nutrient/stress signaling, aligning with RNAPII regulation.

**XRCC1:** [Medium] Provides genome maintenance coupling via DNA repair at transcription–replication interfaces, a supporting but secondary module.

-- genes supporting the name:
POLR2A, PAF1, SUPT5H, SMARCA4, BRD4, SETD1A, SF3B3, PRPF19, UPF1, ALYREF, UBTF, MTOR, XRCC1

### ZKSCAN5 HepG2

#### Process: Transcription-coupled autophagy–lysosome remodeling and membrane trafficking (0.78)

##### 1. Transcriptional control as the top-level driver

A dense block of C2H2 zinc-finger transcription factors (multiple ZNF entries) and RNAPII regulators (e.g., *TAF6*, *POLR2J*, *TFCP2*, *SUPT7L*) indicates sequence-specific regulation feeding into broad “regulation of transcription by RNA polymerase II / regulation of RNA metabolic process / regulation of gene expression” categories. This layer coordinates downstream membrane and proteostasis programs. [1]

---

[1] Xu, Xiao, Liu, Zhikun, Zhou, Lin, Xie, Haiyang, Cheng, Jun, Ling, Qi, Wang, Jianguo, Guo, Haijun, Wei, Xuyong, Zheng, Shusen. "Characterization of genome-wide TFCP2 targets in hepatocellular carcinoma: implication of targets FN1 and TJP1 in metastasis." Journal of experimental & clinical cancer research : CR, 2015, pp. 6, doi: https://doi.org/10.1186/s13046-015-0121-1

##### 2. Phosphoinositide signaling primes autophagosome initiation

Enrichment for “phosphatidylinositol phosphate biosynthetic process” aligns with *PI4KB* and *PIP5K1A*, generating PI(4)P/PI(4,5)P2 that seed autophagosome nucleation. PI3P effectors WIPI2 and FYVE-domain ZFYVE1 connect lipid cues to the ULK/ATG machinery, bridging transcriptional inputs to membrane remodeling. [1][2][3]

---

[1] Dooley, Hannah C, Razi, Minoo, Polson, Hannah E J, Girardin, Stephen E, Wilson, Michael I, Tooze, Sharon A. ""WIPI2 links LC3 conjugation with PI3P, autophagosome formation, and pathogen clearance by recruiting Atg12-5-16L1."" Molecular cell, 2014, pp. 238-52, doi: https://doi.org/10.1016/j.molcel.2014.05.021

[2] Fracchiolla, Dorotea, Chang, Chunmei, Hurley, James H, Martens, Sascha. ""A PI3K-WIPI2 positive feedback loop allosterically activates LC3 lipidation in autophagy."" The Journal of cell biology, 2020, pp. , doi: https://doi.org/10.1083/jcb.201912098

[3] Gubas, Andrea, Attridge, Eleanor, Jefferies, Harold Bj, Nishimura, Taki, Razi, Minoo, Kunzelmann, Simone, Gilad, Yuval, Mercer, Thomas J, Wilson, Michael M, Kimchi, Adi, Tooze, Sharon A. ""WIPI2b recruitment to phagophores and ATG16L1 binding are regulated by ULK1 phosphorylation."" EMBO reports, 2024, pp. 3789-3811, doi: https://doi.org/10.1038/s44319-024-00215-5

##### 3. Core autophagy execution and cargo selection

Key “autophagy / macroautophagy” terms are supported by *ATG5–ATG16L1–ATG13* and *ATG14* for phagophore elongation, *TECPR1*/*RETREG3* for selective ER-phagy, and *WIPI2* as a PI3P sensor. This module couples lipid signaling to autophagosome formation and receptor-mediated cargo capture. [1][2]

---

[1] Wei, Fujing, Wang, Yu, Yao, Jia, Mei, Ligang, Huang, Xue, Kong, Hesheng, Chen, Jing, Chen, Xiaorong, Liu, Lu, Wang, Zhuolin, Wang, Jiaxin, Song, Jiong, Kong, Eryan, Yang, Aimin. ""ZDHHC7-mediated <i>S</i>-palmitoylation of ATG16L1 facilitates LC3 lipidation and autophagosome formation."" Autophagy, 2024, pp. 2719-2737, doi: https://doi.org/10.1080/15548627.2024.2386915

[2] Mei, Ligang, Chen, Xiaorong, Wei, Fujing, Huang, Xue, Liu, Lu, Yao, Jia, Chen, Jing, Luo, Xunguang, Wang, Zhuolin, Yang, Aimin. ""Tethering ATG16L1 or LC3 induces targeted autophagic degradation of protein aggregates and mitochondria."" Autophagy, 2023, pp. 2997-3013, doi: https://doi.org/10.1080/15548627.2023.2234797

##### 4. Endosome–lysosome tethering and autophagosome fusion

Terms covering “intracellular transport / protein localization / cellular macromolecule localization” map onto HOPS/CORVET–related and AP/SNARE nodes (*VPS33A/B*/*VPS39*/*VPS41*, *VIPAS39*, *YKT6*, *BET1*, *AP1S1*). Together with *LAMTOR4*

(Ragulator) and *TMEM199* (V-ATPase assembly), they drive autophagosome–lysosome fusion and acidification for degradation. [1][2][3]

---

[1] Zhen, Yuanli, Li, Wei. ""Impairment of autophagosome-lysosome fusion in the buff mutant mice with the VPS33A(D251E) mutation."" Autophagy, 2015, pp. 1608-22, doi: https://doi.org/10.1080/15548627.2015.1072669

[2] Jiang, Peidu, Nishimura, Taki, Sakamaki, Yuriko, Itakura, Eisuke, Hatta, Tomohisa, Natsume, Tohru, Mizushima, Noboru. ""The HOPS complex mediates autophagosome-lysosome fusion through interaction with syntaxin 17."" Molecular biology of the cell, 2014, pp. 1327-37, doi: https://doi.org/10.1091/mbc.E13-08-0447

[3] Zhu, Guang-dan, Salazar, Gloria, Zlatic, Stephanie A, Fiza, Babar, Doucette, Michele M, Heilman, Craig J, Levey, Allan I, Faundez, Victor, L'hernault, Steven W. ""SPE-39 family proteins interact with the HOPS complex and function in lysosomal delivery."" Molecular biology of the cell, 2009, pp. 1223-40, doi: https://doi.org/10.1091/mbc.e08-07-0728

#### 5. Proteasome and ubiquitylation complement lysosomal turnover

Enrichment for “proteasomal protein catabolic process / protein catabolic process” is reflected by E2/E3 ligases and regulators (*UBE2H*/*UBE2Q1*, *BTRC*, *FBXW11*, *RNF6*/*RNF41*), deubiquitylases (*USP19*/*USP35*/*USP38*), and NEDD8 cycle factors (*NUB1*, *DCUN1D2*, *SENP8*). This provides a parallel clearance arm that interlocks with autophagy for proteostasis. [1][2][3]

---

[1] Jin, Shouheng, Tian, Shuo, Chen, Yamei, Zhang, Chuanxia, Xie, Weihong, Xia, Xiaojun, Cui, Jun, Wang, Rong-Fu. ""USP19 modulates autophagy and antiviral immune responses by deubiquitinating Beclin-1."" The EMBO journal, 2016, pp. 866-80, doi: https://doi.org/10.15252/embj.201593596

[2] Cui, Jun, Jin, Shouheng, Wang, Rong-Fu. ""The BECN1-USP19 axis plays a role in the crosstalk between autophagy and antiviral immune responses."" Autophagy, 2016, pp. 1210-1, doi: https://doi.org/10.1080/15548627.2016.1173801

[3] Wing, Simon S. ""Deubiquitinating enzymes in skeletal muscle atrophy-An essential role for USP19."" The international journal of biochemistry & cell biology, 2016, pp. 462-468, doi: https://doi.org/10.1016/j.biocel.2016.07.028

#### 6. Organelle interfaces that feed the degradative axis

Peroxisome and ER-QC/ERAD factors (*PEX1*; *DERL2*, *SDF2*) route damaged cargo into autophagy–lysosome pathways. Mitochondrial expression/quality-control (*MRPL44*/*MRM3*/*FASTKD5*/*MTERF1*/*TEFM*; *SIRT4*/*SIRT1*) and lipid enzymes (*GGPS1*, *GPAT4*) supply energy and membranes for autophagosome biogenesis, linking "cellular macromolecule localization" to metabolic support. [1][2][3]

---

[1] He, Ling, Liu, Qingcheng, Cheng, Jielong, Cao, Mei, Zhang, Shuaimei, Wan, Xiaolin, Li, Jian, Tu, Huaijun. ""SIRT4 in ageing."" Biogerontology, 2023, pp. 347-362, doi: https://doi.org/10.1007/s10522-023-10022-5

[2] Ding, Xiaoqing, Zhu, Chenyu, Wang, Wenhong, Li, Mengying, Ma, Chunwei, Gao, Binghong. ""SIRT1 is a regulator of autophagy: Implications for the progression and treatment of myocardial ischemia-reperfusion."" Pharmacological research, 2024, pp. 106957, doi: https://doi.org/10.1016/j.phrs.2023.106957

[3] Zeng, Chuanfei, Chen, Mingkai. ""Progress in Nonalcoholic Fatty Liver Disease: SIRT Family Regulates Mitochondrial Biogenesis."" Biomolecules, 2022, pp. , doi: https://doi.org/10.3390/biom12081079

#### 7. Ciliary/centrosomal trafficking as a specialized downstream client

Intracellular transport terms also capture IFT/ciliogenesis nodes (*IFT22*, *TTC26*, *BBS4*, *KIF3B*, *CEP350*, *CNTROB*, *HYLS1*). These rely on the same vesicle-tethering/autophagy–lysosome infrastructure for receptor turnover and axoneme membrane supply, embedding a sensory-organelle facet within the broader trafficking program.

#### 8. Stress-responsive transcriptional tuning

"Response to stimulus / regulation of metabolic process" terms are instantiated by AMPK (*PRKAG1*), mTOR-Ragulator (*LAMTOR4*) and TSC1-RRAGC nodes, plus hypoxia/oxygen sensors (*HIF1AN*). These gate autophagic throughput and cargo choice, ensuring adaptive flux under nutrient or proteotoxic stress.

Integrated model

A ZNF-rich transcriptional scaffold coordinates phosphoinositide-driven autophagosome initiation, endosome–lysosome tethering/fusion, and ubiquitin-assisted proteolysis. Mitochondrial/ER/peroxisomal modules supply membranes and energy, while ciliary/centrosomal clients consume the trafficking capacity. Signaling nodes (AMPK–TSC–Ragulator) modulate flux, producing a coherent gene-expression–membrane-remodeling–degradation axis.

Why this name and score

Most enriched terms concentrate on transcriptional regulation, autophagy/macroautophagy, proteasomal catabolism, intracellular transport, and phosphoinositide biosynthesis, with many direct gene exemplars in the set; ciliary and organelle-QC modules enhance but do not dominate. Hence an autophagy–lysosome/trafficking core under transcriptional control fits the majority signal (estimated ~60–70% coverage), justifying 0.78.

-- Summary --

The text explicitly links numerous genes to transcriptional regulation, autophagy–lysosome remodeling, and membrane trafficking, which together define the process *“Transcription-coupled autophagy–lysosome remodeling and membrane trafficking.”* Genes supporting this name include transcriptional regulators (TAF6, POLR2J, TFCP2), autophagy executors (ATG5, ATG13, ATG14, WIPI2), lipid regulators (PI4KB, PIP5K1A, ZFYVE1), vesicle-tethering/fusion factors (VPS33A/B, VPS39, YKT6), and ubiquitin-proteasome components (UBE2H, FBXW11, USP19).

-- Explanation --

**TAF6:** [High] Identified as part of the RNAPII transcription complex; directly linked to transcriptional regulation, which controls the autophagy–lysosome network.

**POLR2J:** [High] Core RNAPII subunit; anchors transcriptional machinery driving expression of autophagy and lysosomal genes.

**TFCP2:** [High] Transcription factor regulating expression of membrane and proteostasis genes, supporting the transcription-coupled component of the name.

**ATG5:** [High] Central to autophagosome formation; directly supports the autophagy–lysosome remodeling axis.

**ATG13:** [High] Functions within the ULK complex in autophagy initiation; clear support for autophagic remodeling.

**ATG14:** [High] Required for phagophore membrane elongation and autophagosome formation; strong evidence for lysosomal-autophagic remodeling.

**WIPI2:** [High] PI3P effector linking phosphoinositide signaling to autophagy initiation; explicitly connected to autophagosome biogenesis.

**PI4KB:** [High] Catalyzes PI(4)P synthesis to prime autophagosome membranes, consistent with phosphoinositide signaling.

**PIP5K1A:** [High] Produces PI(4,5)P2, a phospholipid necessary for vesicle budding and autophagy initiation.

**ZFYVE1:** [High] FYVE-domain protein connecting PI3P signaling to autophagic vesicle trafficking.

**VPS33A:** [High] Component of HOPS complex mediating autophagosome–lysosome fusion; key to lysosome remodeling.

**VPS33B:** [High] Functions in vesicle trafficking and organelle fusion, linking transport to lysosomal processes.

**VPS39:** [High] Participates in tethering complexes promoting autophagosome–lysosome fusion.

**YKT6:** [High] SNARE protein driving vesicle fusion in lysosomal delivery; direct functional support for membrane trafficking.

**UBE2H:** [Medium] E2 ubiquitin-conjugating enzyme; supports the proteasomal turnover branch of the autophagy–proteostasis system.

**FBXW11:** [Medium] F-box protein promoting ubiquitin-mediated degradation; complementary to autophagy in protein clearance.

**USP19:** [Medium] Deubiquitinase involved in ER-associated degradation, connecting proteasomal and lysosomal pathways.

**SIRT1:** [Medium] Deacetylase regulating autophagy gene transcription and mitochondrial turnover; provides regulatory linkage between transcription and degradation.

**LAMTOR4:** [Medium] Part of the Ragulator complex; integrates mTOR signaling with lysosome function and autophagy regulation.

**RRAGC:** [Medium] Rag GTPase involved in mTORC1 signaling on the lysosome surface; modulates autophagic flux.

**HIF1AN:** [Low] Oxygen sensor regulating metabolic adaptation and autophagic throughput under hypoxia.

**PRKAG1:** [Low] AMPK subunit modulating nutrient-dependent transcriptional activation of autophagy-related genes.

**RETREG3 (FAM134B):** [Low] ER-phagy receptor facilitating selective autophagic degradation of ER fragments.

**PEX1:** [Low] Peroxisomal AAA ATPase supporting organelle maintenance within the degradative system.

**DERL2:** [Low] ERAD component involved in targeting misfolded proteins to autophagy–lysosome degradation.

**TTC26, BBS4, CEP350, KIF3B, HYLS1:** [Low] Ciliogenesis factors dependent on vesicle trafficking machinery; peripheral support to the membrane-trafficking function.

-- genes supporting the name:
TAF6, POLR2J, TFCP2, ATG5, ATG13, ATG14, WIPI2, PI4KB, PIP5K1A, ZFYVE1, VPS33A, VPS33B, VPS39, YKT6, UBE2H, FBXW11, USP19, SIRT1, LAMTOR4, RRAGC, HIF1AN, PRKAG1, RETREG3, PEX1, DERL2, TTC26, BBS4, CEP350, KIF3B, HYLS1

### ZNF446 HepG2

#### Process: Transcription factor–directed ciliogenesis with peroxisome import and genome maintenance (0.86)

The enriched categories coalesce around RNAPII-centered transcription and RNA processing together with organelle programs for the primary cilium and peroxisome, and a parallel genome-stability module. Together, these define a coherent network where sequence-specific transcription (dominated by C2H2 ZNFs) drives expression of ciliary/peroxisomal machinery while DNA repair pathways protect cells during organelle remodeling.

##### 1. Transcriptional control and RNA maturation

C2H2 zinc-finger transcription factors and “DNA-binding transcription factor activity, RNA polymerase II-specific” are strongly enriched, along with the “Generic Transcription Pathway/RNA Polymerase II Transcription.” Spliceosomal/RNA processing terms tie transcription to mature gene output, positioning *RNAPII* regulation as the network’s organizing layer. The gene list contains numerous ZNFs and core transcription/RNA-processing factors that populate these categories, supporting this layer as a major gene sink.

##### 2. Ciliogenesis and membrane/organelle logistics

“Cilium assembly/organization,” “Intraflagellar transport,” and “Anchoring of the basal body to the plasma membrane” indicate a robust ciliogenesis module. The set encodes multiple IFT/transition-zone and ciliopathy scaffold components (e.g., *IFT122/88/52/172/43*, *TCTN1/2*, *CEP164/131/89*, *B9D1/B9D2*, *MKS1*, *FUZ*, *CLUAP1*), consistent with axoneme construction, cargo movement, and basal-body docking. Vesicle-trafficking/ESCRT entries (*CHMP1A/6*, *EPN1*, *SCAMPs*) align with “protein transmembrane import into intracellular organelle,” supporting ciliary membrane delivery. [1][2][3]

---

[1] Garcia-Gonzalo, Francesc R, Corbit, Kevin C, Sirerol-Piquer, María Salomé, Ramaswami, Gokul, Otto, Edgar A, Noriega, Thomas R, Seol, Allen D, Robinson, Jon F, Bennett, Christopher L, Josifova, Dragana J, García-Verdugo, José Manuel, Katsanis, Nicholas, Hildebrandt, Friedhelm, Reiter, Jeremy F. ""A transition zone complex regulates mammalian

ciliogenesis and ciliary membrane composition."" Nature genetics, 2011, pp. 776-84, doi: https://doi.org/10.1038/ng.891

[2] Tanos, Barbara E, Yang, Hui-Ju, Soni, Rajesh, Wang, Won-Jing, Macaluso, Frank P, Asara, John M, Tsou, Meng-Fu Bryan. ""Centriole distal appendages promote membrane docking, leading to cilia initiation."" Genes & development, 2013, pp. 163-8, doi: https://doi.org/10.1101/gad.207043.112

[3] Roberson, Elle C, Dowdle, William E, Ozanturk, Aysegul, Garcia-Gonzalo, Francesc R, Li, Chunmei, Halbritter, Jan, Elkhartoufi, Nadia, Porath, Jonathan D, Cope, Heidi, Ashley-Koch, Allison, Gregory, Simon, Thomas, Sophie, Sayer, John A, Saunier, Sophie, Otto, Edgar A, Katsanis, Nicholas, Davis, Erica E, Attié-Bitach, Tania, Hildebrandt, Friedhelm, Leroux, Michel R, Reiter, Jeremy F. ""TMEM231, mutated in orofaciodigital and Meckel syndromes, organizes the ciliary transition zone."" The Journal of cell biology, 2015, pp. 129-42, doi: https://doi.org/10.1083/jcb.201411087

#### 3. Peroxisome matrix import and organization

“Protein import into peroxisome matrix” and “peroxisome organization” are enriched. The gene set encodes *PEX5/10/12/14/16* and *ABCD4*, mapping onto PTS-dependent receptor–docking–translocation steps and peroxisome maintenance, indicating a coordinated peroxisomal biogenesis/turnover arm within the same transcriptional program. [1][2][3]

---

[1] Feng, Peiqiang, Wu, Xudong, Erramilli, Satchal K, Paulo, Joao A, Knejski, Pawel, Gygi, Steven P, Kossiakoff, Anthony A, Rapoport, Tom A. ""A peroxisomal ubiquitin ligase complex forms a retrotranslocation channel."" Nature, 2022, pp. 374-380, doi: https://doi.org/10.1038/s41586-022-04903-x

[2] Feng, Peiqiang, Skowyra, Michael L, Rapoport, Tom A. ""Structure and function of the peroxisomal ubiquitin ligase complex."" Biochemical Society transactions, 2022, pp. 1921-1930, doi: https://doi.org/10.1042/BST20221393

[3] Wu, Jianguo, Lv, Shupei, Xu, Peng, Yue, Suyu, Zhuang, Min. ""PEX14 condensates recruit receptor and cargo pairs for peroxisomal protein import."" Nature structural & molecular biology, 2025, pp. 1644-1656, doi: https://doi.org/10.1038/s41594-025-01601-w

#### 4. Genome maintenance and telomere control

Enrichment for “DNA Repair,” “Global Genome Nucleotide Excision Repair (GG-NER),” and “Resolution of Abasic Sites,” with “Extension of Telomeres,” points to active genome surveillance alongside organelle assembly. The gene set includes base-excision/SSBR and NER/telomere components (*XRCC1*, *OGG1*, *POLL*/*POLM*, *LIG1*, *PNKP*, *ERCC1*, *XPC*, *ACD*/*WRAP53*, *RTEL1*), supporting replication–transcription conflict buffering during ciliogenesis. [1][2][3]

---

[1] Mashayekhi, Fatemeh, Zeinali, Elham, Ganje, Cassandra, Fanta, Mesfin, Li, Lei, Godbout, Roseline, Weinfeld, Michael, Ismail, Ismail Hassan. ""CDK-dependent phosphorylation regulates PNKP function in DNA replication."" The Journal of biological chemistry, 2024, pp. 107880, doi: https://doi.org/10.1016/j.jbc.2024.107880

[2] Kumar, Namrata, Theil, Arjan F, Roginskaya, Vera, Ali, Yasmin, Calderon, Michael, Watkins, Simon C, Barnes, Ryan P, Opresko, Patricia L, Pines, Alex, Lans, Hannes, Vermeulen, Wim, Van Houten, Bennett. ""Global and transcription-coupled repair of 8-oxoG is initiated by nucleotide excision repair proteins."" Nature communications, 2022, pp. 974, doi: https://doi.org/10.1038/s41467-022-28642-9

[3] Breslin, Claire, Mani, Rajam S, Fanta, Mesfin, Hoch, Nicolas, Weinfeld, Michael, Caldecott, Keith W. ""The Rev1 interacting region (RIR) motif in the scaffold protein XRCC1 mediates a low-affinity interaction with polynucleotide kinase/phosphatase (PNKP) during DNA single-strand break repair."" The Journal of biological chemistry, 2017, pp. 16024-16031, doi: https://doi.org/10.1074/jbc.M117.806638

#### 5. Metabolic and cofactor support

“Cytosolic iron–sulfur cluster assembly” is enriched, providing essential cofactors for DNA repair/replication enzymes. Matching CIA/NUBP machinery and mitochondrial assembly genes (*CIAO3*, *NUBP2*, *FOXRED1*, *COQ* genes) appear in the list, linking redox/energy status to repair and organelle programs. [1][2]

---

[1] Lemire, Bernard D. ""Evolution of FOXRED1, an FAD-dependent oxidoreductase necessary for NADH:ubiquinone oxidoreductase (Complex I) assembly."" Biochimica et biophysica acta, 2015, pp. 451-457, doi: https://doi.org/10.1016/j.bbabio.2015.01.014

[2] Maione, Vincenzo, Grifagni, Deborah, Torricella, Francesco, Cantini, Francesca, Banci, Lucia. ""CIAO3 protein forms a stable ternary complex with two key players of the human cytosolic iron-sulfur cluster assembly machinery."" Journal of biological inorganic chemistry : JBIC : a publication of the Society of Biological Inorganic Chemistry, 2020, pp. 501-508, doi: https://doi.org/10.1007/s00775-020-01778-z

#### 6. Growth and sensory context

AKT-axis terms ("AKT phosphorylates targets in the cytosol," "Constitutive AKT1 signaling") situate ciliogenesis and repair under nutrient/growth control. The presence of *AKT2*/*AKT1S1*/*MLST8* in the set supports this gatekeeping role. "Sensory perception/olfactory signaling" further aligns with a cilium-based receptor landscape; the gene list contains multiple ciliary transport/docking factors required for such signaling.

Network integration

The dominant architecture is a TF-heavy RNAPII program that builds and services primary cilia and peroxisomes, while genome-repair/telomere modules safeguard DNA during these organelle processes; Fe–S biogenesis and AKT/mTOR nodes supply cofactors and cue growth-state gating. This integrated motif is repeatedly captured by the enriched terms and represented densely within the gene set.

Confidence

A large fraction of genes map to RNAPII/ZNF regulation, ciliary/peroxisomal modules, and DNA repair/telomeres, with multiple independent term enrichments corroborating the same axes. Hence the high but not maximal confidence (0.86).

-- Summary --

The text identifies genes supporting the name *"Transcription factor–directed ciliogenesis with* *peroxisome import and genome maintenance."* Genes directly linked to transcriptional regulation (e.g., ZNF263, USF2, E4F1), ciliogenesis (IFT122, IFT88, FUZ, MKS1, CEP164, CLUAP1), peroxisome import (PEX5, PEX10, PEX12, PEX14, PEX16), and genome maintenance (XRCC1, LIG1, OGG1, POLM, XPC, ERCC1, RTEL1) provide strong support.

-- Explanation --

**ZNF263:** [High] Described as a C2H2 zinc-finger transcription factor regulating RNAPII-dependent gene expression, supporting the transcriptional control aspect of the process.

**USF2:** [High] Identified as a transcription factor linked to the RNAPII pathway, contributing to the TF-directed regulation of ciliogenesis and organelle biogenesis.

**E4F1:** [High] Cited as a transcriptional regulator that coordinates genome stability with organelle biogenesis, directly reinforcing the integrated TF-driven nature of the process.

**IFT122:** [High] Mentioned among key intraflagellar transport components essential for cilium assembly and axoneme construction.

**IFT88:** [High] Listed as a central IFT complex B component supporting ciliogenesis and vesicle trafficking to the ciliary axoneme.

**FUZ:** [High] Defined as a ciliopathy scaffold protein necessary for basal body docking and ciliary organization.

**MKS1:** [High] Included in transition-zone components controlling ciliary membrane organization, directly supporting ciliogenesis.

**CEP164:** [High] Reported as a distal appendage and basal-body anchoring protein involved in ciliogenesis.

**CLUAP1:** [High] Recognized as an IFT-associated factor involved in ciliary cargo transport, reinforcing the cilium assembly network.

**PEX5:** [High] Functions as a peroxisomal targeting signal receptor, mediating peroxisome matrix import.

**PEX10 / PEX12 / PEX14 / PEX16:** [High] Each involved in docking, translocation, and maintenance of peroxisome matrix proteins, confirming the peroxisome import module.

**XRCC1:** [High] Identified as a scaffold protein for base-excision repair and single-strand break repair, linking the set to genome maintenance.

**LIG1:** [High] DNA ligase essential for DNA replication and repair, participating in genome-stability maintenance.

**OGG1:** [High] 8-oxoguanine glycosylase involved in base-excision repair and oxidative DNA lesion removal, supporting genome maintenance.

**POLM:** [Medium] Polymerase involved in DNA end-joining and base-excision repair, participating in repair of abasic sites.

**XPC / ERCC1:** [High] Core nucleotide excision repair genes that function in global-genome NER, strengthening the DNA-repair component.

**RTEL1:** [High] DNA helicase responsible for telomere maintenance and replication stress resolution.

**CIAO3 / NUBP2:** [Medium] Support “cytosolic iron–sulfur cluster assembly,” linking metabolic cofactors to DNA-repair enzyme maturation.

**AKT2 / AKT1S1 / MLST8:** [Medium] Included under “AKT/mTOR signaling” controlling growth-linked ciliogenesis and genome-repair timing.

-- genes supporting the name:
ZNF263, USF2, E4F1, IFT122, IFT88, FUZ, MKS1, CEP164, CLUAP1, PEX5, PEX10, PEX12, PEX14, PEX16, XRCC1, LIG1, OGG1, POLM, XPC, ERCC1, RTEL1, CIAO3, NUBP2, AKT2, AKT1S1, MLST8

### ZNF557 HepG2

#### Process: Transcription-coupled RNA processing and RNP biogenesis (0.90)

The system is dominated by transcriptional regulators (especially C2H2 zinc-finger TFs) partnered with extensive RNA metabolism machinery. Enrichment spans RNAP I/II/III transcription, co-transcriptional capping, splicing, 3′-end formation, RNA modification, and assembly/turnover of ribonucleoprotein (RNP) complexes, extending into ribosome/tRNA/rRNA biogenesis. Secondary modules (DNA repair, telomere maintenance, sensory/GPCR signaling) likely reflect contextual coupling rather than the primary function.

#### 1. Core transcriptional control

PANTHER terms highlight “regulation of transcription by RNA polymerase II,” “Generic Transcription Pathway,” and “RNA Polymerase II Transcription,” indicating a central RNAPII axis. Gene set content includes basal factors and polymerase subunits (e.g., *TBP*/*GTF2B*/*GTF2E1*; *POLR2D* alongside *POLR1B*/*POLR3A/C/F*), positioning coordinated initiation across polymerases I–III. [1][2]

---

[1] Santana, Juan F, Collins, Geoffrey S, Parida, Mrutyunjaya, Luse, Donal S, Price, David H. ""Differential dependencies of human RNA polymerase II promoters on TBP, TAF1, TFIIB and XPB."" Nucleic acids research, 2022, pp. 9127-9148, doi: https://doi.org/10.1093/nar/gkac678

[2] Maeta, Masanari, Kataoka, Miku, Nishiya, Yusuke, Ogino, Kazutoyo, Kashima, Makoto, Hirata, Hiromi. ""RNA polymerase II subunit D is essential for zebrafish development."" Scientific reports, 2020, pp. 13213, doi: https://doi.org/10.1038/s41598-020-70110-1

#### 2. Co-transcriptional RNA processing

Strong enrichment for “RNA processing,” “mRNA processing,” “RNA splicing (via spliceosome),” and “RNA 3′-end processing” maps onto spliceosomal (*PRPF39*/*ISY1*/*AAR2*), cleavage/polyadenylation (*CSTF1*/*CSTF2*/*CSTF2T*/*PCF11*), and capping modules (*TGS1*/*MEPCE*). These modules channel nascent RNAs to mature transcripts in tight linkage with transcription elongation/termination. [1][2][3]

---

[1] Kamieniarz-Gdula, Kinga, Gdula, Michal R, Panser, Karin, Nojima, Takayuki, Monks, Joan, Wiśniewski, Jacek R, Riepsaame, Joey, Brockdorff, Neil, Pauli, Andrea, Proudfoot, Nick J. ""Selective Roles of Vertebrate PCF11 in Premature and Full-Length Transcript Termination."" Molecular cell, 2019, pp. 158-172.e9, doi:

https://doi.org/10.1016/j.molcel.2019.01.027

[2] Yang, Yuan, Eichhorn, Catherine D, Wang, Yaqiang, Cascio, Duilio, Feigon, Juli. ""Structural basis of 7SK RNA 5'-γ-phosphate methylation and retention by MePCE."" Nature chemical biology, 2019, pp. 132-140, doi: https://doi.org/10.1038/s41589-018-0188-z

[3] Bejjani, Fabienne, Ségéral, Emmanuel, Mosca, Kevin, Lecourieux, Adriana, Bakail, May, Hamoudi, Meriem, Emiliani, Stéphane. ""Overlapping and distinct functions of SPT6, PNUTS, and PCF11 in regulating transcription termination."" Nucleic acids research, 2025, pp. , doi: https://doi.org/10.1093/nar/gkaf179

#### 3. RNA modification and quality control

Enrichment for "RNA methylation," "tRNA modification," and broader "RNA modification" is mirrored by *TRMT1*/*TRMO*/*TRMT10B*/*METTL14*/*NSUN3*/*BCDIN3D* and tRNA wybutosine pathway enzymes (*TYW1/5*). Exosome/decay factors (*EXOSC3*/*EXOSC6*/*DIS3*/*ISG20L2*) supply surveillance and turnover, ensuring transcriptome fidelity. [1][2]

---

[1] Qi, Ya-Nan, Liu, Zhu, Hong, Lian-Lian, Li, Pei, Ling, Zhi-Qiang. ""Methyltransferase-like proteins in cancer biology and potential therapeutic targeting."" Journal of hematology & oncology, 2023, pp. 89, doi: https://doi.org/10.1186/s13045-023-01477-7

[2] Shi, Bin, Liu, Wei-Wei, Yang, Ke, Jiang, Guan-Min, Wang, Hao. ""The role, mechanism, and application of RNA methyltransferase METTL14 in gastrointestinal cancer."" Molecular cancer, 2022, pp. 163, doi: https://doi.org/10.1186/s12943-022-01634-5

#### 4. RNP assembly and ribosome/tRNA/rRNA biogenesis

"Ribonucleoprotein complex biogenesis," "rRNA processing," "ribosome biogenesis," and "tRNA metabolic process" are prominent. The set contains numerous SSU processome/UTP factors (*UTP14A*/*UTP23*/*MAK16*/*RRP36*), rRNA/tRNA enzymes (*RRP8*/*TRUB2*/*PUS3*), and RNAPI initiation factor *RRN3*, tying nucleolar assembly to the broader gene-expression pipeline. [1][2][3]

---

[1] Choudhury, Priyanka, Hackert, Philipp, Memet, Indira, Sloan, Katherine E, Bohnsack,

Markus T. ""The human RNA helicase DHX37 is required for release of the U3 snoRNP from pre-ribosomal particles."" RNA biology, 2019, pp. 54-68, doi: https://doi.org/10.1080/15476286.2018.1556149

[2] Zang, Yue, Ran, Xia, Yuan, Jie, Wu, Hao, Wang, Youya, Li, He, Teng, Huajing, Sun, Zhongsheng. ""Genomic hallmarks and therapeutic targets of ribosome biogenesis in cancer."" Briefings in bioinformatics, 2024, pp. , doi: https://doi.org/10.1093/bib/bbae023

[3] Nøstvik, Miriam, Kateta, Sarah M, Schönewolf-Greulich, Bitten, Afenjar, Alexandra, Barth, Magalie, Boschann, Felix, Doummar, Diane, Haack, Tobias B, Keren, Boris, Livshits, Ludmila A, Mei, Davide, Park, Joohyun, Pisano, Tiziana, Prouteau, Clement, Umair, Muhammad, Waqas, Ahmed, Ziegler, Alban, Guerrini, Renzo, Møller, Rikke S, Tümer, Zeynep. ""Clinical and molecular delineation of PUS3-associated neurodevelopmental disorders."" Clinical genetics, 2021, pp. 628-633, doi: https://doi.org/10.1111/cge.14051

#### 5. Broad TF superstructure

Molecular-function enrichments ("DNA-binding transcription factor activity, RNAPII-specific," "sequence-specific DNA binding") align with the dense C2H2 ZNF cohort (e.g., *ZNF263*/*ZNF646*/*ZNF507*/*ZNF777*), which likely pattern gene-specific outputs that feed into or regulate the RNA-handling machinery. [1][2][3]

---

[1] Weiss, Ryan J, Spahn, Philipp N, Toledo, Alejandro Gómez, Chiang, Austin W T, Kellman, Benjamin P, Li, Jing, Benner, Christopher, Glass, Christopher K, Gordts, Philip L S M, Lewis, Nathan E, Esko, Jeffrey D. ""ZNF263 is a transcriptional regulator of heparin and heparan sulfate biosynthesis."" Proceedings of the National Academy of Sciences of the United States of America, 2020, pp. 9311-9317, doi: https://doi.org/10.1073/pnas.1920880117

[2] Yu, Zhibin, Feng, Jianbo, Wang, Wei, Deng, Zhiyong, Zhang, Yan, Xiao, Lan, Wang, Zeyou, Liu, Changhong, Liu, Qing, Chen, Shuai, Wu, Minghua. ""The EGFR-ZNF263 signaling axis silences SIX3 in glioblastoma epigenetically."" Oncogene, 2020, pp. 3163-3178, doi: https://doi.org/10.1038/s41388-020-1206-7

[3] Yuki, Ryuzaburo, Aoyama, Kazumasa, Kubota, Sho, Yamaguchi, Noritaka, Kubota, Shoichi, Hasegawa, Hitomi, Morii, Mariko, Huang, Xiayu, Liu, Kang, Williams, Roy, Fukuda, Michiko N, Yamaguchi, Naoto. ""Overexpression of zinc-finger protein 777 (ZNF777) inhibits proliferation at low cell density through down-regulation of FAM129A."" Journal of cellular

biochemistry, 2015, pp. 954-68, doi: https://doi.org/10.1002/jcb.25046

#### 6. Coupled auxiliary programs (contextual, not dominant)

DNA repair/maintenance terms (e.g., “NER,” “telomere organization”) correspond to *ERCC3*/*ERCC4*/*XPC*/*TERF2*/*HELQ*/*RAD51D*, providing genome-stability support during high transcriptional throughput. Signaling/GPCR and sensory annotations (olfactory signaling) and cytoskeletal/adhesion categories likely represent cellular context rather than the primary functional thrust.

Network integration

Together, polymerase initiation/elongation, co-transcriptional processing, RNA modification, and RNP assembly form a single, interlocked production line from DNA-templated transcription to mature RNPs and ribosomes. The breadth and depth of enrichment across these steps, plus the high representation of matching genes, justify the process designation and high confidence.

Representative genes anchoring each module (non-exhaustive):

• Transcription: TBP, GTF2B, GTF2E1, MED17/19/26/29, POLR2D/POLR1B/POLR3A/C/F, RRN3.

• Splicing/3′-end: PRPF39, ISY1, AAR2, CSTF1/2/2T, PCF11, CWF19L1.

• Capping/modification: TGS1, MEPCE, TRMT1/TRMO/TRMT10B, METTL14, NSUN3, TYW1/5.

• RNP/rRNA/tRNA biogenesis: UTP14A/UTP23/MAK16/RRP36, RRP8, TRUB2, PUS3, RRN3.

• Surveillance/decay: EXOSC3/EXOSC6, DIS3, ISG20L2.

• TF scaffold: Numerous ZNFs (e.g., ZNF263, ZNF646, ZNF507, ZNF777) with RNAPII-specific DNA-binding activity enrichment.

Why this name and score: The densest, most coherent enrichment spans transcription → RNA processing → RNP assembly, with many genes mapping to these steps; ancillary pathways are comparatively sparse. Thus, “Transcription-coupled RNA processing and RNP biogenesis” best captures the predominant activity with high—but not absolute—confidence (0.90).

-- Summary --
The analysis text identifies a highly integrated system centered on transcription-coupled RNA processing and ribonucleoprotein (RNP) biogenesis. Definite assertions supporting this name include genes directly linked to RNA polymerase I–III transcription, splicing, capping, RNA modification, and RNP assembly. Multiple zinc-finger transcription factors (ZNFs) also reinforce the transcriptional control aspect of the system.

-- Explanation --
**TBP:** [High] Described as a basal transcription factor essential for RNA polymerase II initiation, directly supporting the transcription component of the process.

**GTF2B:** [High] Identified as part of the general transcription machinery required for polymerase II initiation, supporting transcription-coupled regulation.

**POLR2D:** [High] Mentioned as a subunit of RNA polymerase II, directly responsible for transcription of protein-coding genes.

**PRPF39:** [High] Cited as part of the spliceosomal machinery linking transcription elongation to RNA splicing, confirming co-transcriptional RNA processing.

**ISY1:** [High] Listed as a spliceosomal component mediating RNA splicing during co-transcriptional processing.

**AAR2:** [High] Explicitly part of spliceosome assembly, connecting transcription to mRNA maturation.

**CSTF1 / CSTF2 / CSTF2T / PCF11:** [High] Each identified as a 3′-end cleavage/polyadenylation factor participating in mRNA processing downstream of transcription.

**TGS1 / MEPCE:** [High] Stated to perform RNA capping modifications during co-transcriptional processing.

**TRMT1 / TRMO / TRMT10B / METTL14 / NSUN3 / BCDIN3D:** [High] Described as methyltransferases or RNA modification enzymes that ensure transcript stability and function within the RNA maturation pipeline.

**TYW1 / TYW5:** [High] Identified as tRNA modification enzymes supporting tRNA metabolic processes within RNP biogenesis.

**EXOSC3 / EXOSC6 / DIS3 / ISG20L2:** [High] Cited as exosome components responsible for RNA surveillance and degradation, ensuring RNA quality control in the transcription-coupled pathway.

**UTP14A / UTP23 / MAK16 / RRP36:** [High] Defined as small subunit (SSU) processome factors involved in rRNA processing and ribosome biogenesis.

**RRP8 / TRUB2 / PUS3 / RRN3:** [High] Mentioned as nucleolar enzymes for rRNA/tRNA processing and RNAPI initiation, linking transcription to ribosome assembly.

**ZNF263 / ZNF646 / ZNF507 / ZNF777:** [Medium] Cited as C2H2 zinc-finger transcription factors that regulate RNA polymerase II–specific transcription and control expression of RNA metabolism genes.

**ERCC3 / ERCC4 / XPC / TERF2 / HELQ / RAD51D:** [Medium] Identified as DNA repair factors supporting genome stability during high transcriptional activity, a secondary but related function.

-- genes supporting the name:
TBP, GTF2B, POLR2D, PRPF39, ISY1, AAR2, CSTF1, CSTF2, CSTF2T, PCF11, TGS1, MEPCE, TRMT1, TRMO, TRMT10B, METTL14, NSUN3, BCDIN3D, TYW1, TYW5, EXOSC3, EXOSC6, DIS3, ISG20L2, UTP14A, UTP23, MAK16, RRP36, RRP8, TRUB2, PUS3, RRN3, ZNF263, ZNF646, ZNF507, ZNF777, ERCC3, ERCC4, XPC, TERF2, HELQ, RAD51D
